# Tc17-driven antibody-independent mucosal immunity is critical for protection against extracellular bacterial pneumonia

**DOI:** 10.64898/2026.08.26.747429

**Authors:** Yuheng Liu, Jingjing Zhang, Zhifu Chen, Rui Liao, Chao Li, Qin Xiao, Shan Guan

## Abstract

*Klebsiella pneumoniae* (Kp) is a WHO high-priority pathogen for vaccine development, yet previous efforts failed largely because key protective immune mechanisms remain unclear. Here we show that protective immunity conferred by mucosal mRNA vaccines (but not parenteral) require neither serum IgG nor airway secretory IgA, but instead depends on a previously unrecognized lung-resident CD8⁺IL-17⁺ T-cells (Tc17) that rapidly recruits neutrophils/macrophages to eliminate bacteria. To therapeutically harness this paradigm, we developed INSPIRE, a machine learning-engineered exosome platform incorporating donor-screened, miRNA-bioactive backbones (miR-21-mediated airway barrier penetration and miR-155-associated dendritic-cell activation through SOCS1/Inpp5d axis) and computationally designed peptides that boosts 11.6-fold mRNA encapsulation and 3-fold dendritic-cell cross-presentation. Intranasal INSPIRE-mRNA vaccination confers near-complete protection against clinically relevant Kp strains while intramuscular counterparts fail (below ∼30% survival). Leveraging pIgR^-^/^-^ and IL-17^-^/^-^ mice coupled with T-cell depletions, we demonstrate the protection is Tc17-dependent. This work overturns the antibody-centric dogma and redefines a non-canonical Tc17-correlate for extracellular bacterial pneumonia.

## Introduction

The escalating crisis of antimicrobial resistance has placed *Klebsiella pneumoniae* (Kp) at the forefront of global health threats. ^1–4^ Recognized by the World Health Organization (WHO) as a high priority pathogen for vaccine development, Kp exemplifies the failure of classical vaccinology against extracellular bacterial pneumonia. ^3–5^ Decades of efforts focusing on systemic antibodies have yielded no licensed vaccine, largely because they overlook a cardinal feature of pulmonary Kp infection: the pathogen’s rapid colonization in the airway mucosa demands an immediate, tissue-resident defence that systemic immunity alone cannot provide. ^4,6–10^ Consequently, there is a pressing need to reimagine mucosal vaccination-not merely as a route of administration, but as a strategy to actively engineer protective immunological niches directly within the site of infection, i.e. respiratory tract. ^11–16^

While mRNA technology offers programmability for multi-antigen coverage and rapid development, ^17–20^ its application to Kp vaccine is still uncharted and most probably constrained by two intertwined challenges. ^17,18,21–23^ First, the airway presents formidable physical and immunological barriers: mucus, tight junctions, and tolerogenic dendritic cells (DC) limit both nanoparticle penetration and the generation of effector immunologic cells. ^13,15,16,23^ Second, and more fundamentally, the protective correlates for extracellular bacterial pneumonia remain incompletely defined. Although secretory IgA (sIgA) is often assumed to be the key mediator of mucosal defence, we and others have found that sIgA is neither sufficient nor strictly required for Kp clearance. ^11,24–26^ This paradox suggests that an effective Kp vaccine must elicit a different class of immunity—one that is lung-localized, durably inducible, and functionally distinct from systemic humoral responses. Yet, because Kp is an extracellular pathogen, the potential contribution of CD8⁺ T cells has been largely ignored. Whether a vaccine can purposely induce CD8⁺ T cells that produce IL-17 (Tc17 cells) with tissue-resident features, and whether such cells can control an extracellular bacterium, represents an unexplored paradigm in mucosal immunology. ^27–33^

Here we resolve these challenges by developing INSPIRE (**I**ntranasal **N**anoengineered **S**mart **P**latform for **I**mmune **R**eprogramming via **E**xosomes)-a machine learning (ML)-programmed, bioactive exosome platform that fundamentally potentializes mucosal mRNA vaccination. Rather than using inert delivery vehicles, we first performed a systematic functional screen of donor-cell-derived exosomes and identified mature dendritic cell-derived exosomes (mBMDC-Exos) as a privileged backbone. ^34–38^ Mechanistically, this backbone integrates two natural miRNA programs: miR-21-mediated, reversible epithelial junctional modulation that enhances transepithelial passage through PTEN-, SMAD7- and PDCD4-related pathways, and miR-155-mediated relief of SOCS1/Inpp5d inhibitory circuits to license DC activation. ^39–45^ On the other hand, native exosomes lack sufficient mRNA encapsulation and DC-directed transfection efficiency. To overcome this, we incorporated an ML-guided peptide engineering strategy that converted the bioactive mBMDC-Exos backbone into INSPIRE, elevating mRNA encapsulation from ∼5% to 58% and dramatically enhancing DC cross-presentation. ^46,47^

Applying the INSPIRE to an intranasal mRNA vaccine encoding conserved Kp antigens elicits durable acquired immunities featured by a dominant mucosal immune response together with concurrent humoral and cellular immune activation. This mucosal Kp-mRNA vaccination turn out to confer durable, cross-strain mucosal protection against highly virulent, drug-resistant Kp strains, while conventional parenteral immunization via the intramuscular route fails to do so, only yielding survival rates below ∼30%. Mechanistic investigations reveal an unexpected observation that protection does not depend on the high titers of airway sIgA and serum IgG induced, but instead on a previously unappreciated CD8⁺IL-17⁺ lung-resident T-cell axis. Using pIgR^-^/^-^ and IL-17^-^/^-^ knockout mice, we demonstrate that airway antibodies are dispensable, whereas IL-17 is essential. Notably, CD8⁺ T-cell depletion severely abrogated protection, revealing the Tc17 subsets-long considered irrelevant for extracellular bacteria—are critical effectors. Transcriptomic and functional analyses show that INSPIRE elicits a lung-resident Tc17 program characterized by coordinated upregulation of *Il23r*, *Cxcr3*-*Cxcl9*/*Cxcl10* axis, *Cd69*, and *Itgae* (*Cd103*), leading to rapid neutrophil/macrophage recruitment upon challenge, near-complete bacterial clearance, and marvelous cross-strain protection.

Taken together, this work uncovers a functional CD8⁺IL-17⁺ airway-resident T-cell response as a dominant, non-redundant mediator of protection against Kp, overturning the antibody-centric paradigm. Leveraging ML-engineered exosomes that actively shape the pulmonary immune niche rather than passively delivering antigen, we establish a biologically inspired design principle of combining reversible epithelial barrier modulation, DC-targeted mRNA delivery, and intrinsic miRNA-based adjuvanticity in a single delivery architecture—INSPIRE. Our findings provide both T cell activating design principle for mucosal mRNA vaccines and fundamentally redefine the protective correlates for extracellular drug-resistant bacteria by uncovering the key function of non-canonical Tc17 immunity.

## Results and Discussion

### Functional exosome screening and miRNA profiling identify mature BMDC-derived exosomes, enriched in miR-21 and miR-155, as a bioactive backbone

To identify an exosome source suitable for airway mRNA vaccination, we established a transwell-based epithelial–immune screening system (**Fig. 1a**). The assay measured transepithelial transport, bone marrow-derived dendritic cell (BMDC) activation and epithelial barrier integrity in parallel. We compared exosomes (Exos) from eight donor-cell sources spanning macrophage, epithelial, stromal and dendritic-cell lineages, including Raw264.7 M0 macrophages, Raw264.7 M2 macrophages, BEAS-2B cells, A549 cells, HEK-293T cells, umbilical-cord mesenchymal stem cells (uMSCs), immature BMDC-derived Exos (I-BMDC-Exos) and mature BMDC-derived Exos (M-BMDC-Exos) (**Fig. 1b**). In the three-parameter evaluation matrix, the tested exosome sources showed distinct performance profiles. Among them, M-BMDC-Exos showed the most favourable combined profile, combining higher transepithelial transport and BMDC activation than most other exosome sources with the ability to transiently and reversibly modulate epithelial barrier integrity. M-BMDC-Exos induced an early decrease in TEER, consistent with temporary loosening of epithelial junctions, followed by recovery towards baseline, suggesting reversible epithelial junctional remodelling rather than persistent barrier damage (**Fig. 1b**). We therefore selected M-BMDC-Exos as a candidate bioactive backbone for subsequent mechanistic analysis and engineering.

**Fig. 1.**
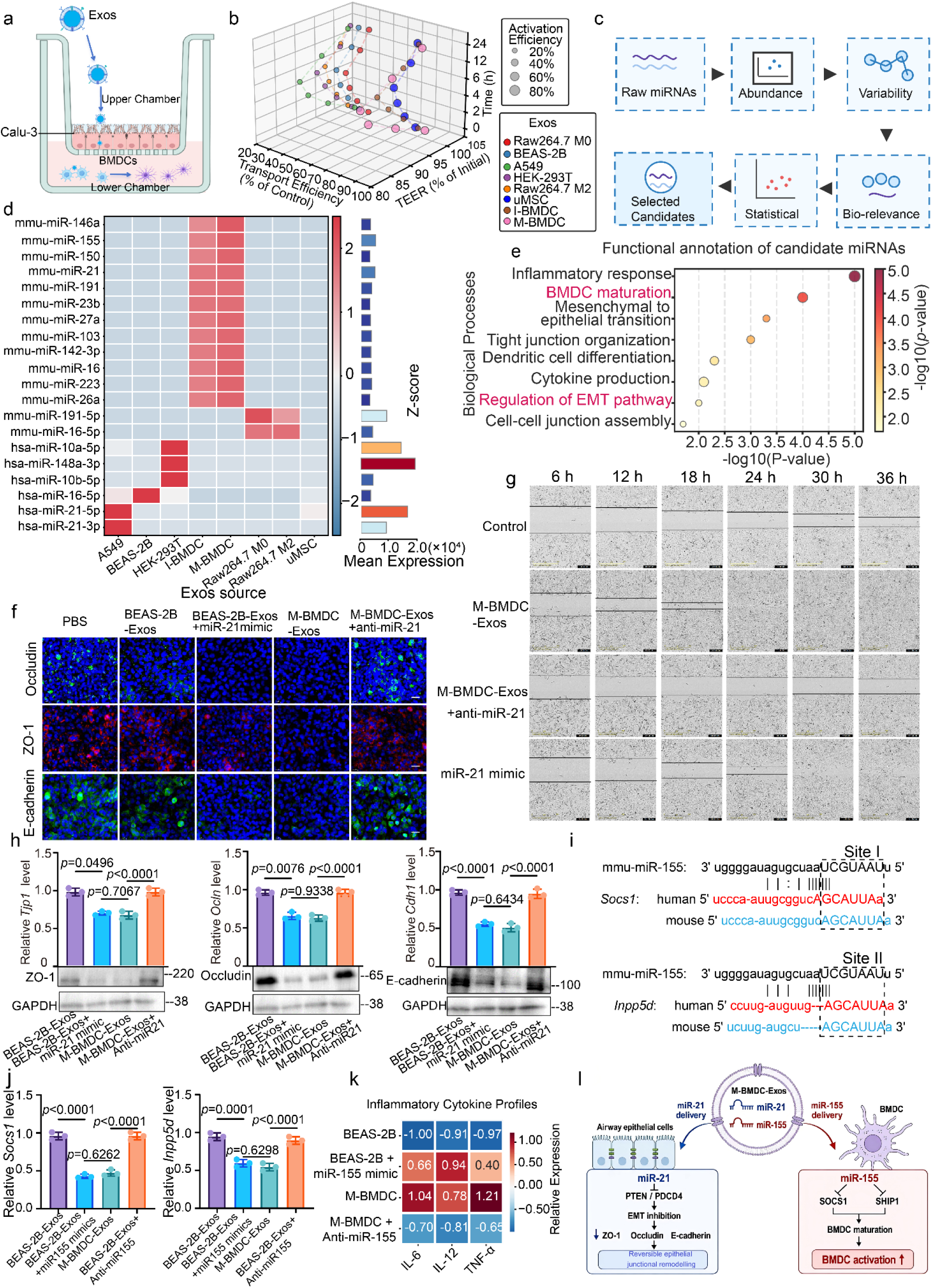
Functional exosome screening and miRNA profiling identify mature BMDC-derived exosomes as a bioactive backbone. **a**, Schematic of the transwell-based epithelial–immune screening system used to evaluate donor-cell-derived exosomes. Calu-3 epithelial cells were cultured on porous transwell inserts to form an epithelial barrier, and bone marrow-derived dendritic cells (BMDCs) were placed in the lower chamber. Exosomes were added to the apical chamber, and transepithelial transport, epithelial barrier integrity and BMDC activation were assessed. **b**, Three-dimensional screening plot integrating exosome transport efficiency, transepithelial electrical resistance (TEER) and incubation time for exosomes derived from eight donor-cell sources. Bubble size indicates BMDC activation efficiency, and colors indicate exosome source. I-BMDC-Exos and M-BMDC-Exos denote exosomes derived from immature and mature BMDCs, respectively. **c**, Stepwise microRNA (miRNA)-screening workflow applied to public exosomal miRNA-sequencing data from the Gene Expression Omnibus (GEO) database GSE190854. Candidate miRNAs were filtered by abundance, donor-source variability, biological relevance and statistical prioritization, followed by selection of candidates for downstream validation. **d**, Heatmap of the top 20 candidate miRNAs across eight donor-cell-derived exosome groups after abundance and variability filtering (RPM > 100 and CV > 50%). Expression values were z-score normalized across groups, and the bar plot on the right shows mean expression levels across groups. **e**, Functional annotation of candidate miRNAs after screening. Biological processes related to inflammatory response, BMDC maturation, epithelial transition, tight-junction organization, dendritic-cell differentiation, cytokine production, regulation of the epithelial-mesenchymal transition pathway and cell–cell junction assembly are shown. **f–h,** Epithelial junctional modulation was analysed 12 h after the indicated treatments. **f**, Representative immunofluorescence images of epithelial junction-associated proteins. human bronchial epithelial cells (16HBE) cells were stained for occludin, zonula occludens-1 (ZO-1) and E-cadherin; nuclei were counterstained with 4′,6-diamidino-2-phenylindole (DAPI). Scale bars, 50 μm. **g**, Representative scratch-wound images of 16HBE epithelial monolayers treated with control medium, M-BMDC-Exos, miR-21-inhibited M-BMDC-Exos or a miR-21 mimic. Scale bars, 600 μm. **h**, Relative transcript levels of *Tjp1*, *Ocln* and *Cdh1*, together with representative western blots of ZO-1, occludin and E-cadherin. Glyceraldehyde-3-phosphate dehydrogenase (GAPDH) was used as the loading control for western blotting. **i**, Sequence alignment showing predicted miR-155-binding sites in the 3′ untranslated regions of *Socs1* and *Inpp5d* across human and mouse sequences. Inpp5d encodes SH2 domain-containing inositol 5′-phosphatase 1 (SHIP1). **j**, Relative expression levels of *Socs1* and *Inpp5d* transcripts after the indicated exosome or miRNA-modulation treatments. **k**, Heatmap showing inflammatory cytokine profiles after the indicated treatments. Interleukin-6 (IL-6), interleukin-12 (IL-12) and tumor necrosis factor-α (TNF-α) expression levels are shown as z-score-normalized relative expression values. **l**, Summary schematic illustrating the miR-21-associated epithelial module and miR-155-associated immune module linked to the bioactivity of M-BMDC-Exos.

To investigate molecular features associated with the donor-source-dependent activity of Exos, we next analysed exosomal miRNA profiles. We used publicly available exosomal miRNA-sequencing data from GEO dataset GSE190854 and established a reproducible multi-step prioritization workflow (**Fig. 1c**). The pipeline integrated abundance filtering, donor-source variability, biological relevance and statistical ranking. This workflow reduced the initial miRNA pool to a top-ranked candidate set, from which the top 20 miRNAs were taken forward for comparative analysis (**Fig. 1d**).

Heatmap analysis of these top 20 miRNAs revealed donor-source-dependent expression patterns across Exo groups. Several miRNAs showed marked intergroup variation, indicating that Exos derived from different donor-cell sources carried distinct miRNA signatures (**Fig. 1d**). We next performed functional annotation of these top-ranked miRNAs to examine whether they were associated with biological processes related to the phenotypes observed in the exosome screen. This analysis linked the candidate miRNA set to inflammatory responses, BMDC maturation, epithelial transition, tight-junction organization, dendritic-cell differentiation, cytokine production, immune regulation and cell-cell junction assembly. These annotated processes were consistent with the two functional properties measured in the screening system: BMDC activation and epithelial-barrier modulation. Within this framework, miR-155 and miR-21 were selected as representative candidates for downstream validation, corresponding to immune-activation-associated and epithelial-barrier-associated modules, respectively (**Fig. 1e**).

We first examined the miR-21-associated epithelial module during the early junctional-remodelling window identified by TEER analysis. Accordingly, epithelial junctional markers and repair-associated phenotypes were analysed 12 h after treatment. Immunofluorescence staining showed that M-BMDC-Exos were associated with reduced occludin, zonula occludens-1 (ZO-1) and E-cadherin signals in epithelial monolayers. Treatment with a miR-21 mimic produced a similar junctional phenotype, whereas inhibition of miR-21 in M-BMDC-Exos partially attenuated these changes **(Fig. 1f**). Scratch-wound assays showed that M-BMDC-Exos altered epithelial repair dynamics, and this effect was partially reversed after miR-21 inhibition (**Fig. 1g**). RT–qPCR analysis showed reduced *Tjp1*, *Ocln* and *Cdh1* transcript levels after M-BMDC-Exo treatment, with partial restoration after miR-21 inhibition. Consistently, western blot analysis showed decreased ZO-1, occludin and E-cadherin expression after M-BMDC-Exo treatment and partial recovery following miR-21 inhibition (**Fig. 1h**). Together, these data support a role for exosomal miR-21 in the early epithelial junctional remodelling associated with M-BMDC-Exos.

In parallel, miR-155 was assigned to an immune-activation-associated module centred on suppressor of cytokine signalling 1 (SOCS1) and SH2 domain-containing inositol 5’-phosphatase 1 (SHIP1; encoded by *Inpp5d*), two negative-feedback regulators associated with BMDC activation and inflammatory signalling. Predicted miR-155-binding sites were identified in the 3’ untranslated regions of *Socs1* and *Inpp5d* (**Fig. 1i**). SOCS1 restrains cytokine-driven Janus kinase–signal transducer and activator of transcription (JAK–STAT) signalling, whereas SHIP1 limits PI3K-dependent and Toll-like receptor (TLR)-associated inflammatory signalling. We therefore selected *Socs1* and *Inpp5d* for validation. M-BMDC-Exos reduced *Socs1* and *Inpp5d* expression, whereas miR-155 inhibition partially restored their expression (**Fig. 1j**). Cytokine profiling further showed that M-BMDC-Exos and miR-155 mimic shifted BMDCs towards a higher relative interleukin-6 (IL-6), interleukin-12 (IL-12) and tumour necrosis factor-α (TNF-α) profile, whereas miR-155 inhibition attenuated this cytokine-associated signature (**Fig. 1k**). These data support a role for exosomal miR-155 in modulating SOCS1/SHIP1-associated inhibitory pathways and BMDC activation-associated cytokine profiles. We integrated these observations into a working model of M-BMDC-Exo bioactivity (**Fig. 1l**). In this model, the miR-21-associated epithelial module links M-BMDC-Exos to epithelial junctional remodelling, as reflected by reduced ZO-1, occludin and E-cadherin expression, thereby providing a potential route for enhanced transepithelial passage. In parallel, the miR-155-associated immune module links M-BMDC-Exos to reduced *Socs1* and *Inpp5d* expression and BMDC activation. Thus, M-BMDC-Exos appear to couple epithelial-barrier modulation with dendritic-cell activation through distinct but coordinated miRNA-associated programs. These analyses identified mature M-BMDC-Exos as a bioactive exosome population with coupled epithelial-modulating and BMDC-activating activities. The associated miRNA signature further linked these two functional properties to miR-21- and miR-155-related programs, providing the rationale for using M-BMDC-Exos as the backbone for subsequent peptide-programmed mRNA delivery.

### Machine-learning-guided peptide design generates INSPIRE for efficient mRNA loading and dendritic-cell activation

Having selected M-BMDC-Exos as the bioactive backbone, we next engineered their surface to improve mRNA encapsulation and dendritic-cell engagement. We designed a modular tripartite peptide composed of a targeting domain, a nucleic-acid-binding domain and a membrane-anchoring domain. To guide peptide selection, we established a machine-learning workflow that integrated peptide-sequence featurization, supervised model training, model interpretation, virtual peptide screening and experimental validation (**Fig. 2a** and **Supplementary Table 1**).

**Fig. 2.**
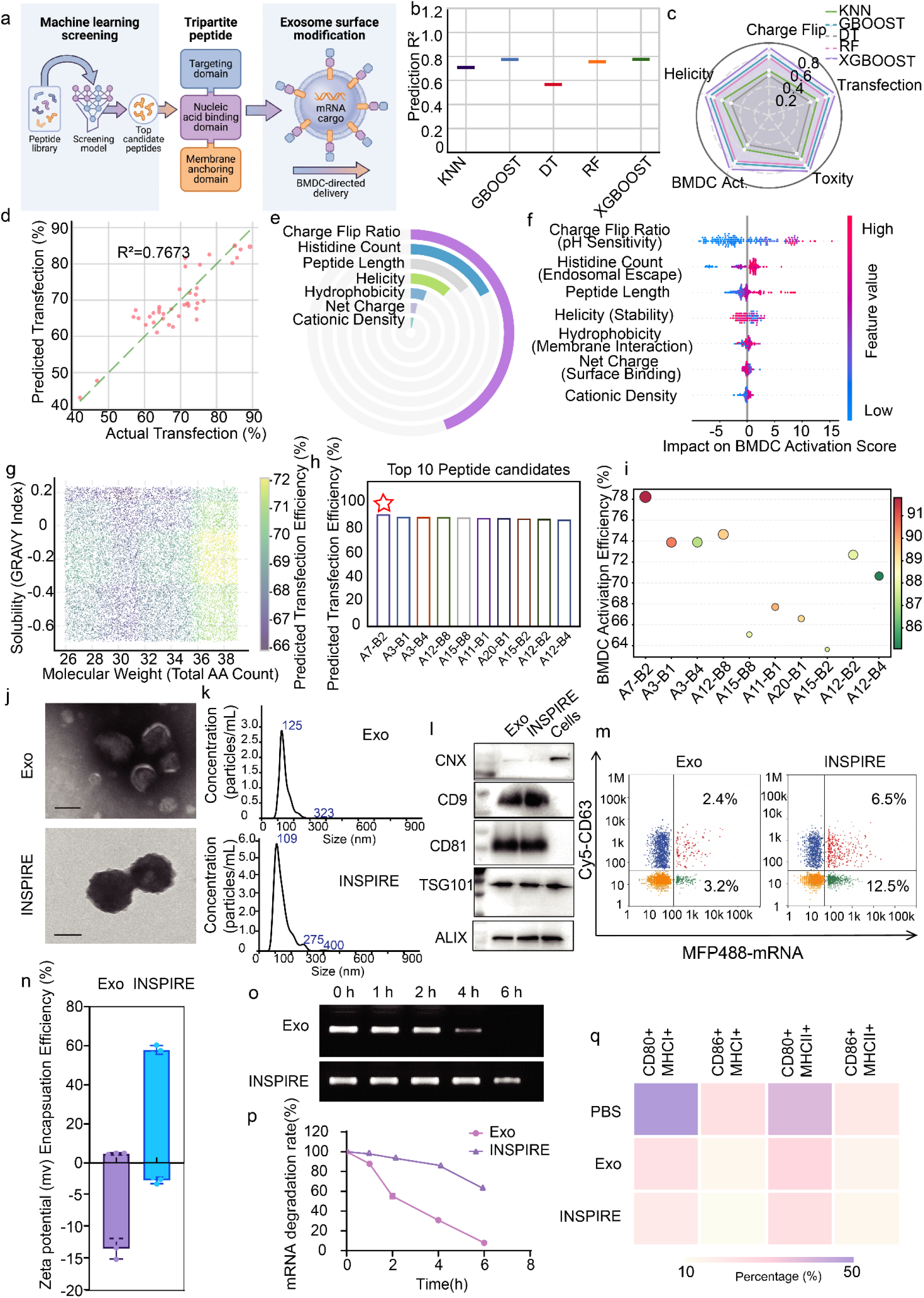
Machine-learning-guided peptide programming generates INSPIRE for mRNA loading and dendritic-cell activation. **a**, Schematic of the machine-learning-guided peptide screening and exosome surface modification strategy. Peptide candidates were generated from sequence libraries and training data, followed by prediction of peptide scores and prioritization of candidate peptide sequences. The selected tripartite peptide contained a targeting domain, a nucleic acid condensation domain and a lipid anchoring domain, enabling exosome surface modification and assembly of a nucleic-acid delivery carrier. **b**, Prediction performance of five machine-learning models for peptide screening. KNN, k-nearest neighbours; GBOOST, gradient boosting; DT, decision tree; RF, random forest; XGBOOST, extreme gradient boosting. Prediction performance is shown as the coefficient of determination, R². **c**, Radar plot comparing the model-predicted peptide-performance profiles across charge-flip behaviour, helicity, bone marrow-derived dendritic cell (BMDC) activation, transfection and toxicity-related outputs. **d**, Correlation between predicted and experimentally measured transfection efficiency in the validation set. The coefficient of determination is shown. **e**, Feature-ranking analysis of peptide descriptors contributing to model-predicted peptide performance. The ranked features include charge-flip ratio, histidine count, peptide length, helicity, hydrophobicity, net charge and cationic density. **f**, SHapley Additive exPlanations (SHAP) analysis showing the impact of individual peptide features on the BMDC activation score. Each dot represents one peptide candidate; colour indicates the feature value. **g**, Virtual screening of the peptide design space. A total of 20,000 virtual peptides were projected according to molecular weight and stability, and colour indicates model-predicted transfection efficiency. **h**, Predicted transfection efficiency of the top ten peptide candidates selected from the virtual screening output, showing the combinatorial effect of lysine/arginine- and histidine-containing peptide modules. **i**, Experimental validation of the top ten peptide candidates in BMDCs. The y-axis shows BMDC activation efficiency, and the colour scale indicates transfection-related performance. **j**, Transmission electron microscopy images of unmodified exosomes and peptide-engineered exosomes, termed INSPIRE. Scale bars, 100 nm. **k**, Nanoparticle tracking analysis of Exo and INSPIRE. **l**, Western blot analysis of exosomal and cellular markers in Exo, INSPIRE and BMDC cell lysates. Exosomal markers included cluster of differentiation 9 (CD9), cluster of differentiation 81 (CD81), tumour susceptibility gene 101 protein (TSG101) and ALG-2-interacting protein X (ALIX). Calnexin (CNX) was used as an endoplasmic-reticulum-associated negative marker for vesicle preparations. **m**, Single-particle flow-cytometric analysis of Exo and INSPIRE using CD63 labelling and MFP488-labelled mRNA. CD63 was used to identify a CD63-positive vesicle subpopulation, whereas total mRNA-positive events include both CD63-positive and CD63-low/negative vesicle-associated particles. **n**, Quantification of mRNA encapsulation efficiency and zeta potential of Exo and INSPIRE. **o**, Agarose gel electrophoresis showing mRNA stability in Exo and INSPIRE after incubation for the indicated times. **p**, Quantification of mRNA degradation rates in Exo and INSPIRE over time. **q**, Heatmap showing BMDC activation after treatment with phosphate-buffered saline (PBS), Exo or INSPIRE. The proportions of CD80⁺ major histocompatibility complex class II-positive (MHC-II⁺) and CD86⁺MHC-II⁺ BMDCs are shown.

To select a predictive model for peptide prioritization, we trained and compared five machine-learning algorithms: k-nearest neighbours (KNN), decision tree (DT), gradient boosting (GBoost), random forest (RF) and extreme gradient boosting (XGBoost). Among these models, XGBoost showed the best overall predictive performance (**Fig. 2b**, **c**) and was therefore selected for downstream screening. In the held-out validation set, model-predicted transfection efficiency showed good agreement with experimentally measured values, with a coefficient of determination of 0.7673 (**Fig. 2d**).

We next interrogated the sequence features associated with model prediction to improve interpretability. XGBoost feature-importance ranking identified charge-flip ratio, histidine content, peptide length, helicity, hydrophobicity, net charge and cationic density as influential descriptors of peptide performance (**Fig. 2e**). SHAP analysis further indicated that charge-flip ratio and histidine content made prominent contributions to the model output, whereas hydrophobicity, helicity, peptide length and cationic density showed feature-dependent effects (**Fig. 2f**). These results provided a feature-level basis for virtual peptide screening.

Using the trained XGBoost model as a prioritization tool, we screened a virtual peptide design space containing 20,000 candidate sequences. Candidate peptides were projected according to total amino-acid count and solubility-related GRAVY index (**Fig. 2g**). Within the screened design space, high-scoring candidates were enriched in a physicochemical region characterized by peptide lengths of 36–38 amino acids and GRAVY index values between −0.4 and −0.2. This region was therefore used to prioritize peptide sequences for experimental validation (**Fig. 2g, h**).

From this prioritized region, the top 10-ranked peptide candidates were selected for synthesis and empirical testing. Their model-predicted transfection efficiencies were first compared (**Fig. 2h**). These candidates were then experimentally evaluated in BMDCs using a validation matrix that integrated measured mRNA transfection and BMDC activation (**Fig. 2i**). This analysis identified A7-B2 as the lead peptide, because it showed a favourable combined profile across model-predicted transfection performance and experimentally measured BMDC activation (**Fig. 2i; Supplementary Table 1**). A7-B2 was therefore used to modify M-BMDC-Exos and generate the standard INSPIRE formulation for subsequent experiments.

We next characterized Exo and INSPIRE by transmission electron microscopy, unmodified exosomes (Exo) displayed a typical smooth, cup-shaped vesicular morphology. Following peptide modification, INSPIRE showed a slightly altered surface appearance after peptide engineering while maintaining overall vesicle integrity (**Fig. 2j**). Nanoparticle tracking analysis (NTA) revealed comparable nanoscale size distributions, with modal size peaks of approximately 125 nm for Exo and 109 nm for INSPIRE (**Fig. 2k**). Western blot analysis confirmed the enrichment of canonical exosomal markers, including cluster of differentiation 9 (CD9), cluster of differentiation 81 (CD81), tumour susceptibility gene 101 protein (TSG101) and ALG-2-interacting protein X (ALIX), in both Exo and INSPIRE, whereas calnexin (CNX), an endoplasmic-reticulum-associated protein, was absent from the vesicle preparations (**Fig. 2l**).

To examine mRNA association with engineered vesicle preparations at the single-particle level, we performed single-particle flow cytometry using CD63 labelling and MFP488-labelled mRNA. Compared with unmodified Exo, INSPIRE increased the CD63⁺mRNA⁺ particle fraction from 2.4% to 6.5%, indicating enhanced mRNA association within the CD63-positive vesicle subpopulation. The overall mRNA-positive particle population also increased from 5.6% in Exo to 19.0% in INSPIRE. Because CD63 represents only one exosomal marker and exosome preparations are marker heterogeneous, this broader mRNA-positive population was considered together with the CD63⁺mRNA⁺ fraction when evaluating vesicle-associated mRNA signals (**Fig. 2m**). These data support increased mRNA association with INSPIRE at the single-particle level.

Bulk loading analysis showed that INSPIRE retained a higher fraction of input mRNA than unmodified Exo, increasing encapsulation efficiency from approximately 5% to 58%. Peptide engineering also shifted the zeta potential towards a less negative surface charge (**Fig. 2n**). Agarose gel electrophoresis showed that INSPIRE more effectively protected loaded mRNA from degradation over time (**Fig. 2o**). Quantification of residual mRNA further confirmed slower mRNA degradation in INSPIRE during the 6 h incubation period (**Fig. 2p**). These results indicate that peptide engineering improves mRNA association, bulk loading and short-term mRNA stability within the engineered exosome formulation.

Finally, we examined whether the ML-selected peptide also translated into the intended BMDC-activation output used during peptide prioritization. Flow-cytometric profiling showed that INSPIRE increased BMDC populations co-expressing activation markers and MHC molecules, including CD80⁺MHC-I⁺, CD86⁺MHC-I⁺, CD80⁺MHC-II⁺ and CD86⁺MHC-II⁺ subsets, compared with PBS and unmodified Exo controls **(Fig. 2q)**. These results indicate that A7-B2 engineering not only improves mRNA association and loading, but also enhances the BMDC activation-associated phenotype of the engineered formulation.

Together, these data show that ML-guided A7-B2 peptide engineering converts M-BMDC-Exos into INSPIRE, an mRNA-associated exosome formulation with improved mRNA loading, enhanced short-term mRNA stability, increased transfection efficiency and strengthened BMDC activation-associated marker expression. INSPIRE therefore combines the bioactive M-BMDC-Exo backbone with peptide-enabled mRNA loading and BMDC-activating properties, supporting its use in subsequent intranasal mRNA vaccination studies.

### INSPIRE enhances dendritic-cell engagement and pulmonary mRNA expression with favorable safety after intranasal delivery

To evaluate how peptide engineering improves the cellular events required for pulmonary mRNA delivery, we first examined transepithelial transport using an Ussing chamber model. Compared with Exo, INSPIRE increased Cy5-labelled mRNA transport into the receptor chamber and produced a higher apparent permeability coefficient, indicating enhanced transepithelial mRNA delivery by INSPIRE (**Extended Data Fig. 1a-c**). We next focused on DC2.4 cells to determine whether improved epithelial passage was accompanied by enhanced uptake by antigen-presenting cells. Flow-cytometric analysis showed that INSPIRE increased intracellular Cy5-mRNA accumulation compared with Exo, as reflected by higher fluorescence intensity and/or increased mRNA-positive DC2.4 cells (**Extended Data Fig. 1d**). To define the entry routes underlying this uptake advantage, we treated DC2.4 cells with pharmacological inhibitors of endocytosis. INSPIRE uptake was markedly reduced at 4 °C and after sodium azide treatment, indicating an active, energy-dependent internalization process. The strong inhibition by protamine and amiloride suggested that charge-dependent membrane engagement and macropinocytosis were major contributors to INSPIRE entry, whereas the partial sensitivity to chlorpromazine and filipin indicated auxiliary involvement of clathrin-mediated and lipid-raft/caveolae-associated uptake routes (**Fig. 3a**).

**Fig. 3.**
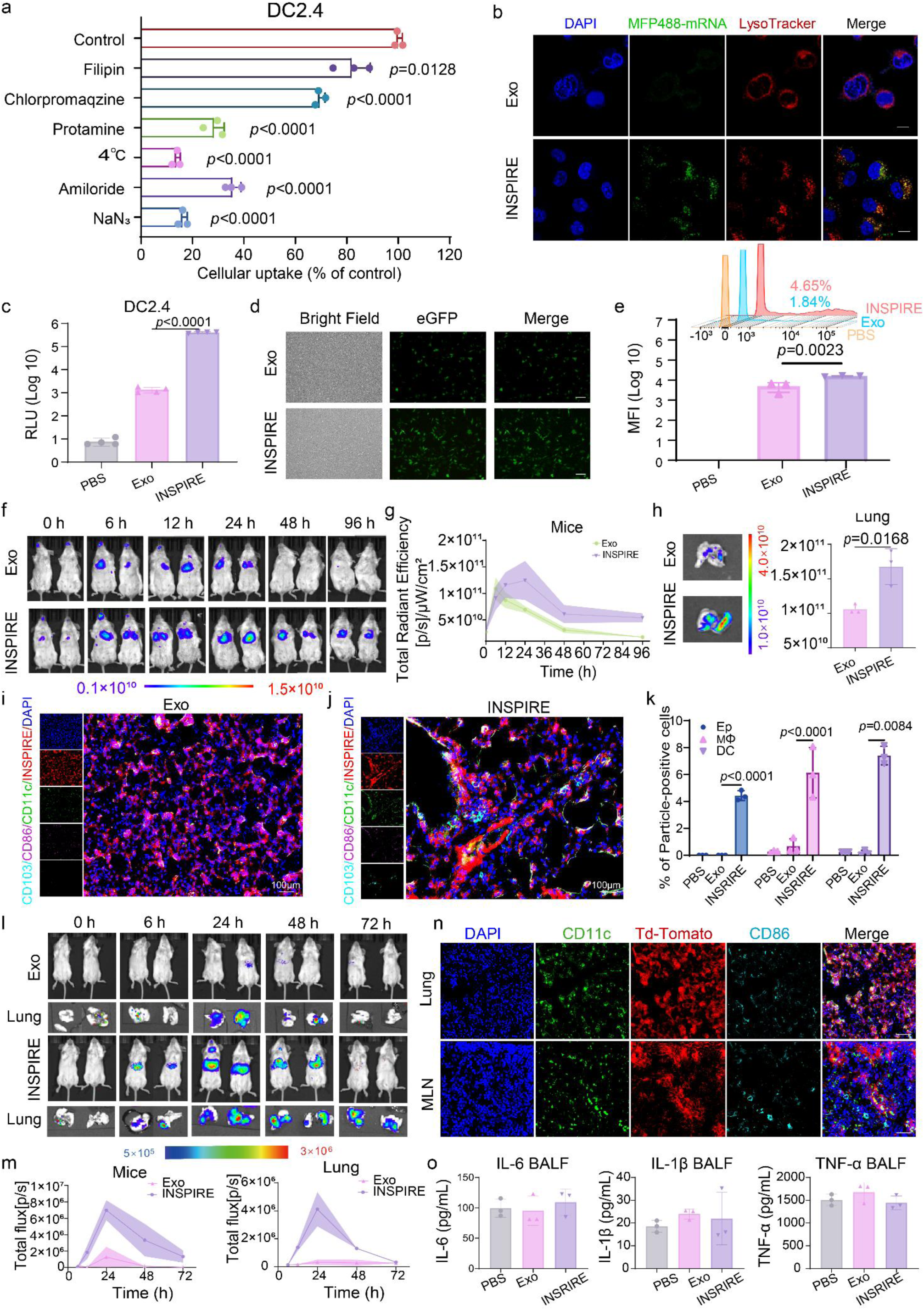
Evaluations of INSPIRE towards pulmonary biodistribution, mRNA expression and safety profiles. **a**, Cellular uptake of INSPIRE in DC2.4 cells after treatment with the indicated endocytosis or uptake inhibitors. Uptake was normalized to the untreated control group and is shown as percentage of control. **b**, Confocal images of DC2.4 cells treated with MFP488-labelled mRNA delivered by INSPIRE or Exo. Nuclei were stained with DAPI (blue), MFP488-labelled mRNA is shown in green and lysosomes were stained with LysoTracker (red). Scale bars, 10 μm. **c**, Firefly luciferase activity in DC2.4 cells after treatment with PBS, Exo or INSPIRE loaded with mRNA encoding firefly luciferase (mFluc). **d**, Bright-field and fluorescence images of DC2.4 cells after treatment with Exo or INSPIRE loaded with mRNA encoding enhanced green fluorescent protein (mEGFP). Scale bars, 10 μm. **e**, Flow-cytometric analysis and quantification of enhanced green fluorescent protein (EGFP) expression in DC2.4 cells after treatment with PBS, Exo or INSPIRE. **f**, In vivo fluorescence imaging of mice after intranasal administration of DiR-labelled Exo or INSPIRE at the indicated time points. **g**, Quantification of fluorescence intensity in the lung from panel **f**. **h**, Representative ex vivo fluorescence images of isolated lungs collected 96 h after intranasal administration of DiR-labelled Exo or INSPIRE as in panel **f**. **i, j,** Representative immunofluorescence images of lung sections after intranasal administration of DiR-labelled Exo (**i**) or INSPIRE (**j**). Nuclei were stained with DAPI (blue), particle-associated signals are shown in red, CD11c in green, CD86 in magenta and CD103 in cyan. Scale bars, 100 μm. **k**, Quantification of particle-positive epithelial cells (Ep), macrophages (Mφ) and dendritic cells (DC) in the lung after treatment with PBS, Exo or INSPIRE. **l**, In vivo and ex vivo bioluminescence imaging of mice and lungs after intranasal administration of mFluc formulated with Exo or INSPIRE at the indicated time points. **m**, Quantification of total bioluminescence flux in whole mice and isolated lungs from panel **l**. **n**, Representative immunofluorescence images of lung and mediastinal lymph node (MLN) sections collected 3 d after intranasal administration of Cre recombinase mRNA in tdTomato reporter mice. Nuclei were stained with DAPI (blue), CD11c is shown in green, tdTomato is shown in red and CD86 is shown in cyan. Scale bar, 100 μm. o, Bronchoalveolar lavage fluid (BALF) levels of IL-6, IL-1β and TNF-α measured after treatment with PBS, Exo or INSPIRE.

We next assessed intracellular mRNA trafficking. INSPIRE-treated DC2.4 cells showed stronger intracellular MFP488-mRNA signals and reduced co-localization with LysoTracker-positive compartments compared with Exo-treated cells, indicating enhanced mRNA accumulation and endolysosomal release (**Fig. 3b**). Consistent with improved intracellular delivery, INSPIRE-mediated Firefly luciferase mRNA (mFluc) delivery produced higher luciferase activity than Exo in DC2.4 cells (**Fig. 3c**). For enhanced green fluorescent protein mRNA (mEGFP) delivery, fluorescence imaging showed stronger EGFP signals in INSPIRE-treated cells, and flow cytometry confirmed increased EGFP-positive cells and higher MFI compared with Exo (**Fig. 3d, e**). Cell Counting Kit-8 (CCK-8) assays showed negligible cytotoxicity in DC2.4 cells after INSPIRE treatment (**Extended Data Fig. 1e**).

We then investigated whether these delivery advantages translated into in vivo scenario after intranasal administration. In vivo imaging of DiR-labelled formulations revealed lung-predominant distribution, with INSPIRE producing higher pulmonary fluorescence signals than Exo over the observation period; this pattern was further confirmed by ex vivo imaging of isolated lungs (**Fig. 3f-h**). Immunofluorescence imaging of lung sections showed substantially stronger INSPIRE signals than Exo, with increased spatial co-localization with CD11c, CD86 and CD103 in pulmonary dendritic-cell-enriched regions (**Fig. 3i, j**). These data suggest that INSPIRE preferentially associates with activated CD103⁺CD11c⁺ dendritic-cell compartments in the lung after intranasal administration. Consistently, quantification across pulmonary cell populations showed that INSPIRE-associated signals were detected primarily in dendritic cells, followed by macrophages and epithelial cells (**Fig. 3k**). mFluc reporter imaging further showed that INSPIRE generated markedly stronger bioluminescence signals than Exo in both live mice and isolated lungs, with peak expression at 24 h and detectable signals persisting up to 72 h after dosing (**Fig. 3l, m**). To determine cell types successfully transfected by INSPIRE, Rosa26-LSL-tdTomato mice, in which tdTomato expression is induced upon Cre recombinase-mediated recombination, were used to trace in vivo Cre mRNA (mCre) transfection. Immunofluorescence analysis showed INSPIRE induced broader tdTomato reporter activation in the lung and mediastinal lymph node (MLN) than Exo, with increased overlap with CD11c and CD86 staining **(Fig. 3n and Extended Data Fig. 1f**). Flow-cytometric analysis further confirmed increased tdTomato reporter signals in pulmonary dendritic cells and macrophages after INSPIRE mediated mCre delivery (**Extended Data Fig. 1g**). To assess in vivo safety profile of INSPIRE, bronchoalveolar lavage fluid (BALF) cytokines were measured. Levels of interleukin-6 (IL-6), interleukin-1β (IL-1β) and tumor necrosis factor-α (TNF-α) remained comparable to PBS or Exo controls (**Fig. 3o**). Consistently, histopathology analysis of major organs, including the lung, liver, spleen, brain, kidney and heart, showed no obvious pathological abnormalities after INSPIRE administration (**Extended Data Fig. 2a**). Serum levels of IL-6, IL-1β and TNF-α, measured as indicators of systemic inflammatory responses, remained at levels that are comparable to PBS controls across Exo and INSPIRE groups (**Extended Data Fig. 2b**). Likewise, serum biochemical indices associated with hepatic, renal and tissue injury, also remained comparable across groups (**Extended Data Fig. 2c**), further supporting the biocompatibility of INSPIRE.

### INSPIRE elicits durable mucosal antibody responses and IL-17-skewed T-cell immunity against Kp

We established an mRNA vaccine (Kp-mRNA) encoding two Kp-derived antigen candidates, carbapenemase KPC-2 and peptidoglycan-associated lipoprotein Pal, which represent complementary resistance- and virulence-associated targets. KPC-2 is a prevalent carbapenemase in carbapenem-resistant Kp, whereas Pal is a cell-envelope-associated lipoprotein linked to envelope integrity and virulence. Fusion-protein vaccination with KPC-Pal (**Supplementary Fig. 1a, Supplementary Table. 2**) has previously been shown to protect mice against pulmonary K. pneumoniae infection. ^48^ BALB/c mice received a prime-boost regimen on days 0 and 21, with lipid nanoparticle intramuscular immunization (LNP-i.m.) and unmodified exosome intranasal immunization (Exo-i.n.) as controls. Using the immunization scheme shown in **Fig. 4a**, INSPIRE-i.n. induced robust serum IgG responses after boosting, with endpoint titres reaching up to 2,560,000; these titres were broadly comparable to those induced by LNP-i.m., higher than those induced by Exo-i.n. and remained recallable after sublethal challenge of YBQ (1×10^6^) on day 730 (**Fig. 4b**). On day 28, INSPIRE-i.n. increased mucosal sIgA titres in bronchoalveolar lavage fluid (BALF) and nasal lavage fluid (NALF) to up to 3,200 and 512, respectively, compared to 10 and 8 in Exo-i.n. counterparts (**Fig. 4c**). After the Day 730 recall challenge, INSPIRE-i.n. maintained higher sIgA titres at day 737 in BALF and NALF, reaching up to 6,400 and 512, respectively, compared with 1,600 and 32 in Exo-i.n. controls (**Fig. 4d**). Mucosal IgG followed a similar pattern, being elevated both at day 28, and at day 737 after the day 730 sublethal YBQ recall challenge, with BALF and NALF IgG titres reaching up to 12,800 and 256, respectively (**Supplementary Fig. 1b**).

**Fig. 4.**
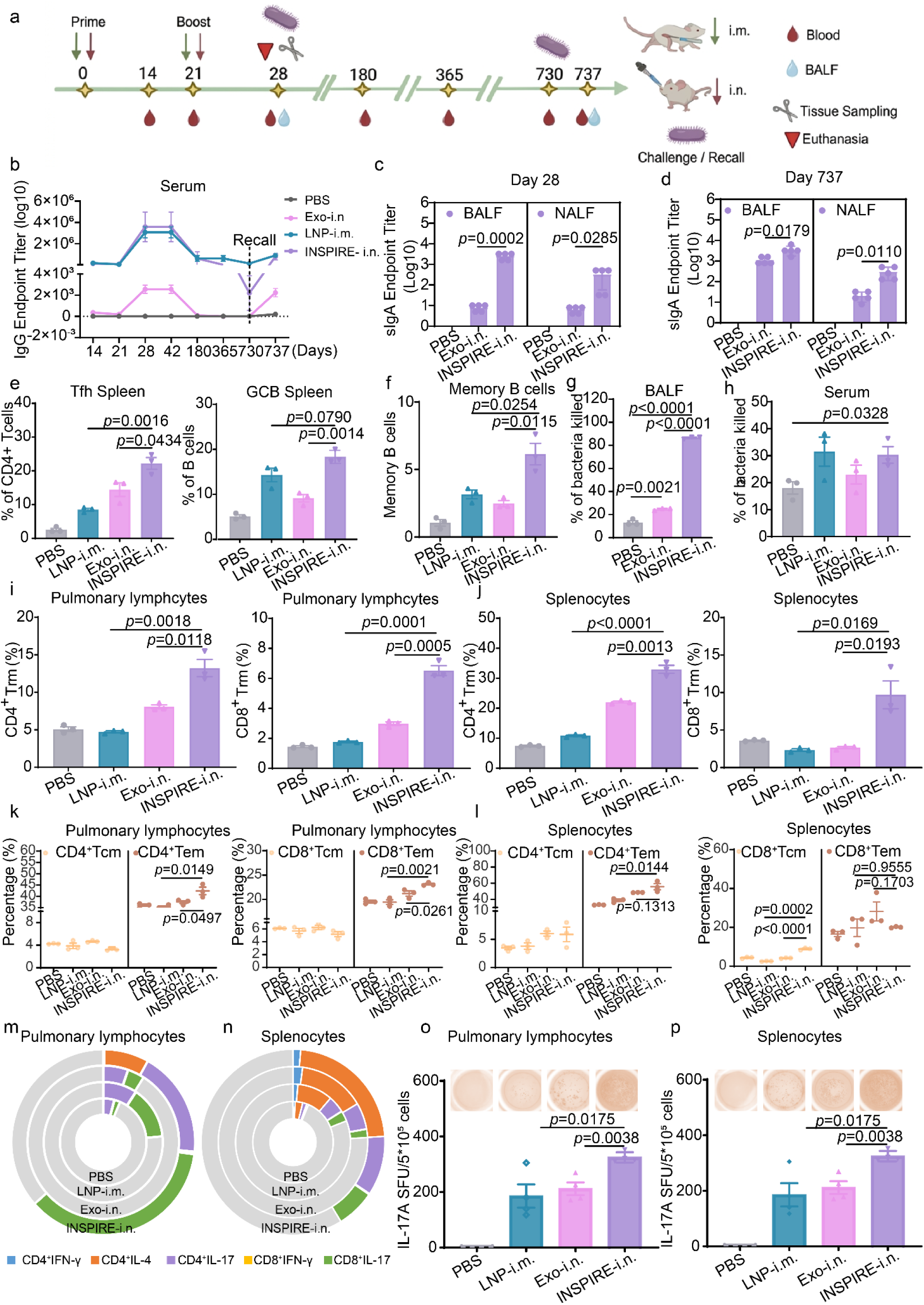
INSPIRE induces durable mucosal antibody responses and IL-17-skewed T-cell immunity against Kp. **a,** Schematic illustration of the immunization, sampling and bacterial challenge timeline. BALB/c mice were immunized intranasally (i.n.) with INSPIRE or unmodified Exo formulated with mRNA encoding the Kp antigen, whose sequence information is provided in **Supplementary Table 2**. Lipid nanoparticle intramuscular immunization (LNP-i.m.) and PBS treatment served as controls. Mice received a prime-boost regimen on days 0 and 21. Long-term recall responses were assessed after sublethal intratracheal challenge with Kp YBQ on day 730, followed by analysis on day 737. **b**, Endpoint titres of Kp antigen-specific serum immunoglobulin G (IgG) collected on days 14, 21, 28, 42, 180, 365, 730 and 737 after prime immunization. The day-737 sample was collected 7 d after sublethal YBQ challenge on day 730 (n = 5 biologically independent animals). **c**, **d**, Secretory immunoglobulin A (sIgA) endpoint titres in bronchoalveolar lavage fluid (BALF) and nasal lavage fluid (NALF) on day 28 (**c**) and day 737 after sublethal YBQ challenge (**d**) (n = 5 biologically independent animals). **e**, Flow-cytometric quantification of splenic T follicular helper (Tfh) cells and germinal-centre B (GCB) cells on day 7 after boost immunization (n = 3 biologically independent animals). **f**, Percentage of Kp antigen-specific memory B cells on day 28 after prime immunization (n = 3 biologically independent animals). **g**, **h**, Opsonophagocytic killing (OPK) activity against Kp YBQ mediated by bronchoalveolar lavage fluid (**g**) and serum (**h**) from immunized mice (n = 3 biologically independent animals). **i**, **j**, Flow-cytometric quantification of CD4⁺ and CD8⁺ tissue-resident memory T (T_RM_) cells in pulmonary lymphocytes (**i**) and splenocytes (**j**) on day 42 after prime immunization (n = 3 biologically independent samples). **k**, **l**, Flow-cytometric quantification of CD4⁺ and CD8⁺ central-memory T (T_CM_) and effector-memory T (T_EM_) cells in pulmonary lymphocytes (**k**) and splenocytes (**l**) on day 28 after prime immunization (n = 3 biologically independent samples). **m**, **n**, Proportions of cytokine-positive CD4⁺ and CD8⁺ T cells in pulmonary lymphocytes (**m**) and splenocytes (**n**) on day 28 after 12 h restimulation with Kp antigen. CD4⁺ T-cell subsets were analysed for interferon-γ (IFN-γ), interleukin-4 (IL-4) and interleukin-17 (IL-17), and CD8⁺ T-cell subsets were analysed for IFN-γ and IL-17 (n = 3 biologically independent animals). Each ring corresponds to one immunization group, and a complete ring denotes 20% cytokine-positive cells within the corresponding CD4⁺ or CD8⁺ T-cell population. **o**, **p**, Enzyme-linked immunospot (ELISpot) analysis of IL-17A-secreting cells in pulmonary lymphocytes (**o**) and splenocytes (**p**) after 12 h restimulation with Kp antigen (n = 3 biologically independent animals). For all experiments, the mRNA dose in Exo and INSPIRE formulations was 2 μg per mouse.

To assess whether this humoral profile was accompanied by cellular features associated with antibody maturation and recall, we found INSPIRE-i.n. substantially increased T follicular helper (Tfh) cells, germinal-centre B (GCB) cells (**Fig. 4e, Supplementary Fig. 1c, d**) and memory B cells relative to the comparator groups (**Fig. 4f**). These cellular changes were accompanied by functional antibody activity, as BALF from INSPIRE-i.n.-immunized mice showed near-complete opsonophagocytic killing, while serum also displayed enhanced antibacterial activity relative to PBS and comparator groups (**Fig. 4g, h**), indicating that the antibody response was not only stronger in magnitude but also effective in vitro bacterial control.

Considering previous studies suggest protection against respiratory bacteria also depends on local T-cell memory ^11^, we next assessed tissue-resident and effector-memory compartments. INSPIRE-i.n. markedly expanded both CD4⁺ and CD8⁺ T_RM_ cells in the lung, with parallel increases also detected in splenocytes (**Fig. 4i, j**). Pulmonary CD4⁺ effector memory T cells (T_EM_) and CD8⁺ T_EM_ cells were likewise increased, whereas in the spleen the advantage over Exo-i.n. was retained but the difference from LNP-i.m. was less pronounced (**Fig. 4k, l**), consistent with the stronger systemic bias of intramuscular immunization. As T_RM_ cells are positioned for rapid local recall at sites of pathogen entry, this pattern suggests that intranasal INSPIRE preferentially strengthens the immune compartment most relevant to airway reinfection.

We further evaluated cytokine-polarized T-cell responses in pulmonary lymphocytes. INSPIRE-i.n. selectively amplified IL-17-producing T-cell subsets detected by intracellular cytokine staining, increasing CD4⁺IL-17⁺ cells to 4.80% compared with 0.05-0.15% in the PBS, LNP-i.m. and Exo-i.n. groups, and CD8⁺IL-17⁺ cells to 5.71% compared with 0.08-0.43% in other counterparts. By contrast, IFN-γ⁺ CD4⁺ and CD8⁺ T cells remained at much lower frequencies after INSPIRE-i.n. vaccination, reaching 0.070% and 0.033%, respectively, while CD4⁺IL-4⁺ cells were minimal across groups (**Fig. 4m**). Thus, INSPIRE-i.n. induced a pulmonary T-cell profile dominated by IL-17-producing subsets, including a prominent CD8⁺IL-17⁺ population.

Although the CD8⁺IL-17⁺ bias was less pronounced compared to the lung, INSPIRE-i.n. also increased IL-17-producing T cells in splenocytes, with CD4⁺IL-17⁺ cells reaching 4.58% in the INSPIRE-i.n. group compared with 0.35-1.45% in the control groups whereas CD8⁺IL-17⁺ cells increasing to 2.40% compared with 0.14-0.68% in the control groups (**Fig. 4n**). Enzyme-linked immunospot (ELISpot) analysis further showed that INSPIRE-i.n. induced the highest frequencies of IL-17A-secreting cells in both pulmonary lymphocytes and splenocytes (**Fig. 4o, p, Supplementary Fig. 1e, 1f**).

Together, these results show that intranasal INSPIRE induced early mucosal antibody responses, recallable humoral immunity, antibody-associated B-cell compartments and reinforced lung-resident T-cell memory. Most notably, INSPIRE-i.n. preferentially amplified pulmonary IL-17-producing T-cell responses, while ELISpot confirmed increased IL-17A-secreting cells, with CD8⁺IL-17⁺ cells emerging as a prominent population.

### INSPIRE-based mRNA vaccination confers robust cross-strain protection against pulmonary Kp infection

Given the promising immune responses elicited by INSPIRE, we next evaluated protective efficacy against pulmonary Kp infection. Kp-mRNA was formulated in INSPIRE, with Exo-i.n. and LNP-i.m. as controls, according to the immunization schedule shown in **Fig. 4a**. Mice were challenged intratracheally with three clinically relevant Kp strains representing distinct capsular polysaccharide (CPS) serotypes: YBQ (K20), A7818 (K2) and YYD (K1). Detailed strain information is provided in the Methods. YBQ, a K20 clinical isolate preserved in our laboratory and isolated from the First Affiliated Southwest Hospital of Army Medical University; YYD, a K1 isolate from the same hospital; and A7818, a K2 isolate from Guangxi Medical University. Across all three challenge models, INSPIRE-i.n. INSPIRE-i.n. attenuated body-weight loss and promoted recovery among surviving mice compared with PBS, LNP-i.m. and Exo-i.n. groups (**Extended Data Fig. 3a**). This benefit was most evident in survival outcomes: INSPIRE-i.n. achieved 100% survival against YBQ and A7818 and retained approximately 80% survival against the highly virulent YYD challenge, whereas PBS controls rapidly succumbed and Exo-i.n. provided incomplete and strain-dependent protection. LNP-i.m. provided only partial protection, with survival remaining no higher than approximately 30% by day 7 (**Extended Data Fig. 3b**). Consistently, INSPIRE-i.n. maintained the lowest global disease scores over 7 days post-challenge (dpc) approaching baseline by day 7 (**Extended Data Fig. 3c**).

We next examined lung pathology and bacterial control in the YBQ pulmonary challenge model. Histological analysis of lung sections at 2 dpc showed severe inflammatory consolidation and alveolar destruction in PBS, with partial alleviation being observed in LNP-i.m. and Exo-i.n. groups. In contrast, INSPIRE-i.n. vaccinated mice displayed largely preserved alveolar architecture with minimal inflammatory infiltration (**Extended Data Fig. 3d**). Quantification of bacterial burdens in lungs further demonstrated that INSPIRE-i.n. not only reduced pulmonary colonization but also effectively blocked dissemination, as the high lung bacterial loads observed in controls were accompanied by clear lung-to-spleen seeding and detectable bacteremia, whereas INSPIRE-i.n. drove near-complete suppression of bacterial recovery in spleen and blood (**Extended Data Fig. 3e**). Finally, INSPIRE-i.n. most effectively restrained pulmonary cytokine surges, yielding the lowest IL-6, IL-1β, and TNF-α levels among all tested groups (**Extended Data Fig. 3f**).

Together, these data show that intranasal INSPIRE vaccination provides strong cross-strain protection against pulmonary Kp infection, with superior survival, reduced lung injury, lower pulmonary bacterial burdens and limited systemic dissemination.

### INSPIRE imprints a lung-localized type-17 and tissue-residency program

To define the pulmonary transcriptional program associated with INSPIRE-induced type-17 and tissue-resident mucosal immunity, we collected lung tissues from mice immunized with INSPIRE-i.n., Exo-i.n. or PBS and performed bulk RNA sequencing (RNA-seq). Differential expression analysis identified 1,750 upregulated and 655 downregulated genes in INSPIRE-i.n. lungs compared with Exo-i.n. lungs (*p*< 0.05; **Fig. 5a**). Following the strategy used to minimize formulation-background effects, Venn analysis was applied to remove Exo-associated baseline changes. This analysis retained 713 INSPIRE-versus-Exo-specific genes and 1,279 genes shared between INSPIRE versus Exo and INSPIRE versus PBS, yielding a INSPIRE-associated differentially expressed gene (DEG) set for downstream analysis (**Fig. 5b**).

**Fig. 5.**
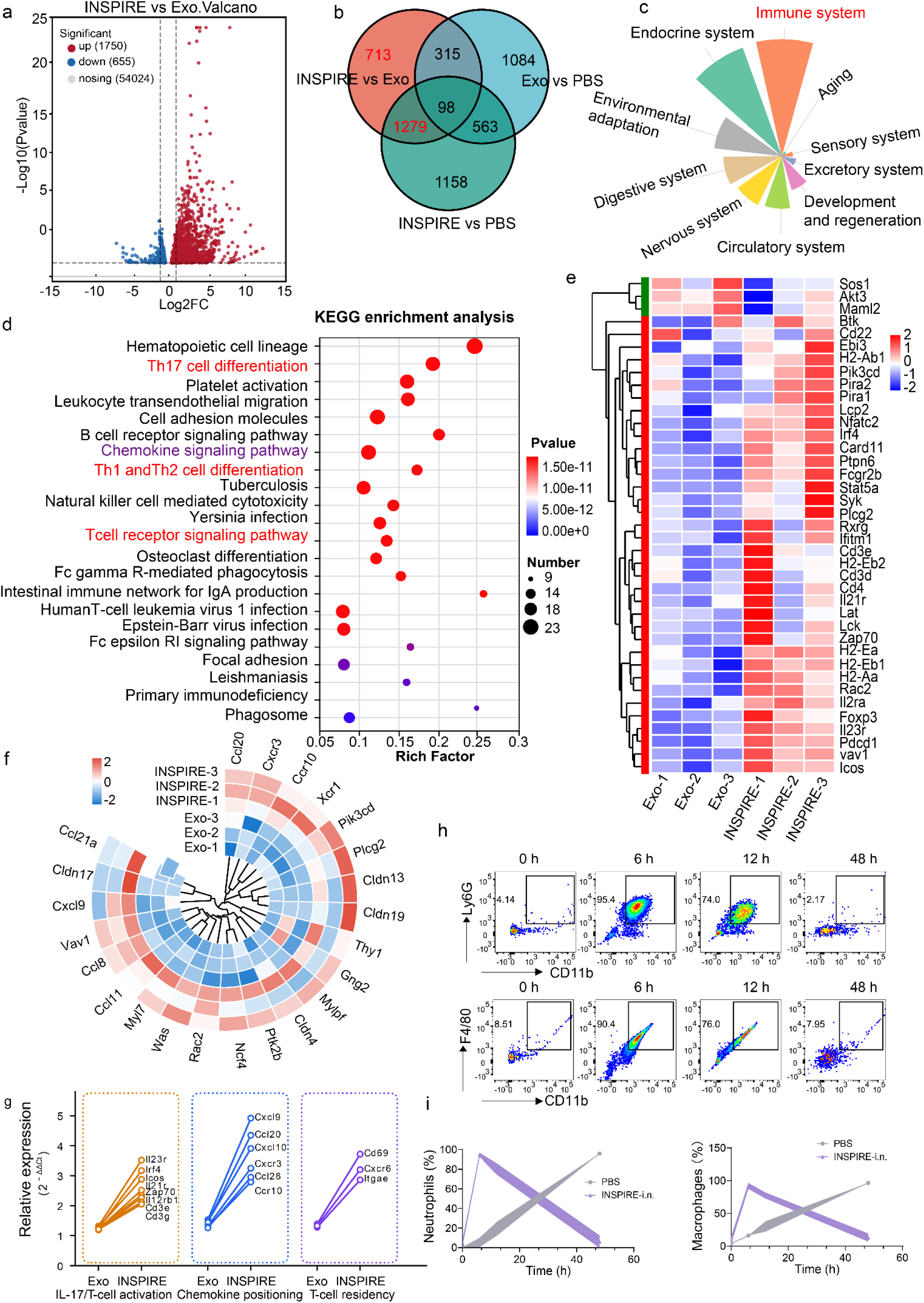
INSPIRE imprints a Tc17-compatible type-17, chemokine-positioning and tissue-residency programme and accelerates myeloid recall in the lung. **a**, Volcano plot of differentially expressed genes (DEGs) in lung tissues from INSPIRE-i.n.-immunized mice compared with Exo-i.n. counterparts. Red dots indicate significantly upregulated genes, blue dots indicate significantly downregulated genes and grey dots indicate non-significant genes (*p* < 0.05). **b**, Venn diagram showing the overlap of DEGs among INSPIRE versus Exo, INSPIRE versus PBS and Exo versus PBS comparisons. The INSPIRE-associated DEG set was generated by retaining 713 INSPIRE-versus-Exo-specific genes and 1,279 genes shared between INSPIRE-versus-Exo and INSPIRE-versus-PBS comparisons. **c**, Functional classification of the INSPIRE-associated DEG set defined in **b**, showing enrichment across biological systems, with immune-system-related terms prominently represented. **d**, Kyoto Encyclopedia of Genes and Genomes (KEGG) pathway enrichment analysis of the INSPIRE-associated DEG set. T-cell-related pathways, including Th17 cell differentiation, Th1 and Th2 cell differentiation and T-cell receptor signalling, are highlighted in red, and the chemokine signalling pathway is highlighted in purple. **e**, Hierarchical clustering heatmap of representative genes contributing to the T-cell-related pathways highlighted in d, including T-cell receptor signalling components and cytokine-responsive genes. **f**, Circular heatmap of representative genes contributing to the chemokine signalling pathway highlighted in d, including CXCR3-associated and mucosal trafficking-related chemokine cues. **g**, qPCR validation of selected genes in lung tissues after immunization. Genes were grouped into three modules: IL-17/T-cell activation-related genes, including *Irf4*, *Icos*, *Il23r*, *Il21r*, *Il12rb1*, *Cd3e*, *Cd3g* and *Zap70*; chemokine-positioning genes, including *Cxcr3*, *Cxcl9*, *Cxcl10*, *Ccl20* and *Ccr6*; and T-cell residency-associated genes, including *Cd69*, *Itgae* and *Cxcr6*. *Itgae* encodes tissue-residency marker CD103. Gene expression was normalized to *Gapdh*. **h**, Representative flow-cytometry plots showing Ly6G⁺CD11b⁺ neutrophils and F4/80⁺CD11b⁺ macrophages in BALF at the indicated time points after sublethal YBQ challenge. **i**, Time-course quantification of neutrophils and macrophages in BALF from PBS or INSPIRE-i.n. immunized mice after sublethal YBQ challenge.

Pathway-classification analysis showed that immune-system-related terms were prominently represented within the INSPIRE-associated DEG set (**Fig. 5c**). Kyoto Encyclopedia of Genes and Genomes (KEGG) enrichment further highlighted T-cell-related pathways, including Th17 cell differentiation, Th1 and Th2 cell differentiation and T-cell receptor signalling, together with chemokine signalling, B-cell receptor signalling and intestinal immune network for IgA production (**Fig. 5d**). Hierarchical clustering of T-cell-related genes showed coordinated induction of T-cell receptor signalling components and cytokine-responsive genes, including *Cd3d*, *Cd3e*, *Lat*, *Lck*, *Zap70*, *Il21r* and *Il23r*, defining a transcriptional background compatible with IL-17-producing T-cell activation rather than nonspecific pulmonary inflammation (**Fig. 5e**). This T-cell module paralleled the expansion of pulmonary IL-17⁺ T-cell subsets, particularly CD8⁺IL-17⁺ cells, observed after INSPIRE-i.n. vaccination (**Fig. 4m**).

Circular heatmap analysis of chemokine-related genes further revealed a chemokine-positioning signature in INSPIRE-i.n. lungs, including the Cxcr3-Cxcl9/Cxcl10 axis and mucosal trafficking-related genes such as *Ccl20*, *Ccr10* and *Ccl28* (**Fig. 5f**). qPCR validation further resolved this signature into three linked modules: an IL-17 / T-cell activation module, including *Irf4*, *Icos*, *Il23r*, *Il21r*, *Il12rb1*, *Cd3e*, *Cd3g* and *Zap70*; a chemokine-positioning module, including *Cxcr3*, *Cxcl9*, *Cxcl10*, *Ccl20*, and *Ccr6*; and a tissue-residency-associated module, including *Cd69*, *Itgae* and *Cxcr6* (**Fig. 5g**). These bulk lung qPCR data support a transcriptional environment enriched for T-cell activation, type-17-associated cytokine responsiveness, chemokine positioning and tissue-residency-associated signals.

To spatially validate this tissue-associated type-17 programme, we further examined CD69 and IL-17A signals in the lung and mediastinal lymph nodes. Immunofluorescence imaging showed increased CD69 and IL-17A co-localization after INSPIRE-i.n. vaccination in both tissues, supporting a tissue-associated type-17 response across the lung and mediastinal lymph nodes (**Supplementary Fig. 2a, b)**. Flow-cytometric analysis further showed that INSPIRE-i.n. increased CD69⁺IL-17⁺ memory T-cell populations within both CD8⁺ and CD4⁺ compartments in pulmonary lymphocytes and mediastinal lymph nodes (**Supplementary Fig. 2c-f**). These data provide spatial and cellular support for a tissue-associated IL-17⁺ memory T-cell response, with CD8⁺IL-17⁺ cells representing a prominent Tc17-like component within this programme.

Given the known roles of IL-17-associated responses in antibacterial neutrophil recruitment and chemokine-positioning axes, including CXCR3-CXCL9/CXCL10 and CCL20–CCR6, in pulmonary and mucosal T-cell positioning, ^49^ we next examined airway cellular recall after YBQ challenge. After sublethal YBQ challenge in mice vaccinated for 28 days, flow cytometry of BALF showed rapid accumulation of Ly6G⁺CD11b⁺ neutrophils and F4/80⁺CD11b⁺ macrophages in INSPIRE-i.n.-immunized mice at 6 h, followed by a decline by 48 h (**Fig. 5h, i**). In PBS-treated mice, myeloid-cell accumulation was delayed and remained elevated at 48 h (**Fig. 5i** and **Supplementary Fig. 2g**). This kinetic pattern indicated that INSPIRE-i.n. vaccination promoted a rapid and more controlled airway myeloid response after bacterial exposure, consistent with the lung type-17 and chemokine-positioning programme defined above, rather than delayed and sustained inflammatory cell accumulation.

Together, these data show that INSPIRE-i.n. vaccination imprinted a lung-localized type-17 and tissue-residency-associated transcriptional programme. The validated T-cell activation and type-17-responsiveness module, CXCR3–CXCL9/CXCL10 and CCL20-CCR6 chemokine-positioning axes, and residency-associated module provided molecular support for the IL-17⁺ T-cell subsets detected by intracellular cytokine staining and the IL-17A-secreting cells detected by ELISpot after INSPIRE-i.n. vaccination (**Fig. 4i, 4m-p**), and were accompanied by rapid, controlled airway myeloid recall after bacterial exposure. Because IL-17-producing CD8⁺ T cells are commonly described as a Tc17-like population, these transcriptional, spatial and cellular findings raised the possibility that INSPIRE-induced CD8⁺IL-17⁺ cells represent a functionally relevant Tc17-like component within the broader type-17 pulmonary protection programme, a hypothesis we next tested by functional perturbation.

### Tc17-like IL-17-dependent mucosal immunity protects against pulmonary *Klebsiella pneumoniae*

To determine whether antibody responses were required for INSPIRE-mediated protection, we first used polymeric immunoglobulin receptor-deficient mice (pIgR^−/−^), in which epithelial transport of polymeric immunoglobulins into mucosal secretions is impaired. In pIgR^−/−^ mice, INSPIRE-i.n. induced serum IgG responses, whereas BALF sIgA remained nearly undetectable. Despite the lack of airway sIgA, INSPIRE-i.n. markedly reduced pulmonary bacterial burdens after YBQ challenge compared with PBS-treated pIgR^−/−^ mice (**Extended Data Fig. 4a**). We next examined B-cell-deficient μMT mice to further assess the contribution of mature B-cell-dependent antibody responses. In μMT mice, INSPIRE-i.n. failed to induce detectable antigen-specific serum IgG, yet still significantly reduced pulmonary bacterial burdens after YBQ challenge (**Extended Data Fig. 4b**). These data indicate that neither airway sIgA nor mature B-cell-dependent antibody responses were strictly required for INSPIRE-mediated bacterial control.

We then examined whether protection required continuous recruitment of circulating lymphocytes at the time of challenge. Fingolimod (FTY720), a sphingosine-1-phosphate receptor modulator, markedly reduced circulating CD3⁺ T-cell frequencies (**Extended Data Fig. 4c, d**). Under FTY720-mediated lymphocyte-egress blockade, INSPIRE-i.n. maintained strong control of pulmonary bacterial burdens, with no significant loss of protection compared with INSPIRE-i.n. vaccination in the absence of FTY720 (**Extended Data Fig. 4d**). These results suggest that INSPIRE-mediated protection was maintained by a pre-established lung-localized immune compartment rather than by continuous recruitment of circulating lymphocytes during acute bacterial challenge.

To test whether the type-17-associated programme identified (**Fig. 4** and **Fig. 5**) was functionally required for protection, we used IL-17^−/−^ mice on the C57BL/6 background, together with background-matched wild-type C57BL/6 controls. In IL-17^−/−^ mice, INSPIRE-i.n. failed to reduce pulmonary bacterial burdens after YBQ challenge, with bacterial loads remaining comparable between PBS- and INSPIRE-i.n.-treated groups. By contrast, in background-matched wild-type C57BL/6 mice, INSPIRE-i.n. markedly reduced pulmonary bacterial burdens (**Fig. 6a**). Consistently, INSPIRE-i.n. significantly reduced IL-6, IL-1β and TNF-α levels in lung homogenate supernatants from wild-type mice after YBQ challenge. In IL-17^−/−^ mice, this inflammatory restraint was markedly impaired, with IL-1β and TNF-α no longer significantly reduced despite a partial reduction in IL-6 (**Fig. 6b-d**). Together with the bacterial-burden results, these data establish IL-17 as a required effector signal for INSPIRE-mediated bacterial control and post-challenge pulmonary inflammatory restraint.

**Fig. 6.**
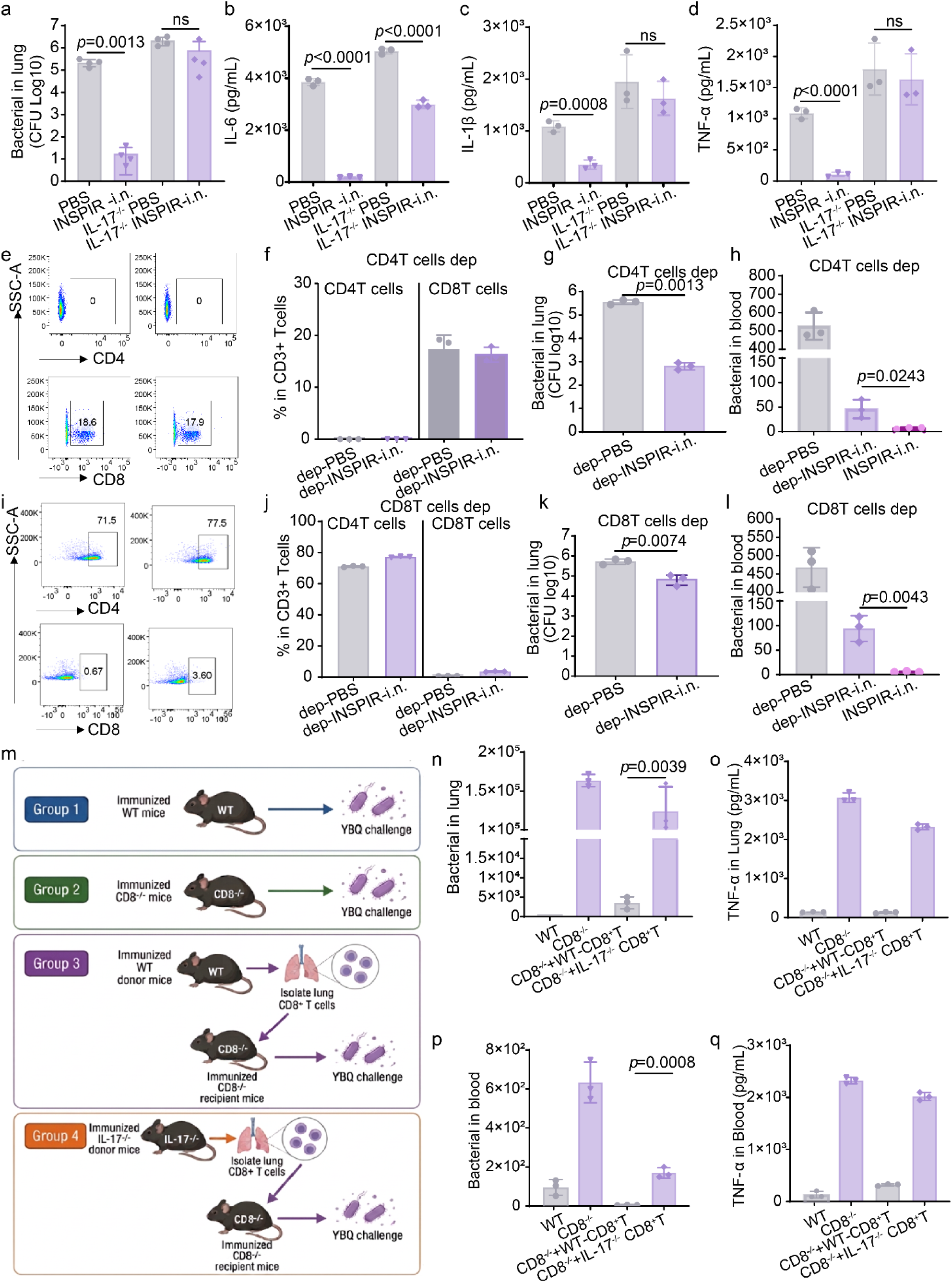
IL-17-dependent CD4⁺ and CD8⁺ T-cell immunity contributes to INSPIRE-mediated protection against pulmonary *Klebsiella pneumoniae* infection. **a–d,** Assessment of IL-17 involvement in INSPIRE-mediated protection. Wild-type and IL-17^−/−^ mice were intranasally immunized with PBS or INSPIRE and challenged intratracheally with *K. pneumoniae* YBQ. Pulmonary bacterial burdens **(a)** and interleukin-6 (IL-6) **(b)**, interleukin-1β (IL-1β) **(c)** and tumour necrosis factor-α (TNF-α) **(d)** levels in lung homogenate supernatants were determined after challenge. **e,f**, Representative flow-cytometry plots **(e)** and quantification **(f)** of CD4⁺ and CD8⁺ T cells among CD3⁺ T cells in lung single-cell suspensions after CD4⁺ T-cell depletion. **g,h**, Pulmonary bacterial burdens **(g)** and blood bacterial burdens **(h)** after YBQ challenge in CD4⁺ T-cell-depleted mice immunized with PBS or INSPIRE. Non-depleted INSPIRE-immunized mice were included as vaccinated controls in **h**. **i,j,** Representative flow-cytometry plots **(i)** and quantification **(j)** of CD4⁺ and CD8⁺ T cells among CD3⁺ T cells in lung single-cell suspensions after CD8⁺ T-cell depletion. **k,l**, Pulmonary bacterial burdens **(k)** and blood bacterial burdens **(l)** after YBQ challenge in CD8⁺ T-cell-depleted mice immunized with PBS or INSPIRE. Non-depleted INSPIRE-immunized mice were included as vaccinated controls in **l**. **m**, Schematic of the adoptive-transfer experiment. INSPIRE-immunized wild-type or IL-17^−/−^ donor mice were used to isolate lung CD8⁺ T cells, which were adoptively transferred into INSPIRE-immunized CD8^−/−^ recipient mice before YBQ challenge. INSPIRE-immunized wild-type and CD8^−/−^ mice without adoptive transfer served as controls. All mice were on the C57BL/6 background. **n–q**, Adoptive-transfer analysis of CD8⁺ T-cell-mediated protection after YBQ challenge. Pulmonary bacterial burdens **(n)**, TNF-α levels in lung homogenate supernatants **(o)**, blood bacterial burdens **(p)** and TNF-α levels in blood **(q)** were measured in INSPIRE-immunized wild-type mice, CD8^−/−^ mice, and CD8^−/−^ mice receiving lung CD8⁺ T cells from wild-type or IL-17^−/−^donors.

Having established the requirement for IL-17, we next asked which T-cell compartments mediated this protection. Antibody-mediated CD4⁺ T-cell depletion reduced CD4⁺ T cells to near-background levels while preserving the CD8⁺ T-cell compartment (**Fig. 6e, f**). After CD4⁺ T-cell depletion, INSPIRE-i.n. still reduced pulmonary bacterial burdens compared with depleted PBS controls, but bacterial loads remained higher than those in undepleted INSPIRE-immunized mice (**Fig. 6g**). A similar pattern was observed in blood, where CD4⁺ T-cell depletion increased systemic bacterial dissemination relative to undepleted INSPIRE-i.n. mice (**Fig. 6h**). These data indicate that CD4⁺ T cells contributed to optimal INSPIRE-mediated bacterial control but did not fully account for the protective effect.

We then performed CD8⁺ T-cell depletion to evaluate the contribution of the CD8⁺ compartment, which has been less extensively investigated in vaccine-mediated protection against extracellular bacterial pneumonia. CD8⁺ T cells were reduced to near-background levels, whereas CD4⁺ T cells were preserved (**Fig. 6i, j**). CD8⁺ T-cell depletion attenuated INSPIRE-mediated protection, increasing pulmonary bacterial burdens compared with undepleted INSPIRE-i.n. mice, although bacterial loads remained lower than in depleted PBS controls (**Fig. 6k**). CD8⁺ T-cell depletion also increased bacterial burdens in blood relative to undepleted INSPIRE-i.n. mice (**Fig. 6l**), indicating impaired containment of pulmonary infection and increased systemic dissemination. Together with the expansion of pulmonary CD8⁺IL-17⁺ T cells after INSPIRE-i.n. vaccination, these results implicate a Tc17-like CD8⁺ T-cell component in INSPIRE-mediated bacterial containment.

To directly examine whether CD8⁺ T-cell-derived IL-17 contributed to INSPIRE-mediated protection, we performed adoptive-transfer experiments in INSPIRE-immunized CD8-deficient recipient mice using CD8⁺ T cells isolated from INSPIRE-immunized wild-type or IL-17-deficient donor mice (**Fig. 6m**). Compared with CD8-deficient mice, transfer of wild-type CD8⁺ T cells markedly reduced bacterial burdens in both lung and blood after challenge (**Fig. 6n, p**). By contrast, transfer of IL-17-deficient CD8⁺ T cells provided significantly weaker protection than wild-type CD8⁺ T-cell transfer, resulting in higher bacterial burdens in both lung and blood (**Fig. 6n, p**). Consistently, wild-type CD8⁺ T-cell transfer reduced lung and blood inflammatory cytokine levels, whereas IL-17-deficient CD8⁺ T-cell transfer showed a weaker ability to restrain TNF-α responses (**Fig. 6o, q**). These data demonstrate that CD8⁺ T-cell-derived IL-17 is a functional contributor to INSPIRE-induced protection.

The proposed working model is summarized in **Extended Data Fig. 4e**. Intranasal INSPIRE promotes mRNA expression within pulmonary antigen-presenting-cell/dendritic-cell compartments and supports MHC-I- and MHC-II-associated antigen presentation, thereby engaging CD8⁺ and CD4⁺ T-cell responses. Although robust humoral responses were induced, perturbation experiments in pIgR^−/−^ and μMT mice showed that airway sIgA and mature B-cell-dependent antibody responses were not strictly required for bacterial control. Instead, protection was IL-17 dependent and involved both CD4⁺ and CD8⁺ T-cell compartments. Within this programme, INSPIRE-expanded CD8⁺IL-17⁺ T cells constitute a Tc17-like component that contributes to pulmonary bacterial containment, limits bloodstream dissemination and supports vaccine-mediated defence against extracellular Kp.

## Conclusions

In summary, this study pioneers an investigation of mRNA vaccine against *Klebsiella pneumoniae*-a WHO priority pathogen that urgently demands vaccine development. Through systematic donor-cell screening, we identified mature dendritic cell-derived exosomes as a privileged mRNA delivery scaffold that naturally couples miR-21-mediated epithelial barrier modulation with miR-155-driven DC activation via SOCS1/Inpp5d relief. To overcome inherent exosomal limitations, an ML-designed peptide was engineered to boost mRNA encapsulation and markedly enhances antigen cross-presentation, yielding INSPIRE. Functionally, INSPIRE mediated Kp-mRNA vaccination induces durable, lung-resident CD8⁺IL-17⁺ T cells that rapidly recruit neutrophils and macrophages, achieve near-complete cross-strain protection against lethal, drug-resistant Kp, and abrogate bacterial dissemination. Mechanistically, using pIgR^−/−^, IL-17^−/−^ and T-cell depletion models, we demonstrate that protection does not require serum related immunity but depends mainly on an IL-17 dependent CD4⁺/CD8⁺ T cell network-with CD8⁺Tc17 cells playing essential, previously unappreciated roles against an extracellular pathogen such as Kp.

Collectively, the data presented here establishes a new framework of mucosal vaccination against extracellular bacterial pneumonia by integrating three previously disconnected elements: a biologically active exosome backbone, machine-learning-guided peptide engineering, and a non-canonical T-cell protective correlate. This blueprint might reframe how we think about vaccine development strategy of Gram-negative bacillus and shift the paradigm from antibody-centric dogma to Tc17-prioritized design principle.

## Methods

### Reagents

Nuclease-free water was purchased from QIAGEN. Opti-MEM I reduced-serum medium, LysoTracker Red DND-99, DiR fluorescent dyes, protein transport inhibitor cocktail and lithium chloride were obtained from Thermo Fisher Scientific. Agarose was purchased from Aladdin. MOPS buffer was purchased from Solarbio. Cell Counting Kit-8 reagent was purchased from MedChemExpress. Quant-iT RiboGreen RNA reagent was obtained from Thermo Fisher Scientific. D-luciferin potassium salt and other routine chemical reagents were obtained at analytical or HPLC grade from Sigma-Aldrich (Shanghai, China), unless otherwise stated. Antibodies used for flow cytometry, and depletion experiments are listed in **Supplementary Table 5**.

### Cell culture

Calu-3, BEAS-2B, A549, RAW264.7 and HEK293T cells were cultured at 37 °C in a humidified incubator containing 5% CO₂. Calu-3 cells were used to form epithelial monolayers for the transwell-based epithelial– immune screening assay. Exosome donor cells used in Fig. 1 included BEAS-2B, A549, RAW264.7, HEK293T, umbilical cord-derived mesenchymal stem cells (UMSCs), immature bone marrow-derived dendritic cells (I-BMDCs) and mature BMDCs (M-BMDCs). BEAS-2B and HEK293T cells were maintained in RPMI-1640 medium supplemented with 10% fetal bovine serum, 100 U ml⁻¹ penicillin and 100 μg ml⁻¹ streptomycin. A549, Calu-3 and RAW264.7 cells were maintained in Dulbecco’s modified Eagle medium (DMEM) containing the same supplements. UMSCs were cultured in low-glucose Dulbecco’s modified Eagle medium (DMEM-LG) supplemented with 10% fetal bovine serum, 100 U ml⁻¹ penicillin and 100 μg ml⁻¹ streptomycin. Cells were passaged every 2-3 d and used during logarithmic growth. All cell lines tested negative for mycoplasma contamination. I-BMDCs and M-BMDCs were generated from mouse bone marrow cells as described below.

### Bone marrow-derived dendritic cells and macrophages

Bone marrow-derived dendritic cells were prepared from 6-8-week-old female BALB/c mice. Bone marrow cells were flushed from femurs and tibias, filtered through a 70 μm cell strainer and cultured in complete RPMI-1640 medium supplemented with granulocyte-macrophage colony-stimulating factor (GM-CSF, 20 ng ml⁻¹) and interleukin-4 (IL-4, 10 ng ml⁻¹). Fresh cytokine-containing medium was replenished every 2 days. Immature BMDCs were collected on day 7. Mature BMDCs were generated by stimulation with lipopolysaccharide (LPS, 100 ng ml⁻¹) for 24 h before use or exosome collection. For exosome preparation, cells were washed and cultured in medium containing exosome-depleted FBS for 24-48 h, and culture supernatants were collected for exosome isolation. Bone marrow-derived macrophages were generated by culturing bone marrow cells with macrophage colony-stimulating factor (M-CSF, 20 ng ml⁻¹) for 7 days.

### Animals

All animal experiments were approved by the Laboratory Animal Welfare and Ethics Committee of Third Military Medical University under approval number [AMUWEC20230184] and followed institutional guidelines for animal care and use. Female BALB/c and C57BL/6 mice aged 6-8 weeks were obtained from Beijing Vital River Laboratories. pIgR^−/−^ mice and IL-17^−/−^ mice on a C57BL/6 background were kindly provided by investigators at Third Military Medical University. Age- and sex-matched C57BL/6 mice were used as wild-type controls for knockout-mouse experiments. Rosa26-LSL-tdTomato reporter mice were purchased from Cyagen Biosciences (Guangzhou, China). Animals were housed under specific pathogen-free conditions with controlled temperature, humidity and a 12 h light/dark cycle. Sterile food and water were provided ad libitum. Mice were acclimatized for at least 7 d before experiments and randomly assigned to treatment groups.

### Bacterial strains

Three *Klebsiella pneumoniae* strains were used for challenge studies. YBQ is a K20 clinical isolate preserved in our laboratory and originally isolated from the First Affiliated Southwest Hospital of Army Medical University. YYD is a K1 isolate from the same hospital. A7818 is a K2 isolate obtained from Guangxi Medical University. Bacteria were cultured in Luria-Bertani medium at 37 °C with shaking until logarithmic phase. Cultures were washed with sterile PBS and diluted to the required challenge dose. The final inoculum was confirmed by serial dilution and colony counting.

### In vitro-transcribed messenger RNA (IVT-mRNA) synthesis

IVT-mRNAs were synthesized and purified as previously described. ^48^ Briefly, IVT-mRNA encoding the indicated synthetic design sequences (SDS; sequences are provided in Supplementary Table 3) was synthesized by from linearized DNA templates using a T7 RNA transcription kit (E131-01A, Novoprotein, Shanghai, China). N1-methylpseudouridine (Glycogene, Wuhan, China) was incorporated during transcription, and a Cap 1 structure was added after transcription. The mRNAs used in this study included mFluc, mEGFP, mCre and Kp-mRNA. Kp-mRNA encoded a KPC-2–linker–Pal fusion antigen, and the sequence information is provided in Supplementary Table1. IVT-mRNA was purified by lithium chloride precipitation or chromatography-based purification according to the experimental requirement. RNA concentration and purity were measured with a NanoDrop One spectrophotometer, and RNA integrity was assessed by agarose gel electrophoresis or Bioanalyzer analysis when required.

### Isolation of donor-cell-derived exosomes

Donor cells were cultured in medium containing exosome-depleted fetal bovine serum before exosome collection. Culture supernatants were collected after 48 h and cleared by sequential centrifugation to remove cells, debris and larger vesicles. Briefly, supernatants were centrifuged at 300g for 10 min, 2,000g for 20 min and 10,000g for 30 min at 4 °C, followed by filtration through 0.22 μm filters. Exosomes were collected by ultracentrifugation at 100,000g for 60 min, washed with sterile PBS and centrifuged again under the same conditions. The final exosome pellets were resuspended in PBS and stored at −80 °C.

Exosomes were prepared from multiple donor-cell sources for the initial screening, including epithelial cells, macrophage-lineage cells, immature BMDCs, mature BMDCs and BMDMs. Mature BMDC-derived exosomes were selected for INSPIRE construction on the basis of their combined transepithelial transport and BMDC activation activities.

### Characterization of exosomes and INSPIRE

Particle size, polydispersity index and zeta potential were measured by dynamic light scattering at 25 °C. Particle morphology was examined by transmission electron microscopy after negative staining. Exosome marker proteins, including CD63, TSG101 and Alix, were analysed by Western blotting, with calnexin used as a negative cellular-contaminant marker where appropriate. Protein concentration was determined by BCA assay. IVT-mRNA encapsulation efficiency was quantified using the RiboGreen RNA assay as described below.

### Screening of bioactive exosome backbones

Calu-3 epithelial monolayers were established on PET transwell inserts for 21 d. Transepithelial electrical resistance was measured before and after treatment with donor-cell-derived exosomes. For transport and immune-activation screening, exosomes were added to the apical chamber, and BMDCs were seeded in the basolateral compartment. After incubation, BMDCs were collected for flow-cytometric analysis of maturation markers. BMDC activation was defined by the frequencies of CD80⁺MHC-II⁺ and CD86⁺MHC-II⁺ cells. TEER recovery was monitored to distinguish reversible epithelial barrier modulation from sustained epithelial disruption.

### Public miRNA-Seq Data Mining

Exosomal miRNA-sequencing profiles were retrieved from the NCBI Gene Expression Omnibus (GEO) under accession GSE190854. After abundance and variance filtering, differential miRNA analysis was performed using DESeq2 with thresholds of adjusted *p*<0.05 and ∣log_2_FC ∣>1. Candidate miRNAs were further prioritized by functional relevance (miRDB, TargetScan). Heatmaps were generated with z-score-normalized expression values.

### RNA-Seq and Differential Expression Analysis

Total RNA from lung tissues was sequenced on an Illumina platform. Reads were aligned to the mouse genome (GRCm38) using HISAT2 and quantified with Feature Counts. Differential expression analysis (DESeq2) was applied with *p* < 0.05 and ∣log_2_FC∣>1. INSPIRE-associated genes were defined as those altered in INSPIRE vs. Exo and concoprdantly altered in INSPIRE vs. PBS, or uniquely altered in INSPIRE vs. Exo. GO and KEGG enrichment analyses were conducted using cluster Profiler (adjusted *p*<0.05).

### qRT-PCR Validation

Relative expression of *Socs1* and *Inpp5d* was measured by qRT-PCR using gene-specific primers and SYBR Green, normalized to *Gapdh* via the 2^−ΔΔCt^method. Using R packages (ggplot2, pheatmap, circlize). Data are presented as mean ±SD. Two-group comparisons used two-tailed unpaired Student’s t-tests.

### Single-Particle Analysis of Exosome Cargo

Exosomes and INSPIRE were labeled with specific fluorescent probes. Cy5-labeled CD63 antibodies were used to identify CD63-positive vesicles, and MFP488-labeled mRNA probes were employed to detect mRNA-associated particles. Single-particle flow analysis was conducted using a flow cytometer equipped for nanoscale particle detection (e.g., Apogee A50 Micro, Apogee Flow Systems). Particle populations were defined as CD63⁺mRNA⁺ (P1), CD63⁺mRNA⁻ (P2), CD63⁻mRNA⁻ (P3), and CD63⁻mRNA⁺ (P4) based on fluorescence intensity thresholds.

### mRNA Stability Assay

The stability of encapsulated mRNA in Exo and INSPIRE was assessed over time. Exosome formulations containing mRNA were incubated at 37°C for various durations. At specified time points, samples were collected, and mRNA was extracted. The integrity of mRNA was then evaluated by agarose gel electrophoresis and quantified densitometrically. The degradation rates of mRNA were calculated based on the reduction in band intensity over time.

### Exosomal miRNA analysis

Total RNA was extracted from exosomes using a miRNA-compatible RNA isolation kit. miR-21 and miR-155 abundance was measured by RT–qPCR. For functional validation, anti-miR-21 or anti-miR-155 oligonucleotides were used to reduce the corresponding miRNA activity. miR-21-associated epithelial effects were examined in 16HBE cells, whereas miR-155-associated DC maturation effects were examined in BMDCs or immature BMDC-derived exosome conditions.

### Scratch-wound assay and epithelial junction staining

16HBE cells were grown to confluence in culture plates. Linear scratches were generated with sterile pipette tips, and detached cells were removed by PBS washing. Cells were then treated with mature BMDC-derived exosomes, miRNA-modified exosomes or control oligonucleotides. Images were collected at the indicated time points, and wound closure was quantified using ImageJ.

For junctional protein staining, cells were fixed with 4% paraformaldehyde, permeabilized with 0.1% Triton X-100 and blocked with BSA. Primary antibodies against ZO-1, occludin and E-cadherin were applied overnight at 4 °C, followed by fluorophore-conjugated secondary antibodies. Nuclei were stained with DAPI, and images were acquired by confocal microscopy using identical settings across groups.

### Machine Learning-Guided Lipopeptide Design, Virtual Screening, and Combinatorial SAR Profiling

A tripartite peptide library comprising 200 rational sequences was constructed by combinatorially coupling 20 DC-targeting A-modules (A-A20) with 10 mRNA-condensing and pH-responsive B-modules (B1–B10), flanked by a conserved C-terminal anchoring motif (-GG-C) for subsequent DSPC– PEG – NHS lipid conjugation. For each full-length peptide, nine biophysical and sequence descriptors were quantified: residue counts for His (endosomal proton-sponge escape), Glu/Asp, and Lys/Arg (cationic mRNA complexation); total sequence length (26 – 39 residues); theoretical net charges at physiological pH 7.4 (−7.0 to +7.8) and endosomal pH 5.5 (−0.6 to +17.6); charge-flipping ratio (ΔQ, −0.1 to 66.5); grand average of hydropathicity (GRAVY index, −0.69 to +0.23); and predicted alpha-helix propensity (0.26 to 0.84).

Experimentally measured in vitro mRNA transfection efficiency (28.63%–91.81%), primary BMDC activation rate (37.30%–78.23%, determined by flow cytometry for CD80/CD86/MHC expression), and cell proliferation inhibition rate (28.55%–73.96%, measured by MTT assay) served as multi-task target vectors. Five supervised regression architectures—k-nearest neighbors (KNN), decision tree (DT), random forest (RF), gradient boosting regressor (GBoost), and extreme gradient boosting (XGBoost) — were trained and systematically benchmarked. To prevent data leakage, missing values were imputed using training set medians, and descriptors were standardized via Z-score normalization strictly within each training fold. The dataset (200 peptides) was partitioned into an 80% training set (n = 160) and a 20% independent hold-out test set (n = 40) using 5-fold cross-validation (random seed = 42). Model predictive precision and generalization were evaluated using the coefficient of determination (R²) and root-mean-square error (RMSE), while multi-task robustness across biophysical and immunological dimensions was profiled via pentagonal radar mapping.

Among the evaluated algorithms, ensemble boosting models demonstrated superior predictive fidelity (test R^2^ = 0.774), exhibiting tight adherence to the parity regression line within the 95% confidence interval. Feature contribution and TreeSHAP analyses identified the charge-flipping ratio, histidine content, and GRAVY index as the primary determinants governing delivery performance.

The optimized ensemble model was subsequently deployed to screen an in silico high-throughput virtual library of 20,000 candidate peptides across the continuous biophysical design space (sequence length vs. GRAVY index). Model predictions indicated that high-performance candidates clustered at sequence lengths of 36–38 residues and GRAVY values between −0.4 and −0.2. In parallel, combinatorial structure-activity relationship (SAR) bubble matrix profiling uncovered a strong cooperative synergy between high histidine counts (His >= 4) and moderate cationic charges (Lys/Arg = 5–8) for maximizing transfection potency.

Ten top-ranked candidates from this high-scoring cluster were synthesized (>95% purity by RP-HPLC) and validated in BMDC cultures. Peptide A7-B2 (FSRSLHSLL-GPTI-GG-WEAHLAHALAHALAHHLAHAL-C), which combines a DCIR-targeting A7 module with a histidine-enriched B2 module, yielded the highest combined mRNA transfection efficiency (91.81%) and BMDC activation rate (78.23%), and was selected for exosome surface engineering.

### Preparation and Surface Engineering of Peptide-Modified Exosomes (INSPIRE)

INSPIRE nanoparticles were fabricated by engineering mature BMDC-derived exosomes with the machine-learning-prioritized lead peptide (A7-B2). The peptide, containing both the dendritic-cell-engaging domain and the pH-responsive/mRNA-condensing domain, was lipid-anchored onto the exosomal lipid bilayer via DSPC–PEG–NHS coupling. Briefly, peptide A7-B2 was reacted with DSPC–PEG–NHS in PBS (pH 7.4–8.0) under gentle agitation at room temperature to facilitate NHS ester-mediated coupling with accessible primary amines on the peptide. The resulting DSPC– PEG– peptide conjugate was purified to remove unreacted free peptide and excess lipid crosslinkers.

Purified mature BMDC-derived exosomes were subsequently incubated with the DSPC–PEG–peptide conjugate under gentle mixing, driving spontaneous hydrophobic insertion of the DSPC lipid anchors into the exosomal membrane. Unincorporated conjugates and free reagents were removed by ultrafiltration using a 100 kDa molecular weight cut-off centrifugal filter. The concentrated retentate was washed three times with sterile PBS and resuspended to obtain the final peptide-engineered exosome formulation (INSPIRE).

### IVT-mRNA loading into Exo and INSPIRE

IVT-mRNA was loaded into Exo or INSPIRE by electroporation. Briefly, purified exosomes were diluted in Gene Pulser electroporation buffer and mixed with IVT-mRNA at the optimized particle-to-mRNA ratio. The mixture was transferred into a 4 mm electroporation cuvette and precooled on ice. Electroporation was performed using a Gene Pulser Xcell system under square-wave conditions. Based on the optimized exosome-loading protocol, the pulse parameters were set as 200 V, 10 ms, five pulses and a 1 s interval between pulses. After electroporation, samples were incubated on ice for recovery and then concentrated by ultrafiltration using a 100 kDa cut-off centrifugal filter. Free mRNA and electroporation buffer were removed by repeated washing with sterile PBS. The final formulations were freshly used or stored at 4 °C for short-term experiments.

### Characterization of Exo and INSPIRE

The hydrodynamic diameter, polydispersity index and zeta potential of Exo and INSPIRE were measured by dynamic light scattering at 25 °C. Particle concentration was determined by nanoparticle tracking analysis. Morphology was examined by transmission electron microscopy. For TEM imaging, samples were placed on copper grids, negatively stained with 2% phosphotungstic acid or uranyl acetate for 1 min, dried at room temperature and imaged under a transmission electron microscope.

Exosomal marker proteins were analysed by western blotting. Equal amounts of exosomal protein were separated by SDS–PAGE and transferred onto PVDF membranes. Membranes were blocked with 5% BSA or skim milk, incubated overnight at 4 °C with primary antibodies against CD63, TSG101 and Alix, and then incubated with HRP-conjugated secondary antibodies. Calnexin was used as a negative cellular-contaminant marker where appropriate. Protein bands were visualized using chemiluminescence.

### IVT-mRNA encapsulation efficiency

The encapsulation efficiency of IVT-mRNA in Exo or INSPIRE was measured using the Quant-iT RiboGreen RNA assay. Formulations were divided into intact and lysed groups. For the lysed group, vesicles were disrupted with 2% Triton X-100 and incubated at 37 °C for 30 min. RNA standards were prepared in TE buffer. Each sample or standard was mixed with diluted RiboGreen reagent in a black 96-well plate and incubated at room temperature for 3 min. Fluorescence was measured at excitation/emission wavelengths of 480/520 nm. Encapsulation efficiency was calculated according to the following equation:

Encapsulation efficiency (%) = (total mRNA − free mRNA) / total mRNA × 100.

### Stability of mRNA-loaded INSPIRE against nuclease digestion

To assess mRNA protection, free IVT-mRNA, Exo-loaded mRNA and INSPIRE-loaded mRNA were incubated with RNase I at 37 °C for the indicated time points. Vesicles were then collected by ultracentrifugation or ultrafiltration, lysed with Triton X-100 to release encapsulated mRNA and analysed by agarose gel electrophoresis. Free IVT-mRNA exposed to RNase I served as the degradation control. Intact mRNA bands after vesicle lysis indicated protection of IVT-mRNA by Exo or INSPIRE.

### Fluorescent labelling of Exo and INSPIRE

For biodistribution and cellular uptake assays, Exo or INSPIRE was labelled with DiR. Fluorescent dye was added to the exosome suspension at the indicated concentration and incubated at 37 °C for 2 h. Unbound dye was removed by ultrafiltration with a 100 kDa cut-off centrifugal filter. Labelled vesicles were washed with PBS before use. Fluorescence-labelled formulations were freshly prepared for cellular uptake, tissue distribution and in vivo imaging experiments.

### In vitro mRNA uptake and transfection

For uptake experiments, cells were seeded in 24-well plates and cultured overnight. Cy5-mRNA-loaded Exo or INSPIRE was added to cells at the indicated mRNA dose. After 6 h, cells were washed with PBS, detached and analysed by flow cytometry. Uptake was quantified as the percentage of Cy5-positive cells and mean fluorescence intensity.

IVT-mRNA transfection assays were performed as previously described.^49,50^ Briefly, DC2.4 cells were seeded in 96-well plates 24 h before treatment. Exo or INSPIRE loaded with mFluc or mEGFP was added to the cells and incubated for 4-6 h before replacement with fresh complete medium. Luciferase activity was measured 24 h later using a firefly luciferase assay kit. EGFP expression was assessed by fluorescence microscopy and flow cytometry.

#### Uptake-inhibition assay

DC2.4 cells were pretreated with endocytosis inhibitors before addition of Cy5-mRNA-loaded INSPIRE. Sodium azide was used for energy depletion, amiloride for macropinocytosis inhibition, protamine for charge-associated membrane interaction competition, chlorpromazine for clathrin-mediated endocytosis inhibition and filipin for lipid-raft/caveolae-associated uptake inhibition. For low-temperature treatment, cells were prechilled and maintained at 4 °C. After inhibitor pretreatment, INSPIRE was added and incubated for 6 h. Relative uptake was calculated by normalizing fluorescence signals to untreated INSPIRE-treated controls.

### Endo/lysosomal release imaging

DC2.4 cells were seeded in confocal dishes and treated with Exo or INSPIRE loaded with XFD488-labelled mRNA. After incubation, cells were stained with LysoTracker Red DND-99 and counterstained with DAPI. Confocal images were acquired using identical settings. Co-localization between mRNA and lysosomal signals was quantified using ImageJ or equivalent software.

### Cell viability assay

Cell viability was measured using the Cell Counting Kit-8 assay. DC2.4 cells were seeded in 96-well plates and treated with PBS, Exo or INSPIRE formulations for 24 h. CCK-8 reagent was added to each well and incubated at 37 °C for 2 h. Absorbance was measured using a microplate reader. Blank wells containing medium and reagent but no cells were used for background correction. Viability was calculated relative to PBS-treated controls.

### Ussing chamber assay

Rat tracheal or bronchial epithelial tissues were excised and mounted between donor and receptor chambers of an Ussing chamber system. The receptor chamber was filled with HEPES-buffered solution, and Exo or INSPIRE loaded with Cy5-mRNA was added to the donor chamber. Samples were collected from the receptor chamber at the indicated time points and replaced with fresh buffer. Fluorescence intensity was measured using a microplate reader. Apparent permeability coefficients were calculated from transported mRNA amount, epithelial surface area, donor concentration and incubation time.

### In vivo biodistribution

For biodistribution analysis, Exo and INSPIRE were labelled with DiR. BALB/c mice were lightly anaesthetized with isoflurane and intranasally administered labelled formulations. Whole-body fluorescence imaging was performed at the indicated time points using an IVIS imaging system. At selected endpoints, mice were euthanized and lungs or major organs were collected for ex vivo imaging. Fluorescence signals were quantified using Living Image software.

### In vivo mRNA expression

BALB/c mice received intranasal administration of Exo or INSPIRE loaded with mFluc mRNA. At 6, 24, 48 and 72 h after administration, mice were anaesthetized and injected intraperitoneally with D-luciferin. Whole-body bioluminescence imaging was performed using an IVIS imaging system. Lungs were collected for ex vivo imaging where indicated. Total flux was quantified with Living Image software.

### In vivo tracing of mRNA-transfected cells

Rosa26-LSL-tdTomato reporter mice were intranasally treated with Exo or INSPIRE loaded with Cre recombinase mRNA. Three days later, lungs and mediastinal lymph nodes were collected. Tissues were fixed, cryoprotected, embedded in OCT and sectioned. Sections were stained for CD11c and CD86, and nuclei were counterstained with DAPI. tdTomato fluorescence was used as the readout of Cre-mediated recombination.

For flow-cytometric quantification, lungs were perfused with PBS, minced and digested with collagenase II and DNase I. Single-cell suspensions were filtered, erythrocytes were lysed and cells were stained with antibodies for dendritic-cell and macrophage identification. tdTomato-positive dendritic cells and macrophages were quantified by flow cytometry.

### Safety evaluation after intranasal administration

BALB/c mice were intranasally treated with PBS, Exo or INSPIRE. BALF, serum and major organs were collected at the indicated time points. IL-6, IL-1β and TNF-α levels in BALF and serum were quantified by ELISA. Major organs, including lung, liver, spleen, brain, kidney and heart, were fixed in 4% paraformaldehyde, embedded in paraffin, sectioned and stained with haematoxylin and eosin. Serum biochemical markers related to liver, kidney and tissue injury were measured using an automated biochemical analyser or by a commercial testing service.

### Immunization

BALB/c mice were immunized with Kp-mRNA formulated with INSPIRE or unmodified Exo by intranasal administration. LNP-formulated Kp-mRNA delivered intramuscularly was used as a comparator, and PBS-treated mice served as controls. Mice were primed on day 0 and boosted on day 21. Unless otherwise specified, the mRNA dose in Exo and INSPIRE formulations was 2 μg per mouse. For intranasal delivery, mice were lightly anaesthetized with isoflurane and formulations were applied dropwise to the nostrils. For intramuscular immunization, formulations were injected into the hindlimb muscle.

### Collection of serum, BALF and NALF

Serum was collected from peripheral blood after clotting and centrifugation. BALF was obtained by tracheal cannulation and gentle lavage with sterile PBS. NALF was collected by flushing the nasal cavity with sterile PBS. Samples were clarified by centrifugation and stored at −80 °C until use.

### ELISA for antigen-specific antibodies

Kp-antigen-specific IgG and sIgA were measured by ELISA. Plates were coated with recombinant Kp protein overnight at 4 °C and blocked before sample addition. Serially diluted serum, BALF or NALF samples were incubated in the coated plates, followed by HRP-conjugated anti-mouse IgG or anti-mouse IgA detection antibodies. TMB substrate was used for color development, and absorbance was measured at 450 nm. Endpoint titres were defined as the highest dilution giving a signal above the cut-off value.

### IL-17A ELISpot

IL-17A-secreting cells were measured using a mouse IL-17A ELISpot kit. Pulmonary lymphocytes or splenocytes were seeded into antibody-coated plates and restimulated with Kp-antigen antigen or peptide pools for 12 h. Plates were developed according to the kit protocol. Spots were counted using an ELISpot reader and expressed as spot-forming cells per input cell number.

### Opsonophagocytic killing assay

HL-60 cells were differentiated into granulocyte-like cells in RPMI-1640 medium containing10% FBS, 1% glutamine and 0.8% dimethylformamide. Differentiated cells were collected by centrifugation at 1,000 rpm for 5 min, washed with HBSS with and without Ca²⁺/Mg²⁺, and resuspended in opsonophagocytic killing buffer at 1 × 10⁷ cells ml⁻¹. Log-phase Kp YBQ was washed and diluted in opsonophagocytic killing buffer to 2.5 × 10⁵ CFU ml⁻¹. In each well of a 96-well plate, 20 μl of diluted serum or BALF sample was mixed with complement and 10 μl of bacterial suspension. Control wells contained buffer alone or heat-inactivated complement. After shaking for 1 h at room temperature, 50 μl of the HL-60 cell-complement mixture was added to each well, and plates were incubated for another 1 h at 37 °C with 5% CO₂ under shaking. Reactions were stopped on ice for 20 min. Samples were serially diluted, plated on LB agar and incubated overnight at 37 °C for colony counting. Killing activity was calculated relative to wells lacking immune serum or BALF.

### Pulmonary Kp challenge

Immunized mice were anaesthetized and challenged intratracheally with Kp Acute infection was induced using YBQ or YYD at 1 × 10⁷ CFU per mouse, or A7818 at 1 × 10^6^ CFU per mouse. Body weight, survival and global disease scores were recorded daily for 7 d after challenge. Mice reaching humane endpoints were euthanized and counted as non-survivors.

### Bacterial burden analysis

For the pulmonary Kp YBQ challenge model, mice were intratracheally inoculated with 1 × 10⁶ CFU of YBQ per mouse. The inoculum dose was confirmed by serial dilution plating. At the indicated time points after challenge, lungs, spleens and blood were collected aseptically. Solid organs were weighed and homogenized in sterile PBS. Tissue homogenates and blood samples were serially diluted and plated on LB agar plates. Plates were incubated overnight at 37 °C, and colonies were counted the next day. Bacterial burdens were expressed as CFU per gram of tissue for solid organs and CFU per mL for blood samples.

### Histology after bacterial challenge

Lungs were collected at 2 d after challenge and fixed in 4% paraformaldehyde. Paraffin-embedded sections were stained with haematoxylin and eosin. Whole-lung sections and magnified regions were imaged by bright-field microscopy. Lung inflammation, consolidation and alveolar architecture were evaluated from stained sections.

### Cytokine quantification in lung homogenates

Lung tissues were homogenized in PBS containing protease inhibitors and centrifuged at 12,000g for 15 min at 4 °C. Supernatants were collected for ELISA analysis. IL-6, IL-1β and TNF-α concentrations were measured using commercial ELISA kits. Cytokine levels were calculated from standard curves and normalized as indicated.

### Bulk RNA sequencing

Lung tissues were collected from PBS-, Exo-i.n.- and INSPIRE-i.n.-immunized mice at the indicated time point. Total RNA was isolated with DNase I treatment. RNA integrity was assessed using an Agilent Bioanalyzer, and samples with RNA integrity number ≥8.0 were used for library construction. Stranded mRNA-seq libraries were prepared and sequenced on an Illumina platform with paired-end reads.

Raw reads were filtered to remove adapters and low-quality sequences, aligned to the mouse reference genome and quantified at the gene level. Differential expression analysis was performed using DESeq2. INSPIRE-associated genes were defined by retaining INSPIRE-vs-Exo-specific genes and genes shared between INSPIRE-vs-Exo and INSPIRE-vs-PBS comparisons, while excluding Exo-associated background changes. Gene Ontology and Kyoto Encyclopedia of Genes and Genomes enrichment analyses were performed for functional annotation. Volcano plots, Venn diagrams, heatmaps and circular heatmaps were generated using R packages.

### RT-qPCR validation of lung transcriptional modules

Total RNA from lung tissues was reverse-transcribed into cDNA. Quantitative PCR was performed using SYBR Green master mix and gene-specific primers. Gene expression was normalized to *Gapdh* and calculated by the 2^⁻ΔΔCt^ method. Primer sequences are provided in **Supplementary Table 3**. The analysed genes included IL-17/T-cell activation-related genes *Irf4*, *Icos*, *Il23r*, *Il21r*, *Il12rb1*, *Cd3e*, Cd3g and *Zap70*; chemokine-positioning genes *Cxcr3, Cxcl9, Cxcl10, Ccl20 and Ccr6*; and T-cell residency-associated genes *Cd69*, *Itgae* and *Cxcr6*.

### BALF myeloid-cell analysis after sublethal challenge

For airway recall analysis, immunized mice were challenged intratracheally with a sublethal dose of YBQ. BALF was collected at 0 h, 6 h, 12 h and 48 h after challenge. BALF cells were pelleted, washed and stained with antibodies against CD45, CD11b, Ly6G and F4/80. Neutrophils were defined as CD45⁺CD11b⁺Ly6G⁺ cells, and macrophages were defined as CD45⁺CD11b⁺F4/80⁺ cells.

### pIgR^−/−^ mouse experiment

pIgR^−/−^ mice were immunized intranasally with PBS or INSPIRE formulated with Kp-mRNA using the prime-boost schedule described above. Serum and BALF were collected to measure Kp-antigen-specific IgG and sIgA. Mice were challenged intratracheally with YBQ, and lung bacterial burdens were quantified 48 h after challenge.

### Fingolimod treatment

Fingolimod was used to restrict lymphocyte egress and limit the recruitment of circulating lymphocytes to the lung. Immunized mice received fingolimod intraperitoneally at 1 mg/kg once daily, starting 7 d before challenge and continuing until sample collection. Control mice received sterile water. Circulating CD3⁺ T-cell availability was assessed in peripheral blood by flow cytometry before bacterial challenge. Mice were then challenged intratracheally with YBQ, and lung bacterial burdens were measured 48 h after challenge.

### IL-17^−/−^ mouse experiment

Wild-type and IL-17^−/−^ mice were immunized intranasally with PBS or INSPIRE formulated with Kp-mRNA. After YBQ challenge, lungs were collected for bacterial burden analysis. Lung homogenate supernatants were used to measure IL-6, IL-1β and TNF-α by ELISA.

### CD4⁺ and CD8⁺ T-cell depletion

For T-cell depletion, immunized mice received intraperitoneal injections of anti-CD4 or anti-CD8β depleting monoclonal antibodies before bacterial challenge. Anti-CD4 antibody (clone GK1.5) or anti-CD8β antibody (clone Lyt3.2) was administered at 200 μg per mouse on days −3, −1 and +1 relative to challenge. Control mice received the corresponding isotype control antibodies. Depletion efficiency was confirmed in lung single-cell suspensions by quantifying CD4⁺ or CD8α⁺ cells among live CD3⁺ T cells. Mice were challenged intratracheally with YBQ, and lung bacterial burdens were measured 48 h after challenge.

### CD8⁺ T-cell adoptive-transfer experiment

To assess the contribution of CD8⁺ T-cell-derived IL-17 to INSPIRE-mediated protection, adoptive-transfer experiments were performed using CD8-deficient recipient mice on the C57BL/6 background. Donor CD8⁺ T cells were isolated from age- and sex-matched wild-type C57BL/6 or IL-17^−/−^ mice. Donor mice were immunized intranasally with INSPIRE formulated with Kp-mRNA using the same prime-boost schedule as described above. At the indicated time point after booster immunization, spleens, mediastinal lymph nodes and/or lung tissues were collected from donor mice and processed into single-cell suspensions. Lung tissues were perfused with cold PBS, minced and digested with collagenase II and DNase I before filtration through 70 μm cell strainers.

CD8⁺ T cells were purified from donor single-cell suspensions using a mouse CD8⁺ T-cell isolation kit or by fluorescence-activated cell sorting. Purified cells were analysed by flow cytometry to confirm CD3⁺CD8⁺ T-cell purity and viability before transfer. Equal numbers of wild-type or IL-17^−/−^ CD8⁺ T cells were resuspended in sterile PBS and transferred intravenously into CD8-deficient recipient mice. Recipient mice were intranasally immunized with INSPIRE formulated with Kp-mRNA according to the standard prime-boost regimen. Mice were assigned to the following groups: wild-type mice, CD8-deficient mice, CD8-deficient mice receiving wild-type CD8⁺ T cells and CD8-deficient mice receiving IL-17^−/−^ CD8⁺ T cells.

After adoptive transfer, recipient mice were challenged intratracheally with Klebsiella pneumoniae YBQ. Lungs and blood were collected 48 h after challenge. Pulmonary and blood bacterial burdens were determined by serial dilution and colony counting. Lung homogenate supernatants and serum or plasma samples were collected for measurement of TNF-α by ELISA. The protective capacity of wild-type and IL-17^−/−^ CD8⁺ T cells was compared to determine the functional contribution of CD8⁺ T-cell-derived IL-17 to INSPIRE-mediated bacterial control.

### Flow cytometry

Single-cell suspensions were prepared from lung tissues, spleens, bronchoalveolar lavage fluid (BALF) or cultured cells according to the requirements of each experiment. For lung samples, mice were perfused with cold PBS before tissue collection. Lung tissues were minced and digested with collagenase II and DNase I at 37 °C for 1 h, followed by filtration through 70 μm cell strainers. Red blood cells were removed using red blood cell lysis buffer where required. Spleens were mechanically dissociated through 70 μm strainers, followed by erythrocyte lysis. BALF cells were collected by centrifugation and washed with PBS before staining.

For surface staining, cells were first stained with a fixable viability dye to exclude dead cells and then stained with fluorophore-conjugated antibodies for 30 min at 4 °C in staining buffer. For intracellular cytokine analysis, pulmonary lymphocytes or splenocytes were restimulated with Kp-antigen or peptide pools for 12 h in the presence of protein transport inhibitor cocktail. Cells were then stained for surface markers, fixed, permeabilized and stained intracellularly for IFN-γ, IL-4 and IL-17. CD4⁺ T cells were analysed for IFN-γ, IL-4 and IL-17, whereas CD8⁺ T cells were analysed for IFN-γ and IL-17.

For dendritic-cell maturation analysis, BMDCs were gated as live CD11c⁺ cells and analysed for MHC-II, CD80 and CD86 expression. Mature BMDCs were quantified as CD80⁺MHC-II⁺ and CD86⁺MHC-II⁺ populations. For in vivo mCre reporter experiments, lung single-cell suspensions from Rosa26-LSL-tdTomato mice were analysed for td-Tomato expression within pulmonary dendritic cells and macrophages. Dendritic cells and macrophages were defined using CD45, CD11c, CD11b and F4/80 staining.

For humoral immune-cell analysis, T follicular helper cells were identified as CD45⁺CD3⁺CD4⁺CXCR5⁺PD-1⁺ cells, and germinal-centre B cells were identified as B220⁺GL7⁺Fas⁺ cells. Kp-antigen-specific memory B cells were detected using fluorescently labelled antigen probes together with B-cell and memory markers. For T-cell memory analysis, CD4⁺ and CD8⁺ T cells were gated from live CD45⁺CD3⁺ lymphocytes. Tissue-resident memory T cells were defined as CD69⁺CD103⁺ cells, central-memory T cells as CD44⁺CD62L⁺ cells and effector-memory T cells as CD44⁺CD62L⁻ cells within the CD4⁺ or CD8⁺ T-cell compartment.

For BALF myeloid-cell analysis after bacterial challenge, neutrophils were defined as CD45⁺CD11b⁺Ly6G⁺ cells and macrophages as CD45⁺CD11b⁺F4/80⁺ cells. For T-cell depletion experiments, depletion efficiency was confirmed in lung single-cell suspensions by quantifying CD4⁺ and CD8⁺ T cells among live CD45⁺CD3⁺ lymphocytes. For CD8β-depletion experiments, CD8⁺ T cells were detected using a staining antibody against CD8α to avoid interference from the depleting anti-CD8β antibody where applicable. Flow cytometry data were acquired using a BD FACSCanto II cytometer and analysed with FlowJo software. Doublets were excluded using forward-scatter height and area parameters, and dead cells were excluded before population analysis. Compensation was performed using single-stained controls. Representative gating strategies used for population definition are provided in **Supplementary Fig. 3-6**.

### Statistical analysis

Unless otherwise specified, all data for bar charts are presented as mean ±standard deviation (SD). GraphPad Prism 8.0 was used for statistical analysis. Two-group comparisons were performed using two-tailed unpaired Student’s *t*-tests when appropriate. Multiple-group comparisons were analysed by one-way or two-way ANOVA followed by appropriate post hoc tests. Survival curves were compared using the log-rank test. Bacterial burden data were log-transformed when appropriate before statistical testing. *p* < 0.05 was considered statistically significant.

## Acknowledgements

This work was supported by the National Natural Science Foundation of China (NSFC, Grant No. 32370993), the major project of Study on Pathogenesis and Epidemic Prevention Technology System (Grant No. 2025ZD01903302, 2021YFC2302500 and 2024YFC2310804) by the Ministry of Science and Technology of China, and the Natural Science Foundation of Chongqing (CSTB2025NSCQ-GPX0624). We thank Yajuan Zhang (Clinical Medical Research Center, Southwest Hospital, Third Military Medical University, China) for technical support with flow cytometry.

## Author contributions

S.G. conceived and directed the project and supervised the research and contributed experimental materials. Y.L., J.Z., Z.C., R.L., Q.X., C.L. performed experiments. Y.L. and C.L. completed the design of the machine learning model and finalized the curation datasets. Y.L., J.Z. analyzed data. S.G., J.Z., Y.L. wrote the manuscript with comments from all authors.

## Competing interests

The authors declare no competing interests.

## Data availability

All data supporting the results in this study are available within the paper and its Supplementary Information. Source data are provided with this paper.

## Code availability

The source code and model files supporting the findings of this study are publicly available on GitHub at: https://github.com/LYH-Exo/ML-Pep.

## Extended Data Figures

**Extended Data Fig. 1.**
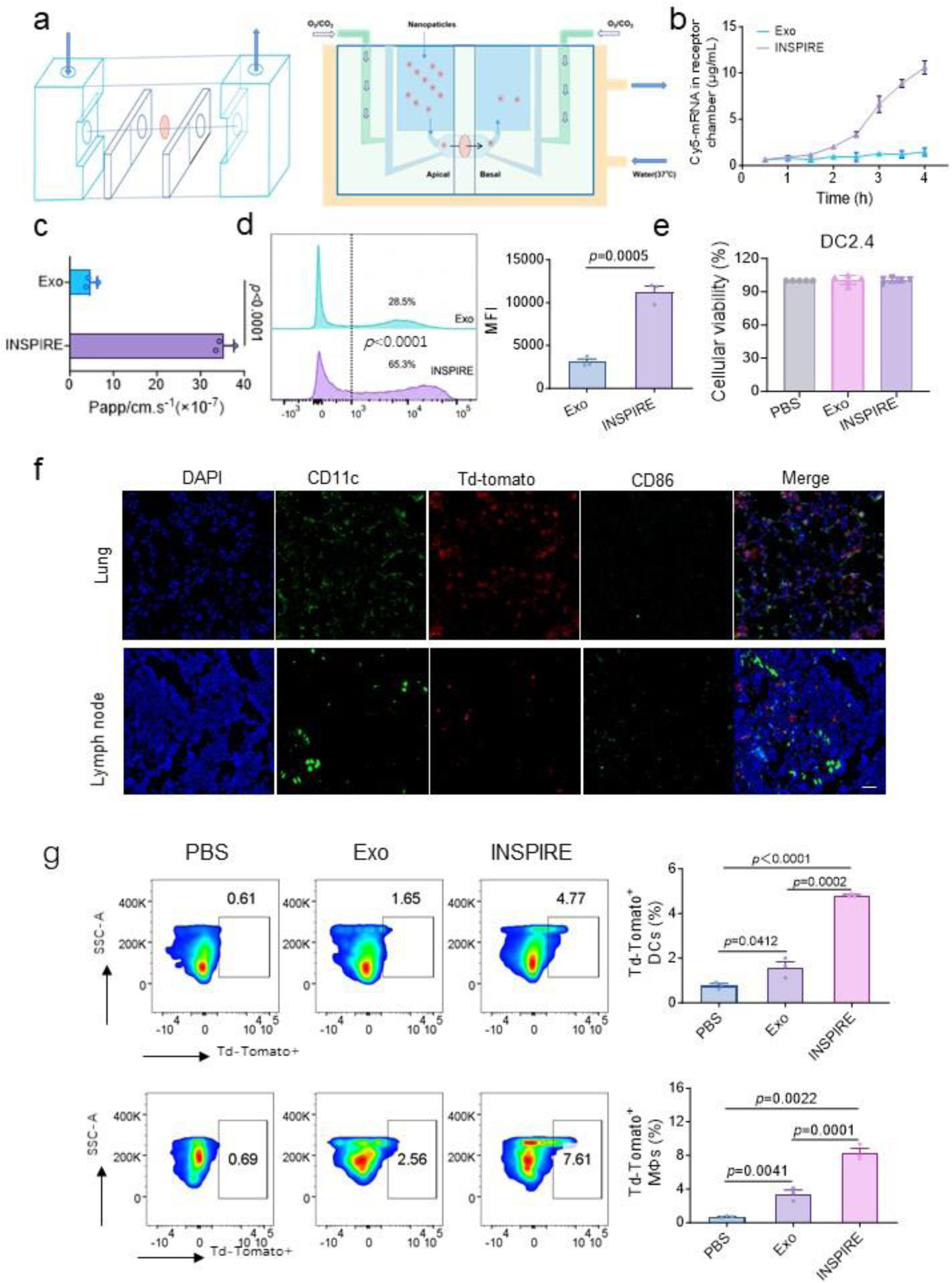
Transepithelial transport, cellular uptake and in vivo reporter activation mediated by INSPIRE. **a**, Schematic illustration of the Ussing chamber system used to evaluate transepithelial transport of Cy5-labelled mRNA delivered by Exo or INSPIRE across an epithelial barrier. **b**, Time-dependent accumulation of Cy5-labelled mRNA in the receptor chamber after treatment with Exo or INSPIRE. **c**, Apparent permeability coefficient of Cy5-labelled mRNA delivered by Exo or INSPIRE in the Ussing chamber assay. **d**, Flow-cytometric analysis and quantification of Cy5-labelled mRNA uptake in DC2.4 cells after treatment with Exo or INSPIRE. **e**, Cell viability of DC2.4 cells after treatment with PBS, Exo or INSPIRE, measured by Cell Counting Kit-8 assay. f, Representative immunofluorescence images of lung and mediastinal lymph node sections from tdTomato reporter mice after intranasal delivery of Cre recombinase mRNA formulated with Exo. Nuclei were stained with DAPI (blue), CD11c is shown in green, tdTomato is shown in red and CD86 is shown in cyan. **g**, Flow-cytometric analysis and quantification of tdTomato-positive dendritic cells and macrophages in the lung after intranasal delivery of Cre recombinase mRNA formulated with PBS, Exo or INSPIRE.

**Extended Data Fig. 2.**
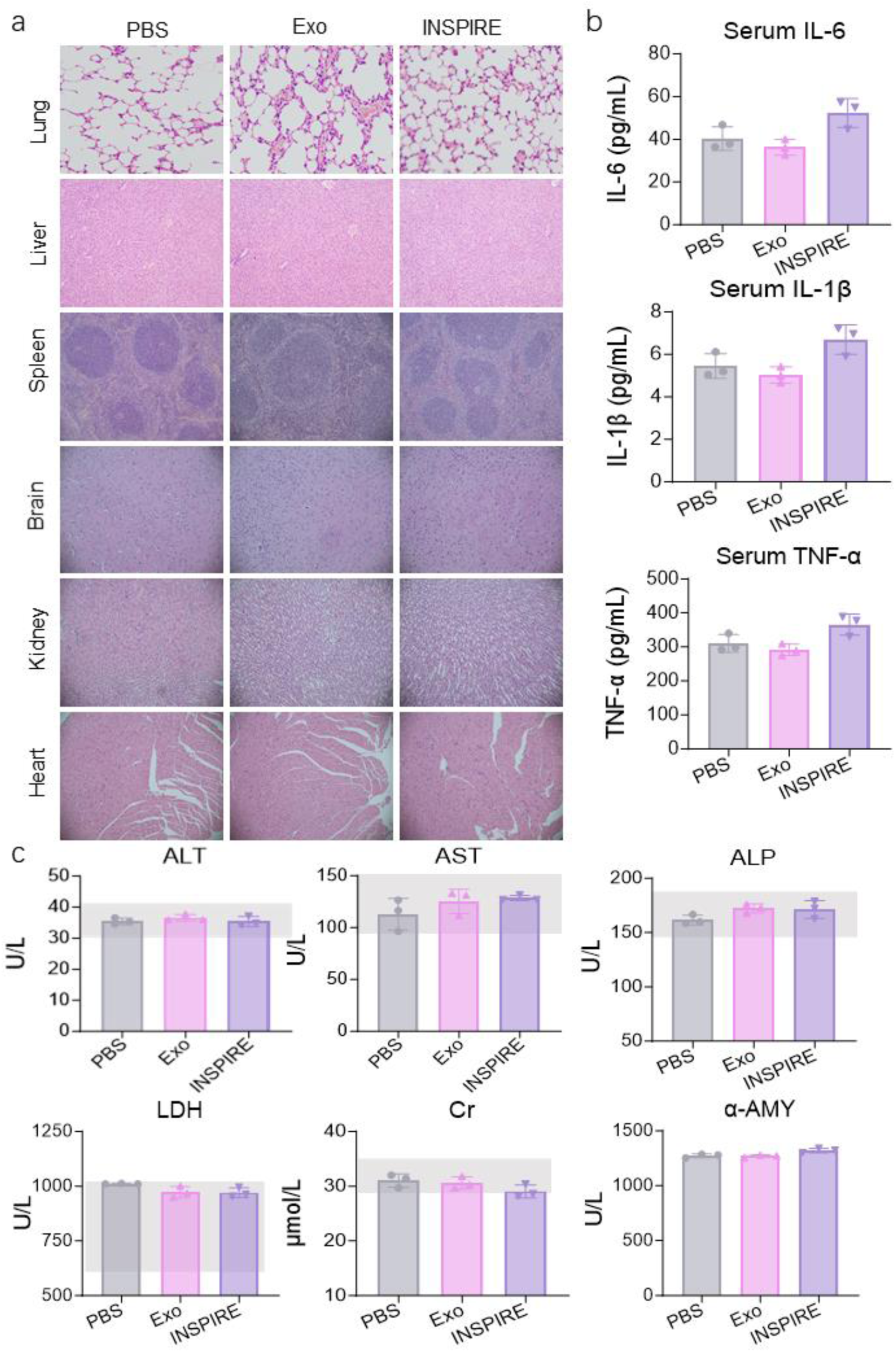
Additional in vivo safety evaluation of INSPIRE after intranasal administration. **a**, Representative haematoxylin and eosin (H&E) staining images of the lung, liver, spleen, brain, kidney and heart collected from mice treated with PBS, Exo or INSPIRE. **b**, Serum levels of interleukin-6 (IL-6), IL-1β and tumour necrosis factor-α (TNF-α) in mice after treatment with PBS, Exo or INSPIRE. **c**, Serum biochemical indices, including alanine aminotransferase (ALT), aspartate aminotransferase (AST), alkaline phosphatase (ALP), lactate dehydrogenase (LDH), creatinine (Cr) and α-amylase (α-AMY), in mice after treatment with PBS, Exo or INSPIRE.

**Extended Data Fig. 3.**
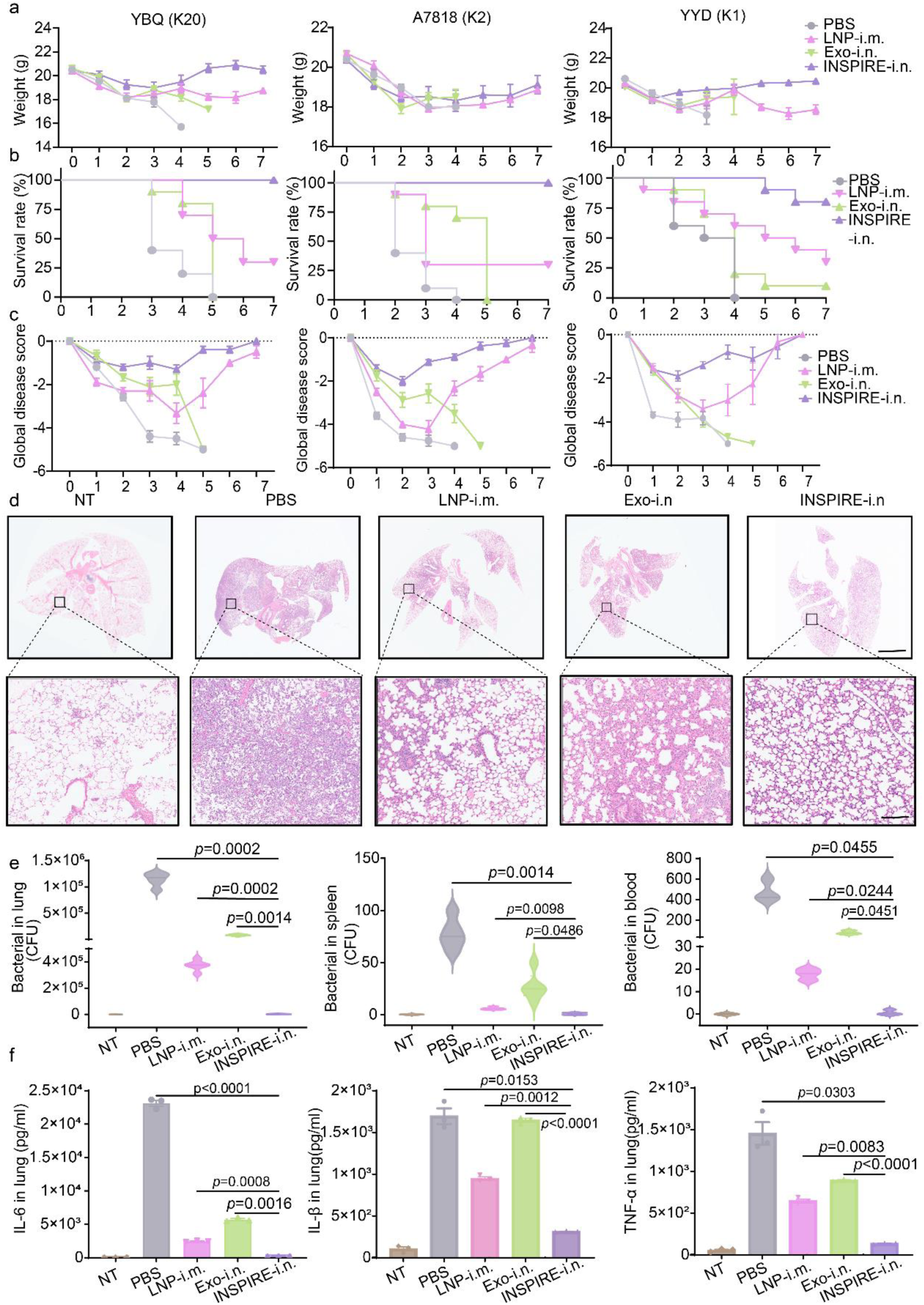
INSPIRE-based mRNA vaccination confers cross-strain protection against pulmonary Kp infection. **a-c**, Body weight changes (**a**), survival rates (**b**), and global disease scores (**c**) of vaccinated mice following intratracheal challenge with three Kp strains (YBQ, A7818, and YYD) over 7 days post-challenge (dpc) (n = 10 biologically independent animals). **d**, Representative H&E-stained lung sections from mice collected 2 dpc after intratracheal challenge with the YBQ strain. NT: untreated and unchallenged control. Scale bars: 2 mm (upper overview), 100 μm (lower magnified insets). **e**, Bacterial burden in the lung, spleen, and blood at 2 dpc after intratracheal challenge with the YBQ strain (n = 5 biologically independent samples). **f,** Pro-inflammatory cytokine levels, IL-6, IL-1β, and TNF-α, in lung homogenates at 3 dpc after intratracheal challenge with the YBQ strain (n = 4 biologically independent samples). Violin plots with medians and quartiles for (**e**).

**Extended Data Fig. 4.**
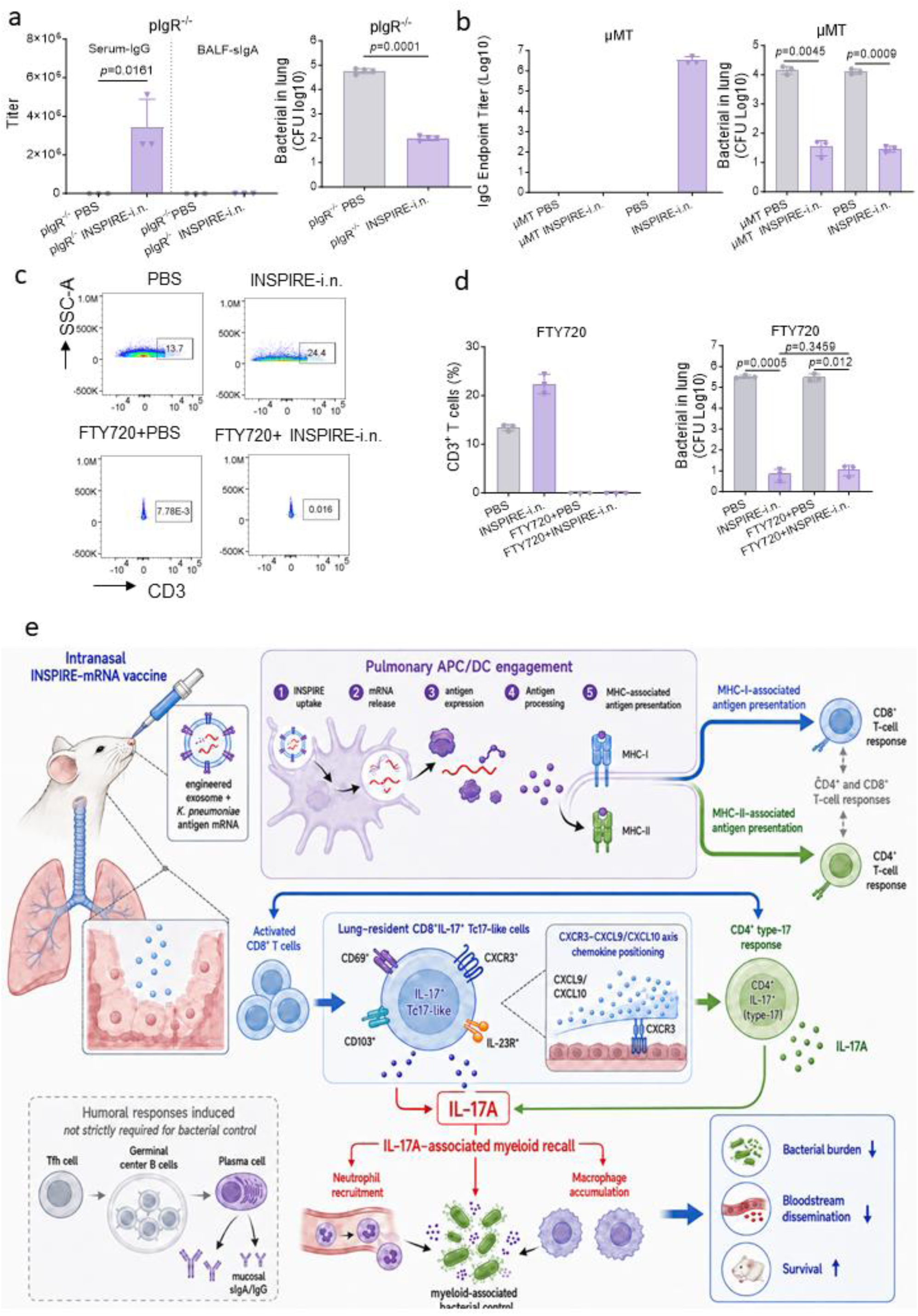
Humoral-immunity and lymphocyte-egress perturbation analyses of INSPIRE-mediated protection. **a**, Kp-Antigen-specific serum IgG and bronchoalveolar lavage fluid (BALF) secretory IgA (sIgA) endpoint titres in polymeric immunoglobulin receptor-deficient (pIgR^−/−^) mice intranasally immunized with PBS or INSPIRE. Pulmonary bacterial burdens were quantified 48 h after intratracheal challenge with *Klebsiella pneumoniae* YBQ. Despite the near absence of BALF sIgA, INSPIRE-i.n. significantly reduced pulmonary bacterial burdens in pIgR^−/−^ mice. **b**, Serum IgG levels, expressed as absorbance at 450 nm, in B-cell-deficient μMT mice intranasally immunized with PBS or INSPIRE. Pulmonary bacterial burdens were measured 48 h after YBQ challenge in μMT mice and background-matched wild-type mice. INSPIRE-i.n. significantly reduced lung bacterial burdens in both genotypes, indicating that mature B-cell-dependent antibody responses were not required for INSPIRE-mediated bacterial control. **c**, Representative flow-cytometry plots showing circulating CD3⁺ T cells in mice treated with PBS or INSPIRE-i.n., with or without fingolimod (FTY720)-mediated lymphocyte-egress blockade. **d**, Quantification of circulating CD3⁺ T-cell frequencies and pulmonary bacterial burdens in the treatment groups shown in **c**. FTY720 markedly reduced circulating CD3⁺ T-cell frequencies but did not significantly impair INSPIRE-mediated control of pulmonary bacterial burden after YBQ challenge. **e**, Proposed model summarizing INSPIRE-mediated mucosal protection against pulmonary Kp.

## Supplementary information

### Supplementary Table

**Supplementary Table1.**
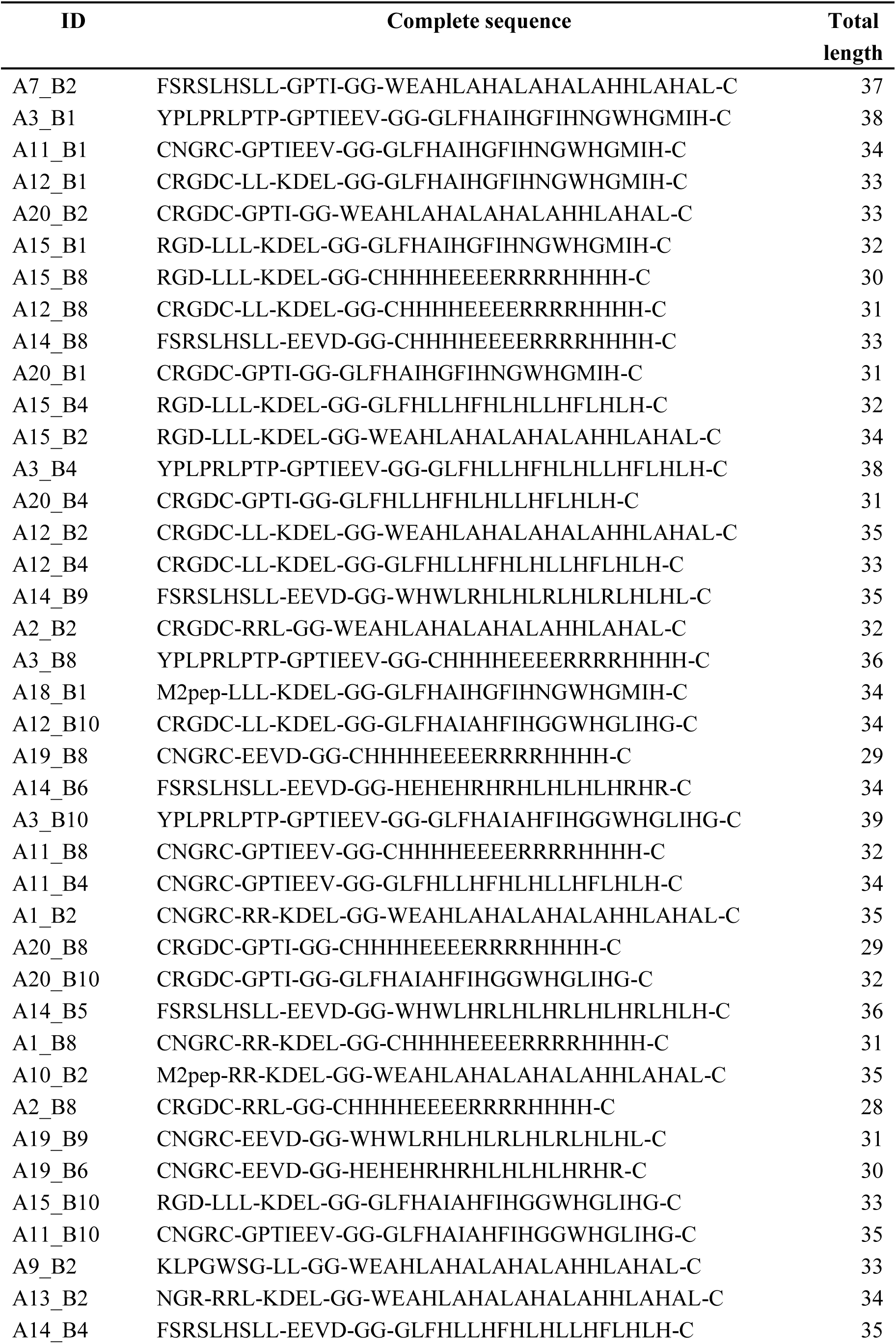
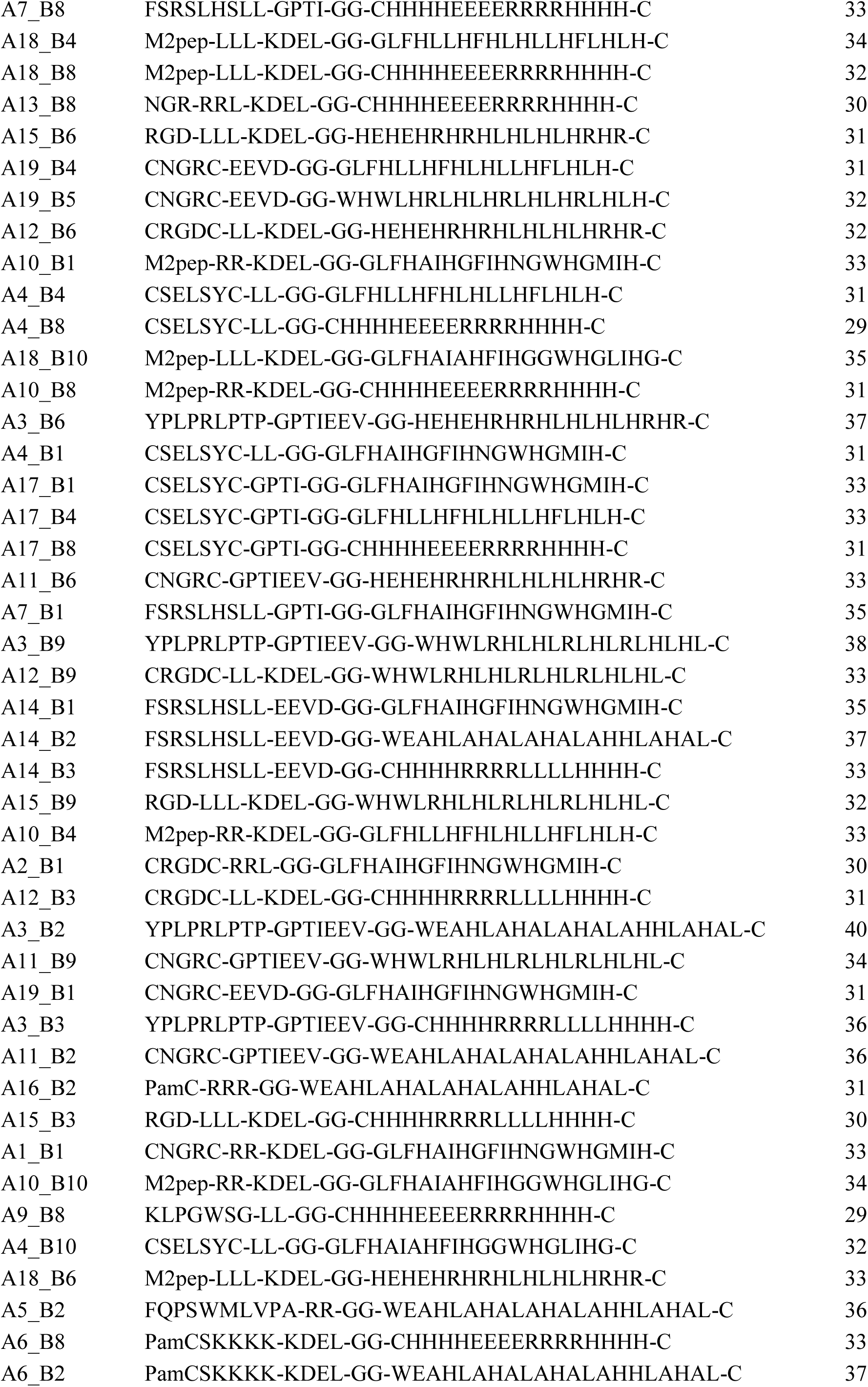
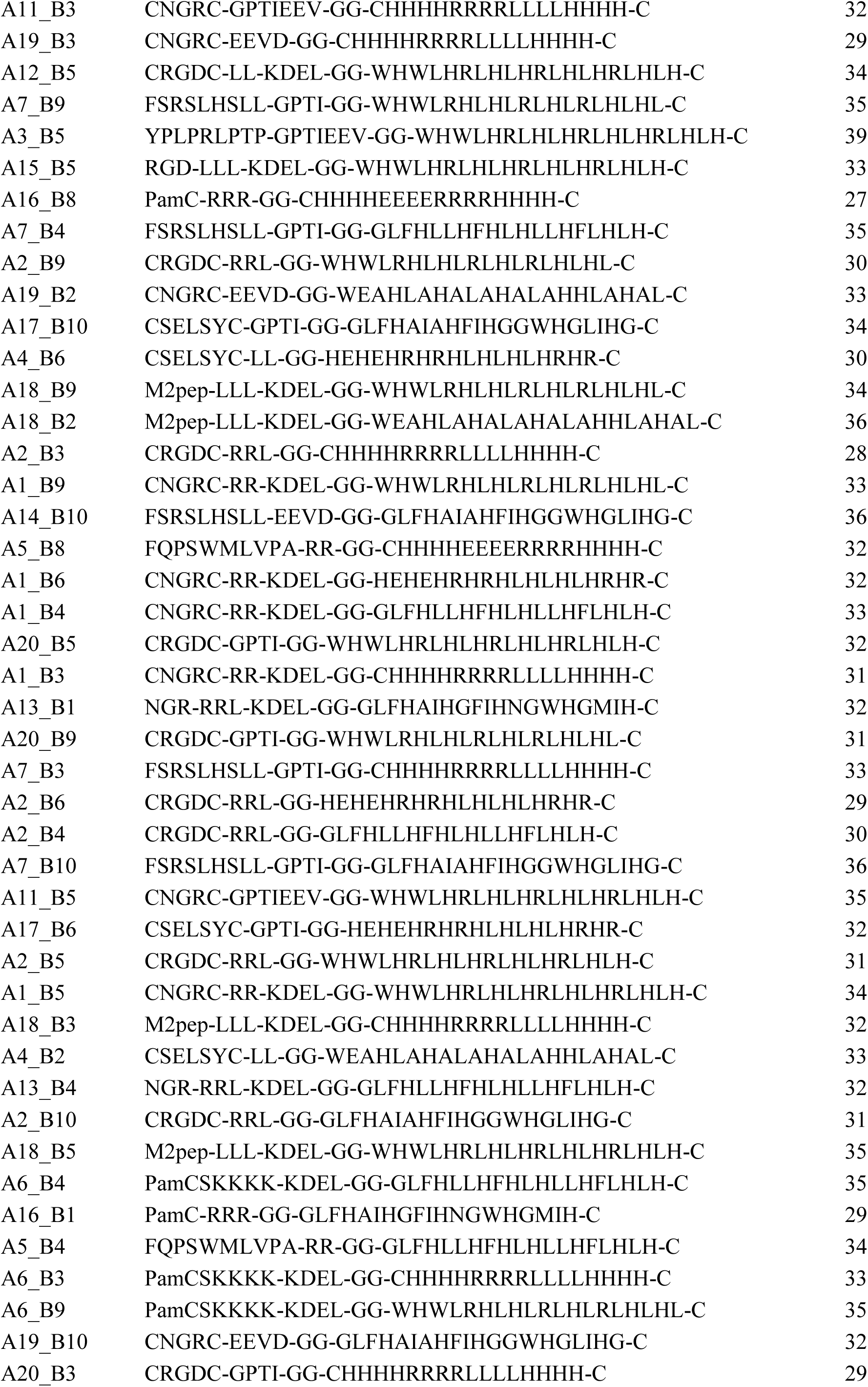
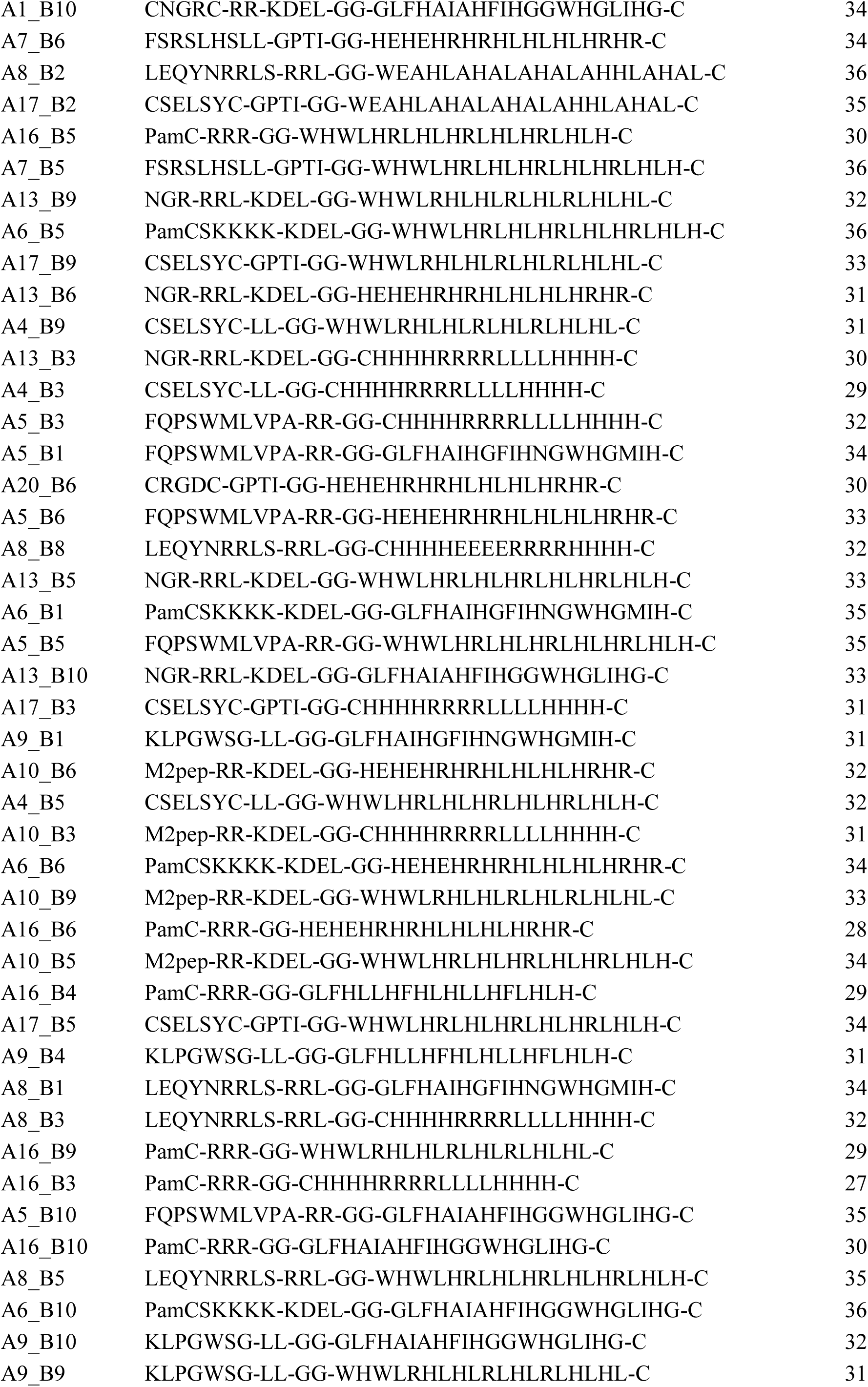
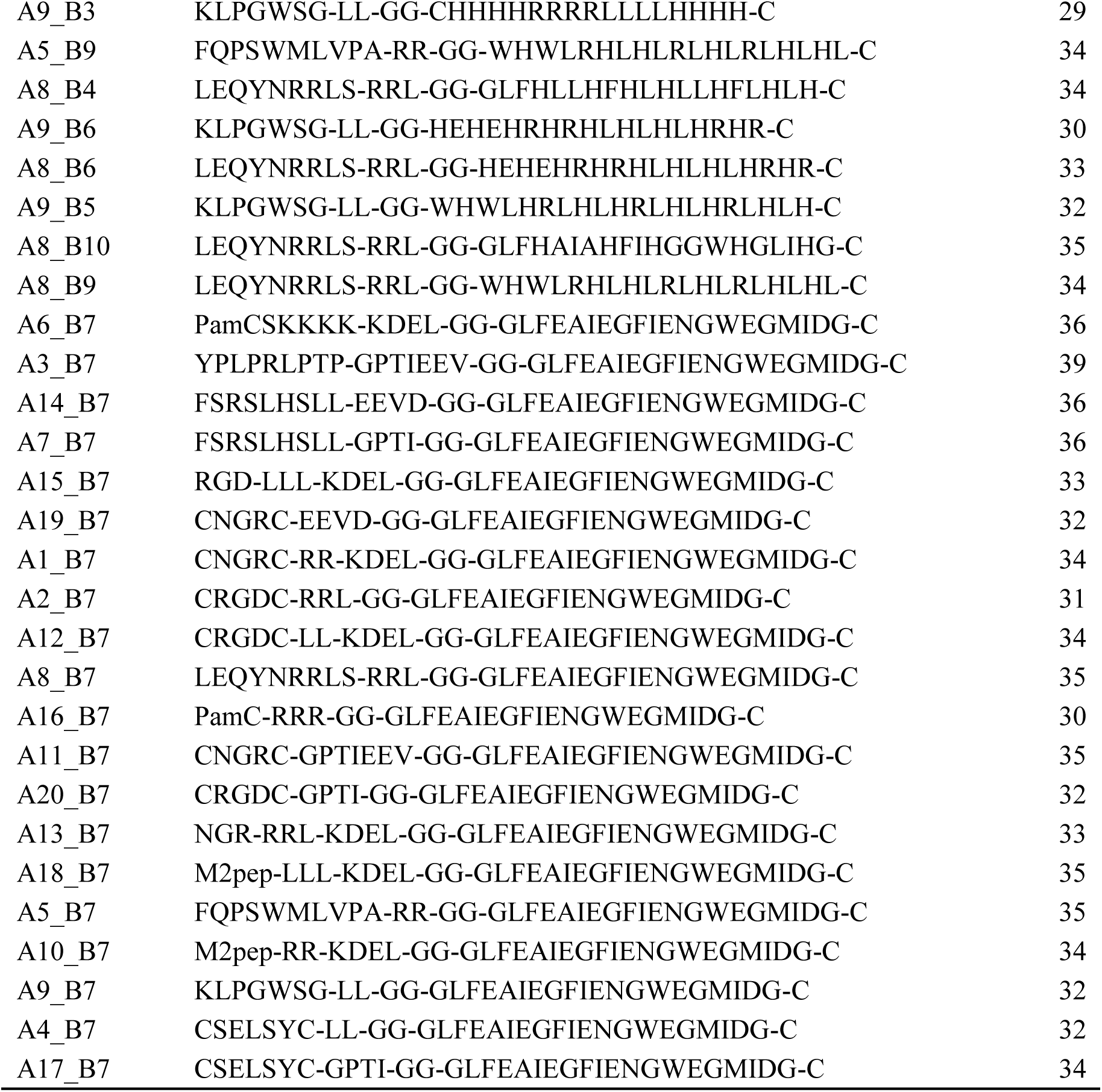
Peptides sequence.

**Supplementary Table 2.**
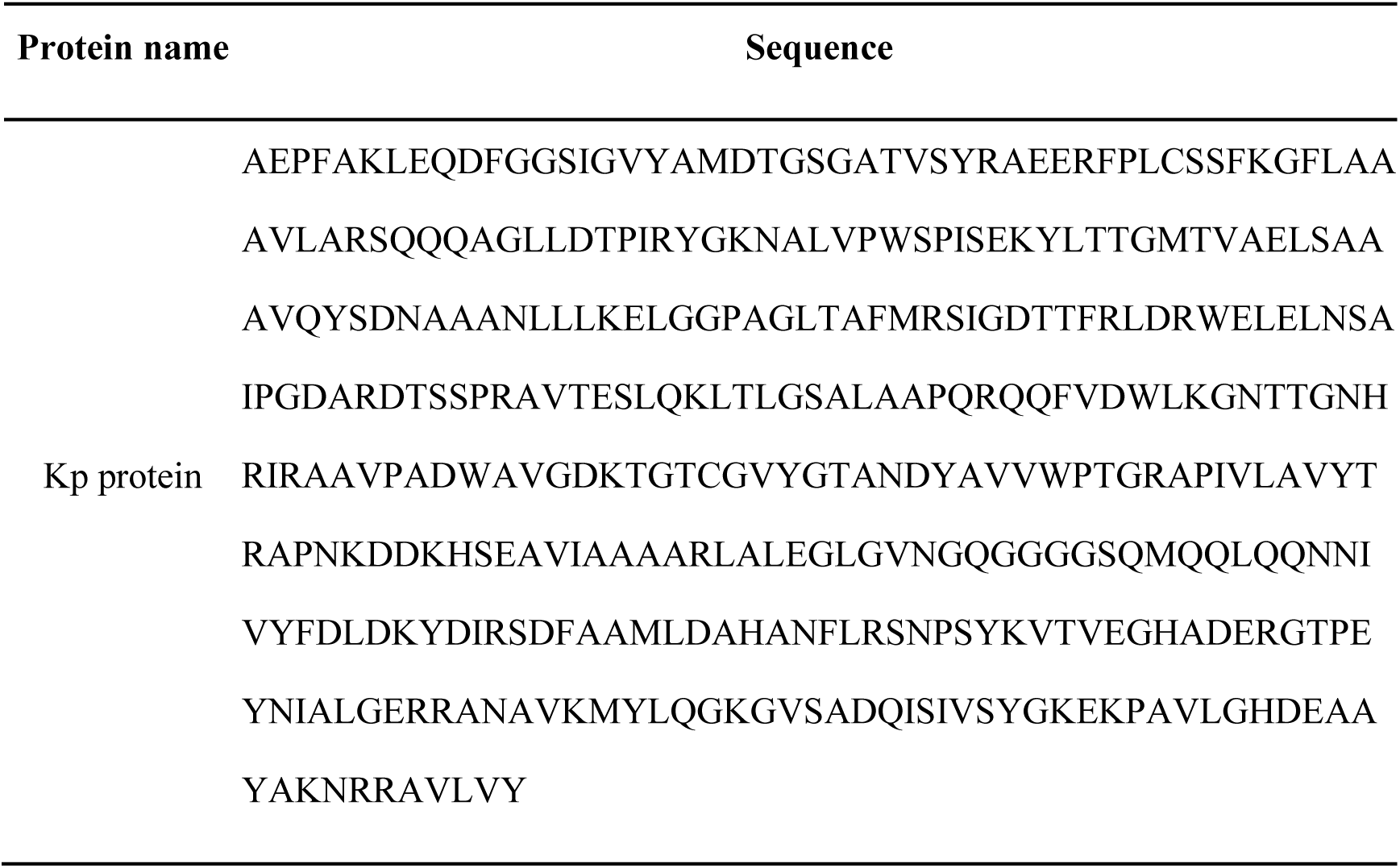
Kp protein sequence.

**Supplementary Table 3.**
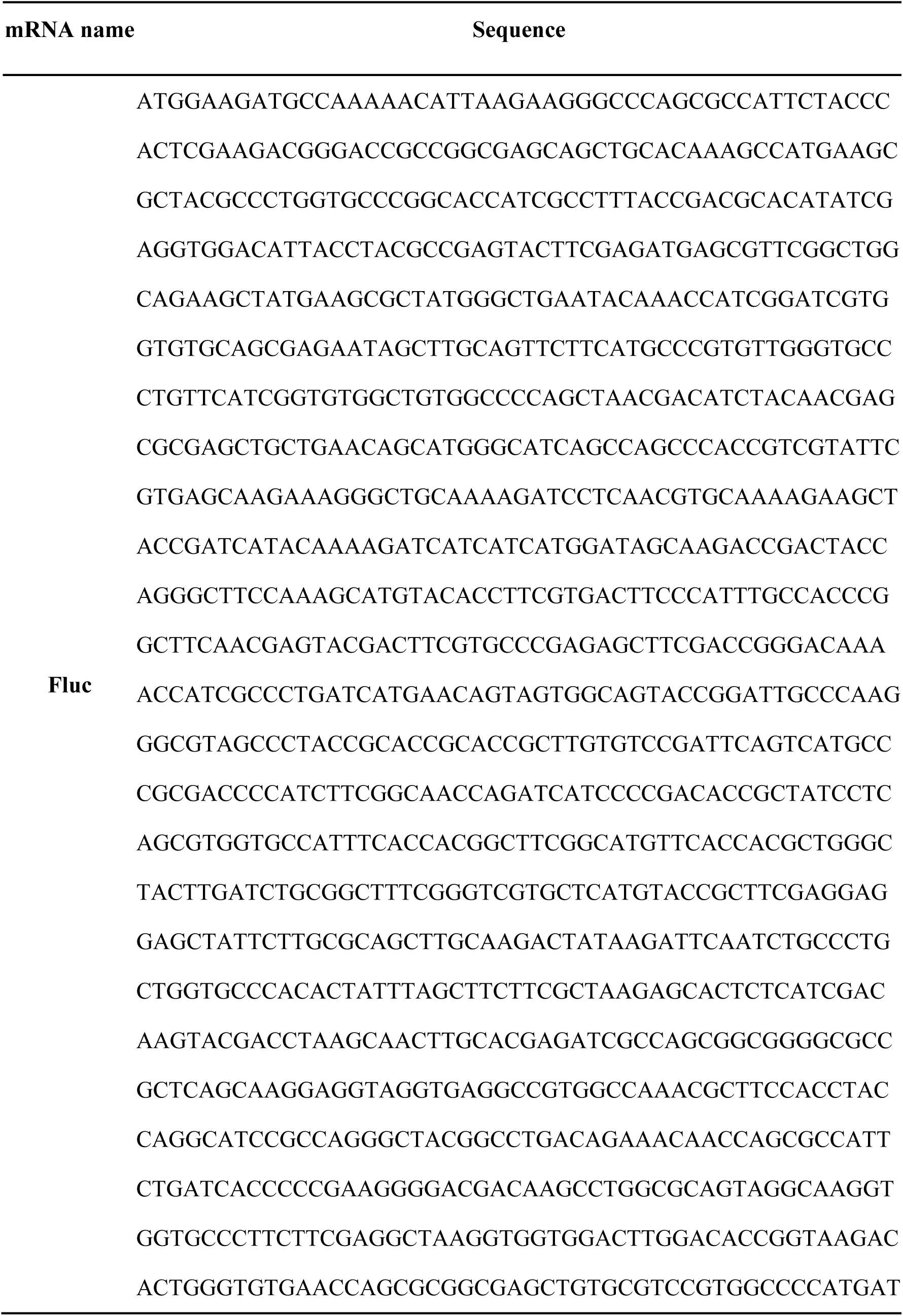
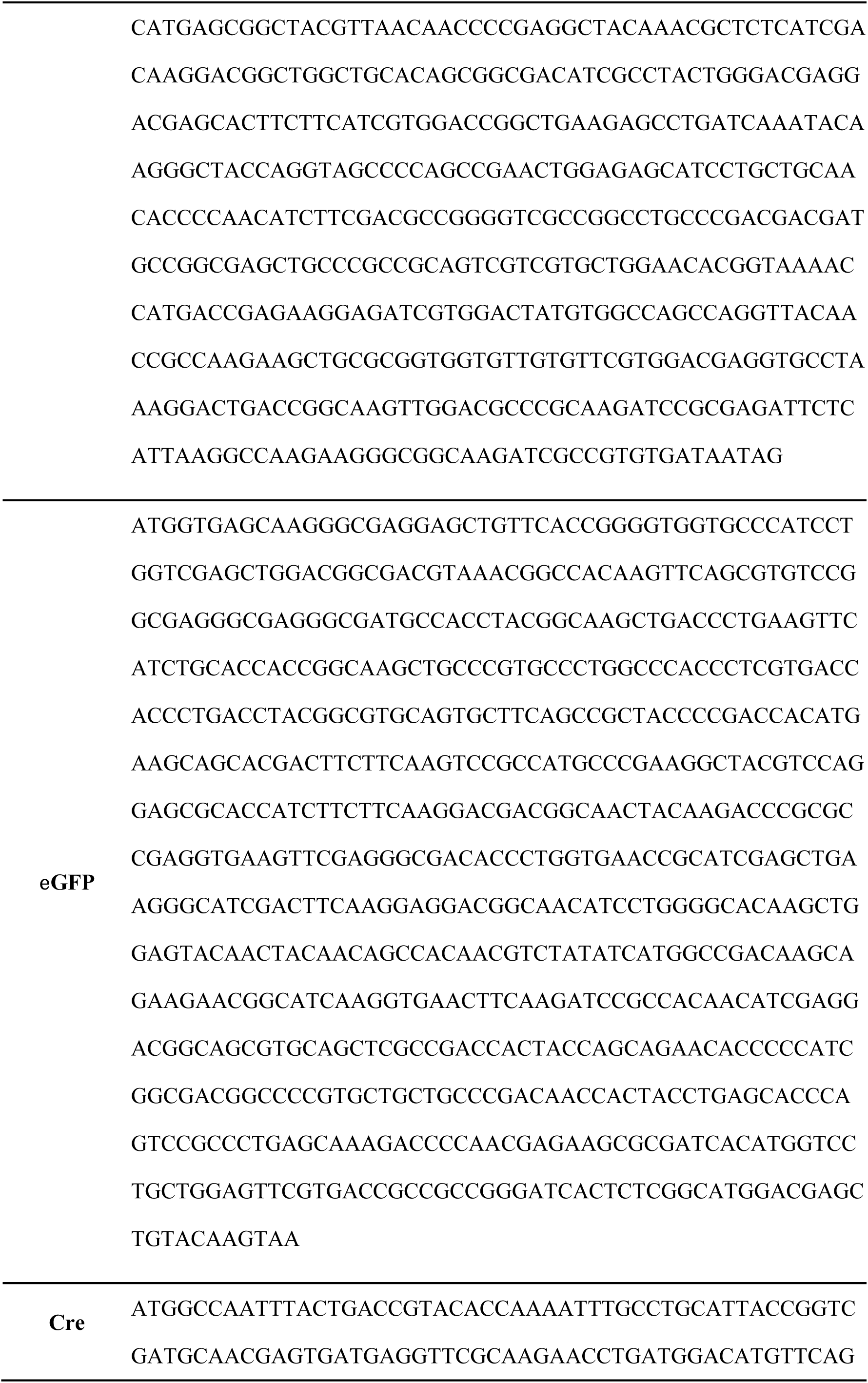
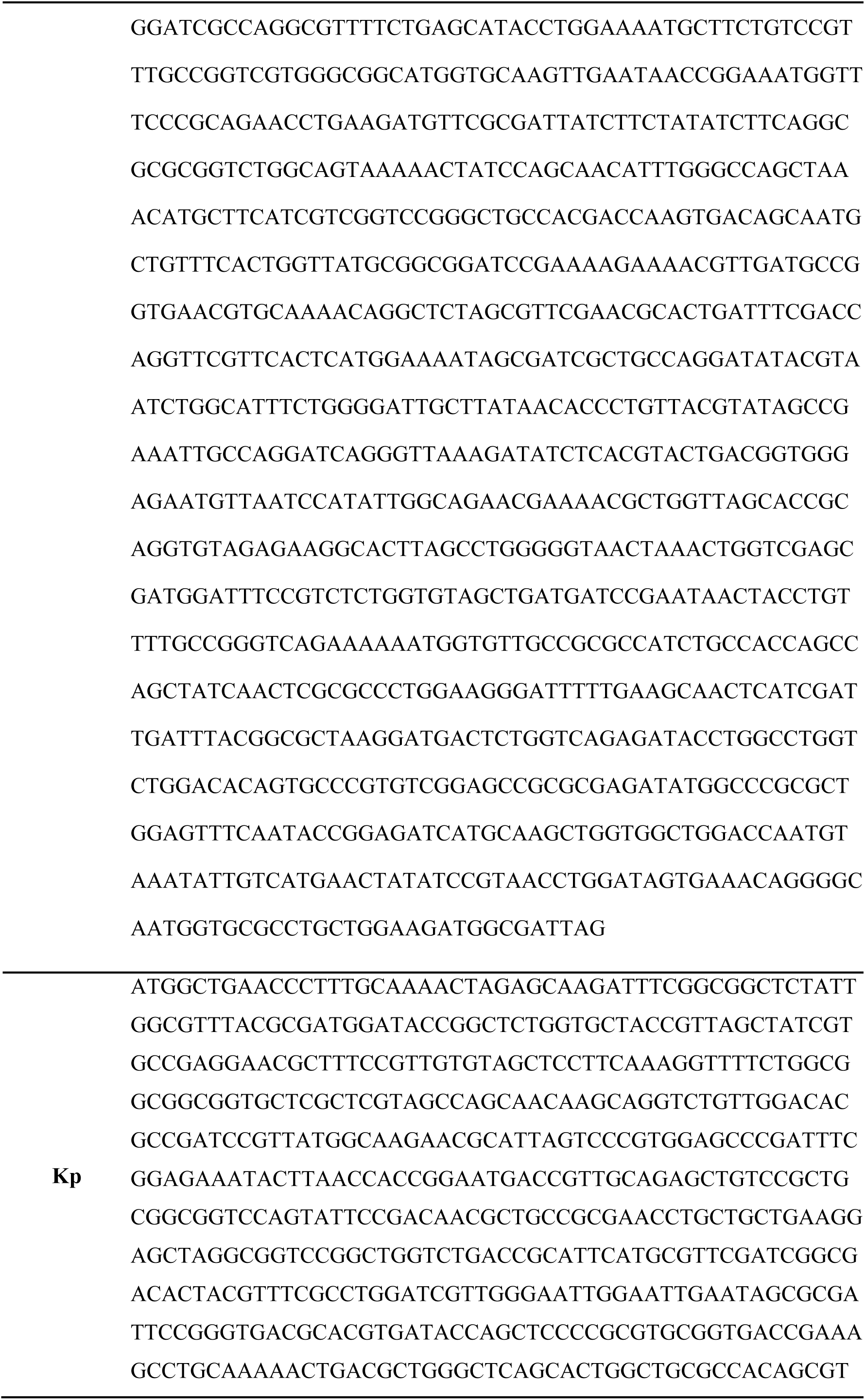
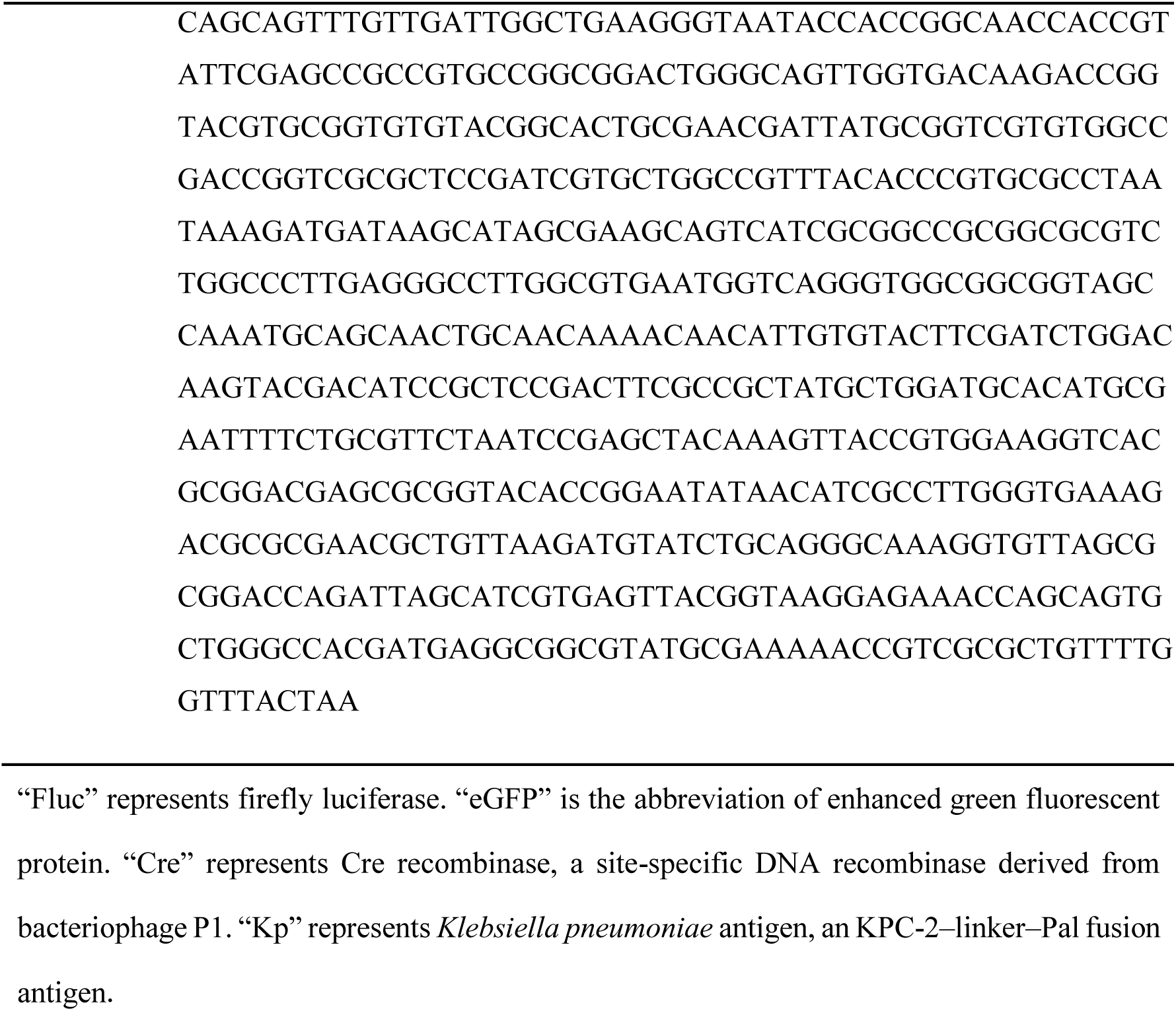
The synthetic design sequence (SDS) sequences of *in vitro* transcribed mRNA adopted in this study.

**Supplementary Table 4.**
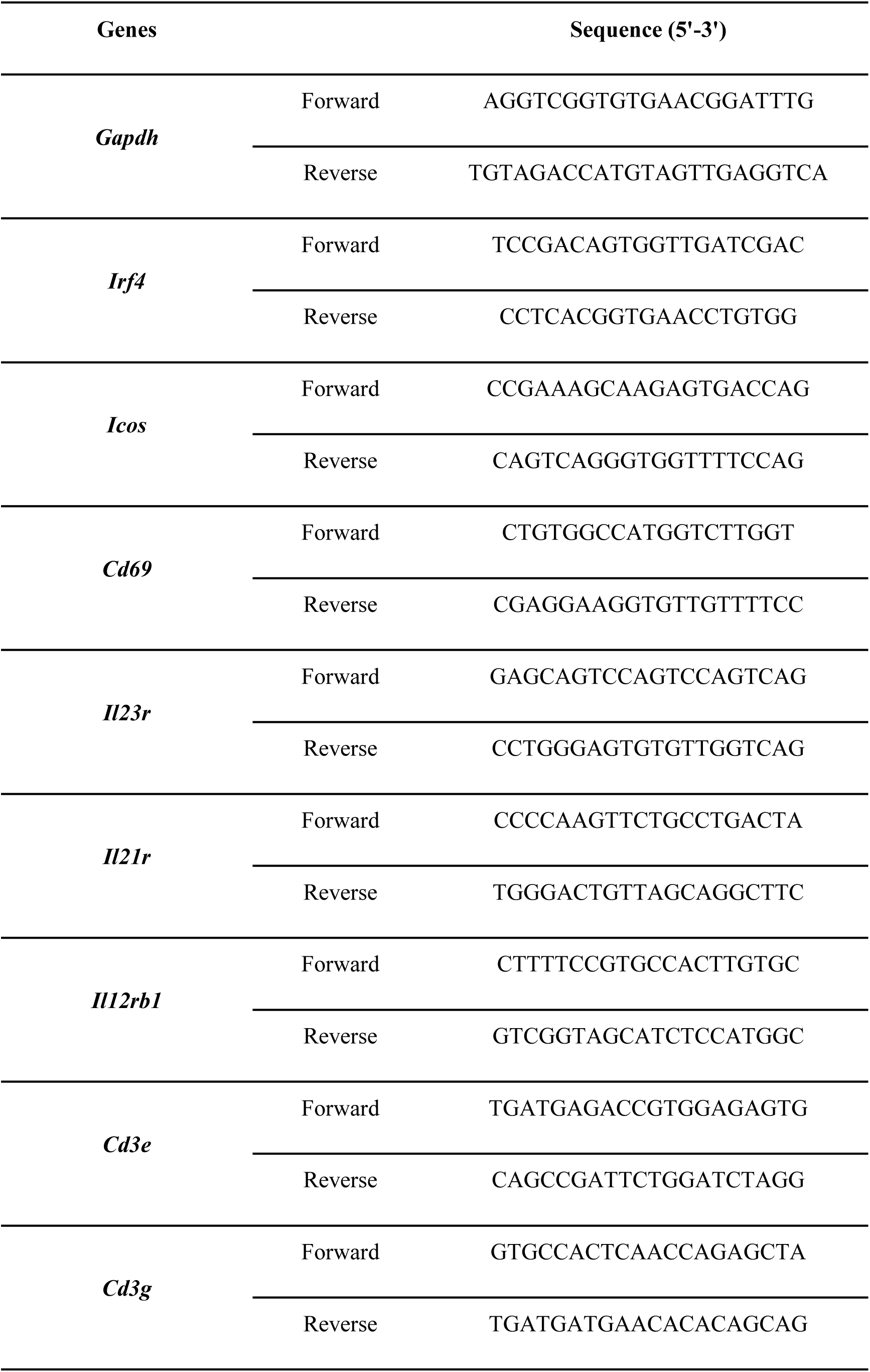
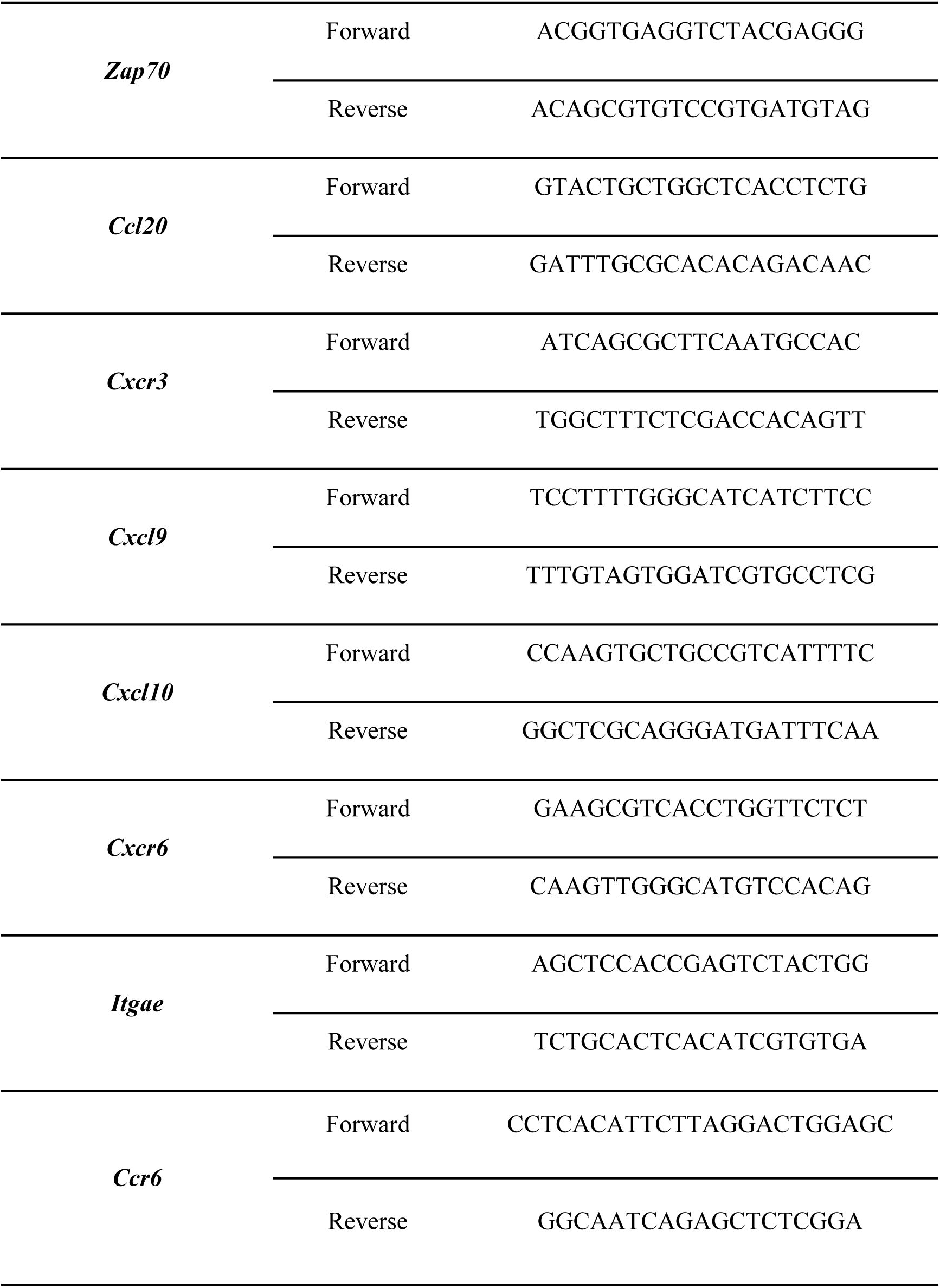
Primer sequences.

**Supplementary Table 5.** Antibodies and depletion reagents used in this study.

| Fixable viability dye | AF700 |
| --- | --- |
| CD80 | Percp-Cy5.5 |
| CD86 | APC |
| MHC-II | BV421 |
| CD45 | APC-Cy7 |
| CD11c | FITC |
| CD11b | PE |
| F4/80 | Percp-Cy5.5 |
| Ly6G | eF450 |
| CD3 | FITC |
| CD4 | Percp-Cy5.5 |
| CD8α | PE-Cy7 |
| CXCR5 | APC |
| PD-1 | PE |
| B220 | eF450 |
| Gr-1 | PE-Cy7 |
| Fas | PE |
| CD69 | BV421 |
| CD103 | BV510 |
| CD44 | APC |
| CD62L | PE |
| IFN-γ | eF450 |
| IL-4 | APC |
| IL-17 | PE |
| Anti-CD4 monoclonal antibody | In vivo CD4<sup>+</sup> T-cell depletion |
| Anti-CD8β monoclonal antibody | In vivo CD8<sup>+</sup> T-cell depletion |
| FTY720 / Fingolimod | Lymphocyte-egress blockade experiment |

**Supplementary Table 6.**
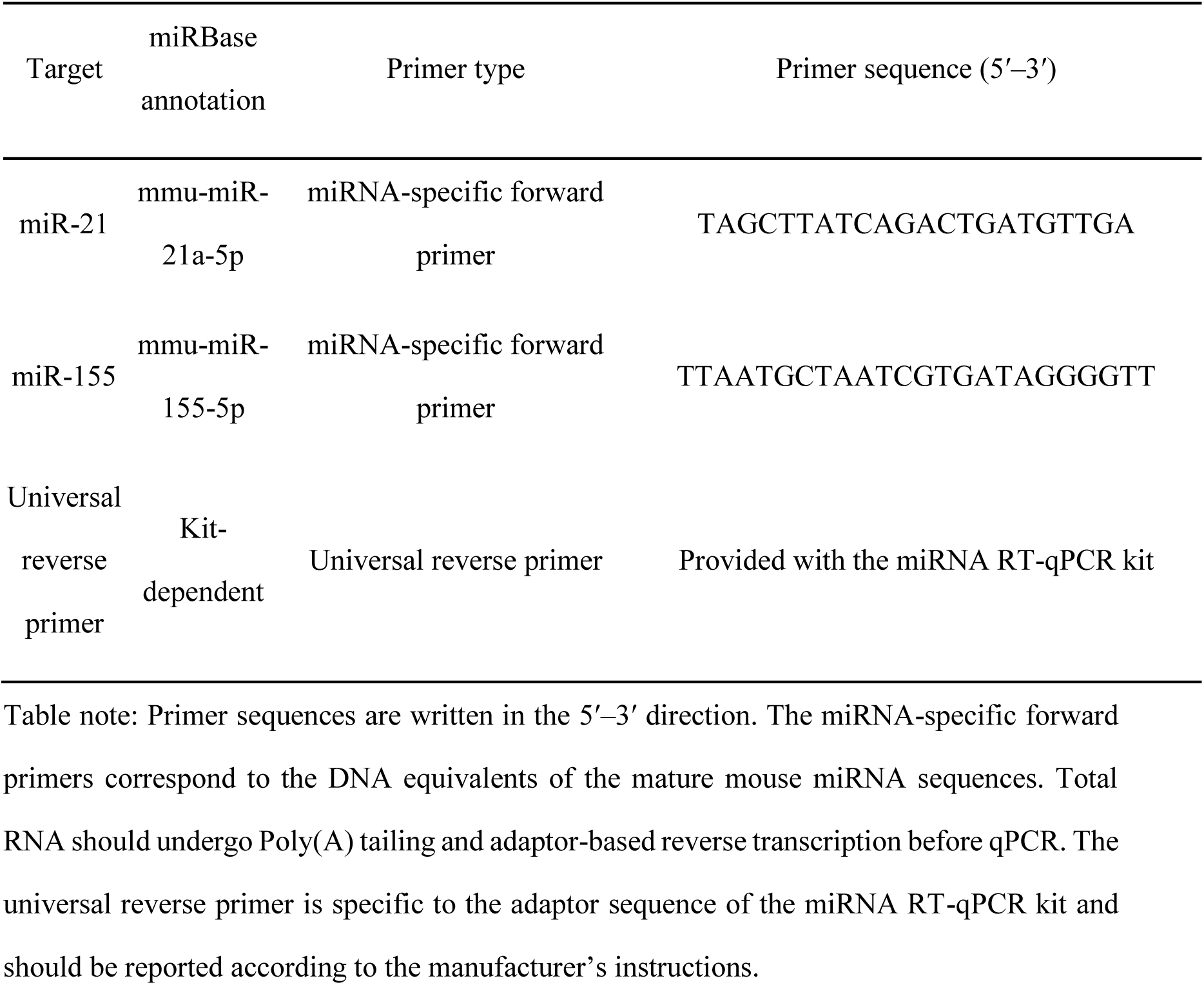
Primer sequences used for mouse mature miRNA RT-qPCR.

### SUPPLEMENTARY FIGURES

**Supplementary Fig. 1.**
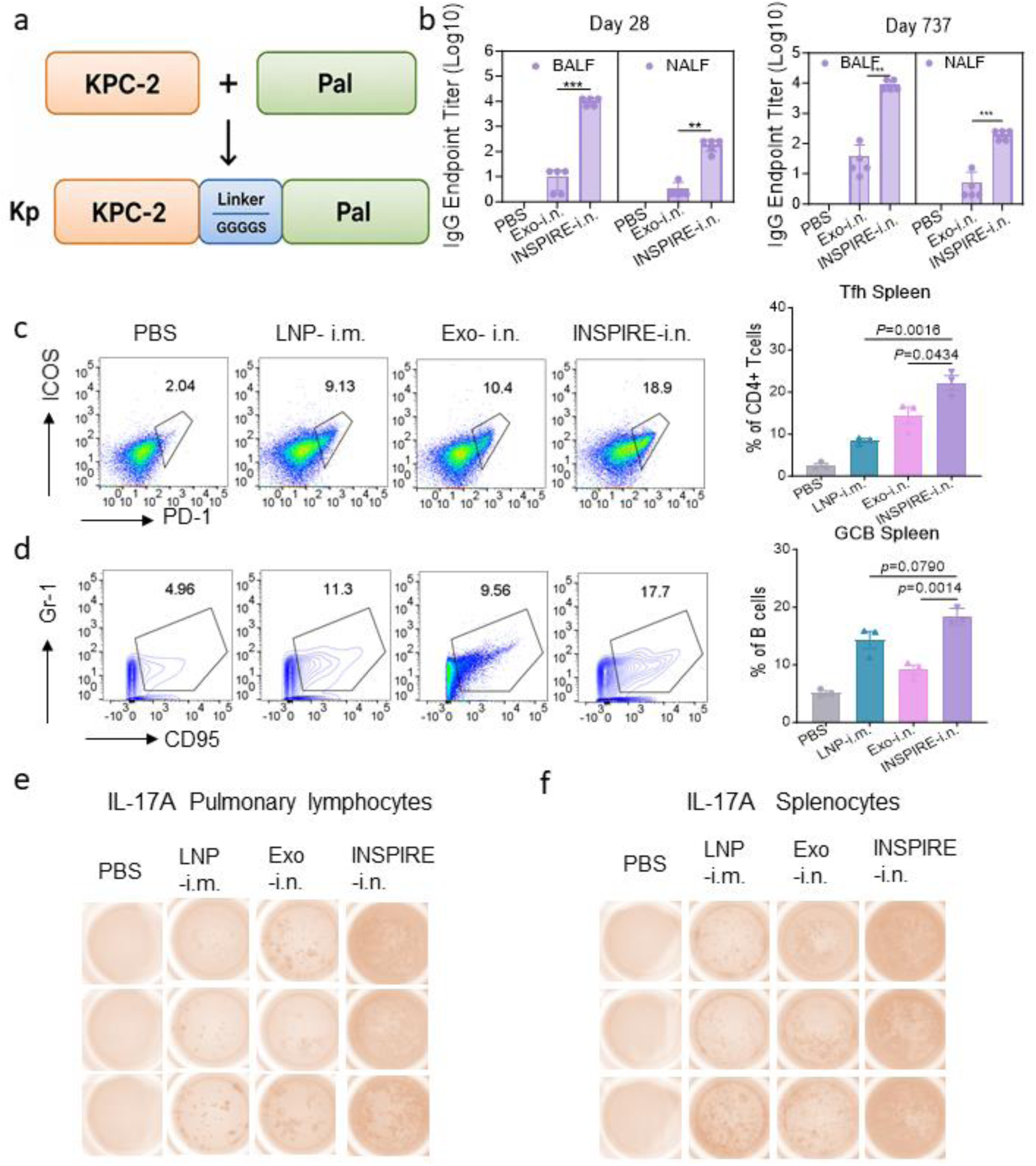
Antigen design and supporting immune analyses for INSPIRE-mediated mucosal mRNA vaccination. **a**, Schematic illustration of the Kp fusion antigen design. KPC-2 and Pal were linked by a flexible GGGGS linker to generate the KPC-2–linker–Pal fusion antigen. **b**, Endpoint titres of Kp-antigen-specific immunoglobulin G (Ig) in bronchoalveolar lavage fluid (BALF) and nasal lavage fluid (NALF) on day 28 and day 737 after immunization and recall challenge. **c**, Representative flow-cytometry plots and quantification of splenic T follicular helper (Tfh) cells from mice treated with PBS, LNP-i.m., Exo-i.n. or INSPIRE-i.n. **d**, Representative flow-cytometry plots and quantification of splenic germinal-centre B (GCB) cells from mice treated as in c. **e**, Representative enzyme-linked immunospot (ELISpot) images of interleukin-17A (IL-17A)-secreting cells in pulmonary lymphocytes after antigen re-stimulation. **f**, Representative ELISpot images of IL-17A-secreting cells in splenocytes after antigen re-stimulation.

**Supplementary Fig. 2.**
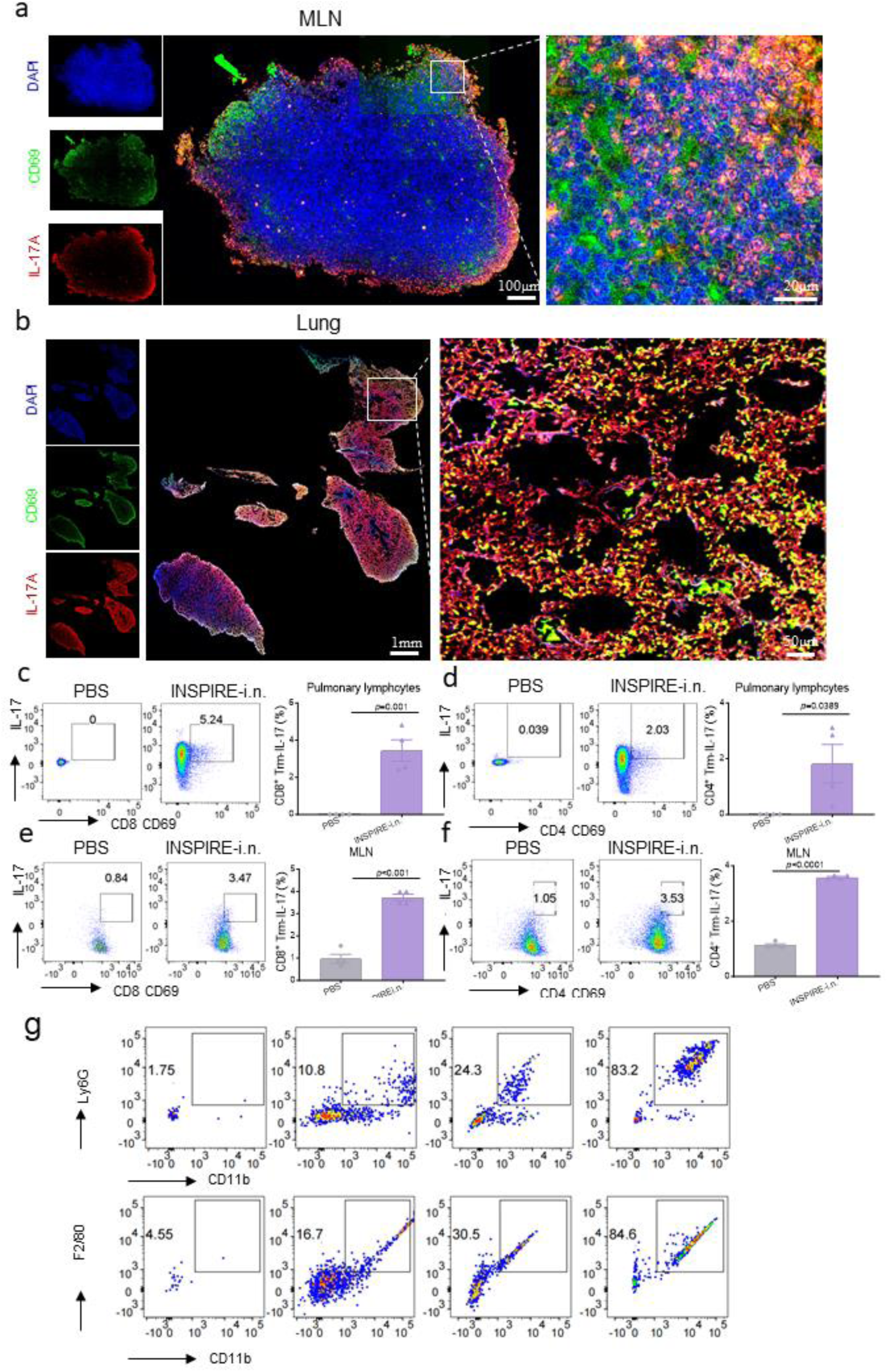
INSPIRE induces CD69-associated type-17 T-cell responses in the lung and mediastinal lymph nodes. **a**,**b**, Representative immunofluorescence images showing CD69 and IL-17A signals in mediastinal lymph nodes (MLN; **a**) and lung sections (**b**) after intranasal INSPIRE vaccination. Nuclei were stained with DAPI (blue), CD69 is shown in green and IL-17A immunofluorescence signal is shown in red. Boxed regions indicate areas shown at higher magnification. Scale bars, 100 μm and 20 μm in a; 1 mm and 50 μm in **b**. **c, d**, Representative flow-cytometry plots and quantification of CD69⁺IL-17⁺ cells within CD8⁺ T cells (**c**) and CD4⁺ T cells (**d**) in pulmonary lymphocytes from PBS- or INSPIRE-i.n.-immunized mice. **e, f,** Representative flow-cytometry plots and quantification of CD69⁺IL-17⁺ cells within CD8⁺ T cells (**e**) and CD4⁺ T cells (**f**) in mediastinal lymph nodes from PBS- or INSPIRE-i.n.-immunized mice.

**Supplementary Fig. 3.**
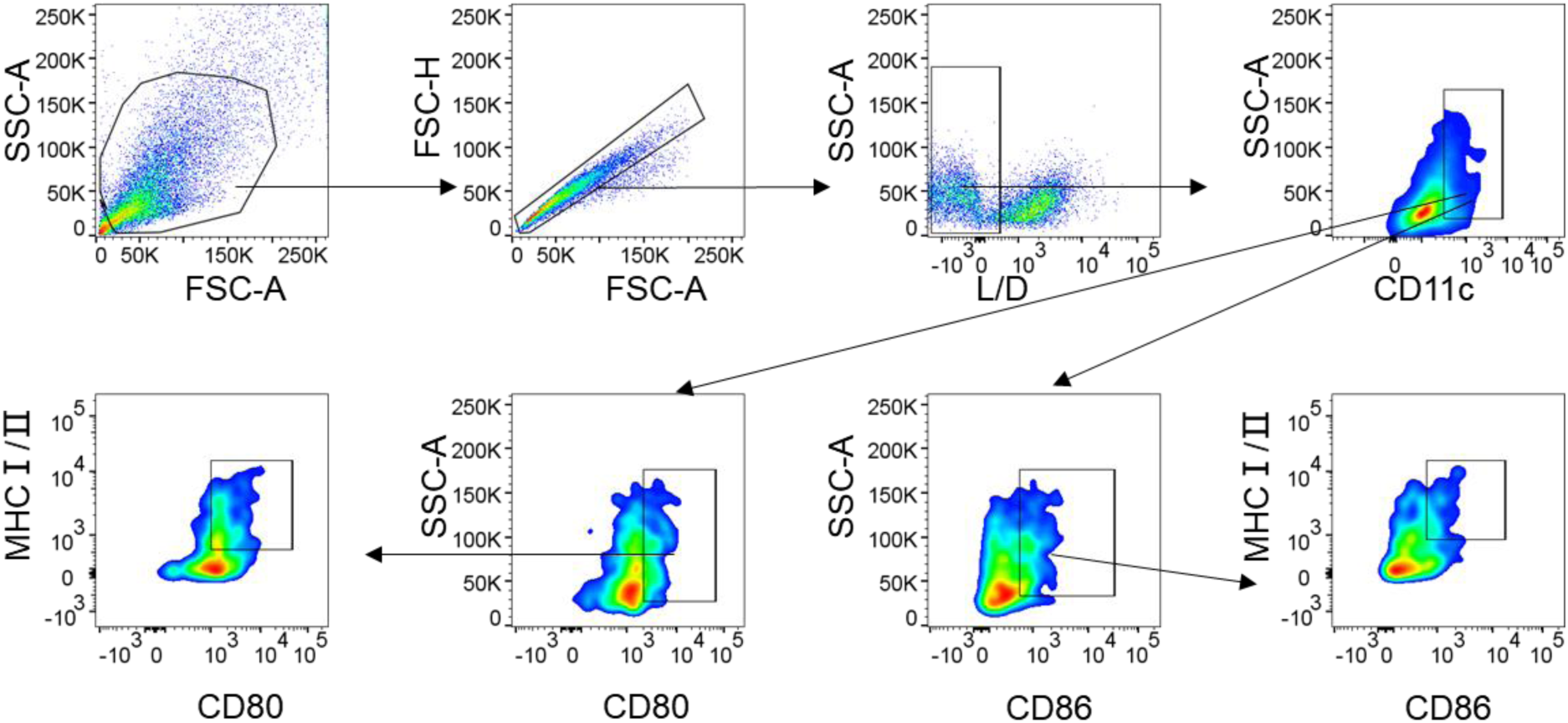
Representative flow-cytometry gating strategy for BMDC activation and antigen-presentation marker analysis. Representative gating strategy for analysing BMDC activation after Exo or INSPIRE treatment. After exclusion of debris and doublets, CD11c⁺ BMDCs were gated and further analysed for the expression of MHC-I, MHC-II, CD80 and CD86. Activated BMDCs were defined according to the upregulation of MHC-II, CD80 and CD86, while MHC-I expression was assessed as an antigen-presentation-associated marker. This strategy was used to quantify BMDC activation and antigen-presentation marker expression after Exo or INSPIRE treatment.

**Supplementary Fig. 4.**
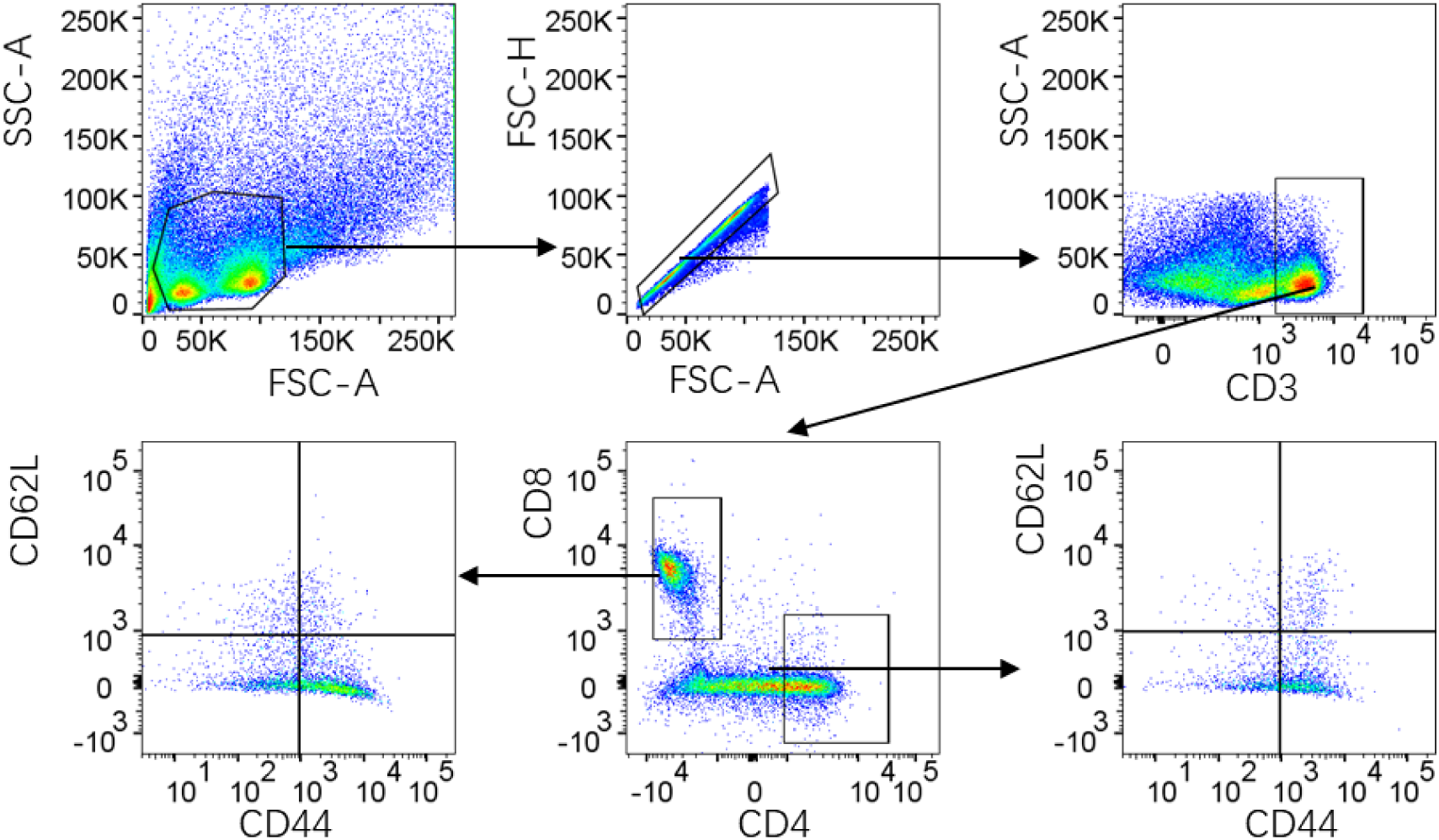
Representative flow-cytometry gating strategy for CD4⁺ and CD8⁺ central-memory and effector-memory T-cell analysis. Representative gating strategy used to quantify CD4⁺ and CD8⁺ central-memory T cells (T_CM_) and effector-memory T cells (T_EM_) in pulmonary lymphocytes and splenocytes. Cells were first gated by forward- and side-scatter properties to exclude debris, followed by singlet selection. CD3⁺ T cells were then identified and further divided into CD4⁺ and CD8⁺ T-cell subsets. Within each subset, memory T-cell populations were analysed according to CD44 and CD62L expression, with CD44⁺CD62L⁺ cells defined as T_CM_ and CD44⁺CD62L⁻ cells defined as T_EM_.

**Supplementary Fig. 5.**
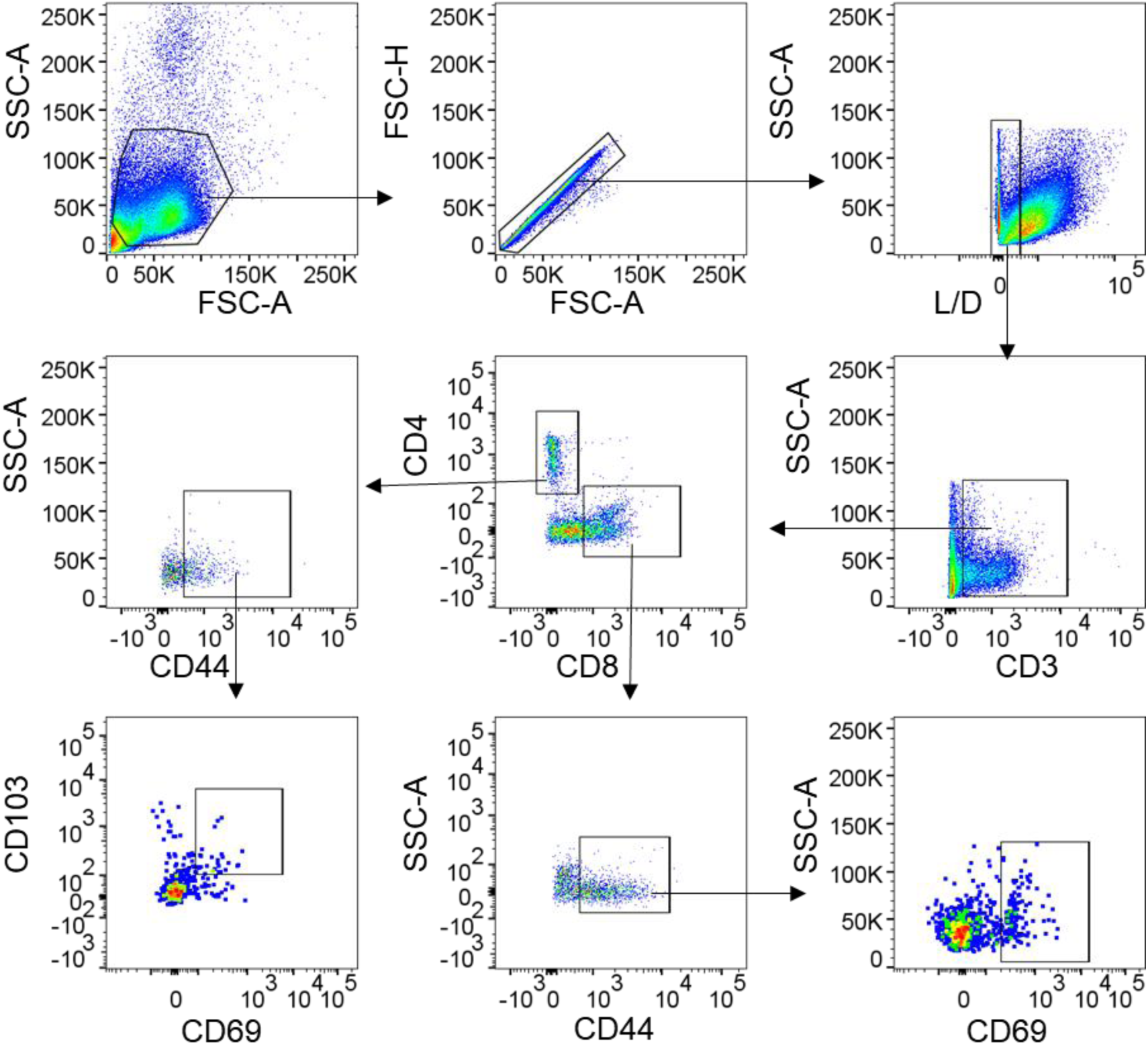
Representative flow-cytometry gating strategy for CD4⁺ and CD8⁺ T_RM_-cell analysis. Representative gating strategy used to quantify CD4⁺ and CD8⁺ tissue-resident memory T-cell populations in pulmonary lymphocytes and splenocytes. Cells were first gated by forward- and side-scatter properties to exclude debris, followed by singlet selection. CD3⁺ T cells were then identified and further separated into CD4⁺ and CD8⁺ T-cell subsets. Within each subset, CD69 and CD103 expression was analysed to define T_RM_-phenotype cells, with CD69⁺CD103⁺ cells quantified as CD4⁺T_RM_ or CD8⁺T_RM_ cells. This gating strategy was used for the quantification shown in Fig. 4i, j.

**Supplementary Fig. 6.**
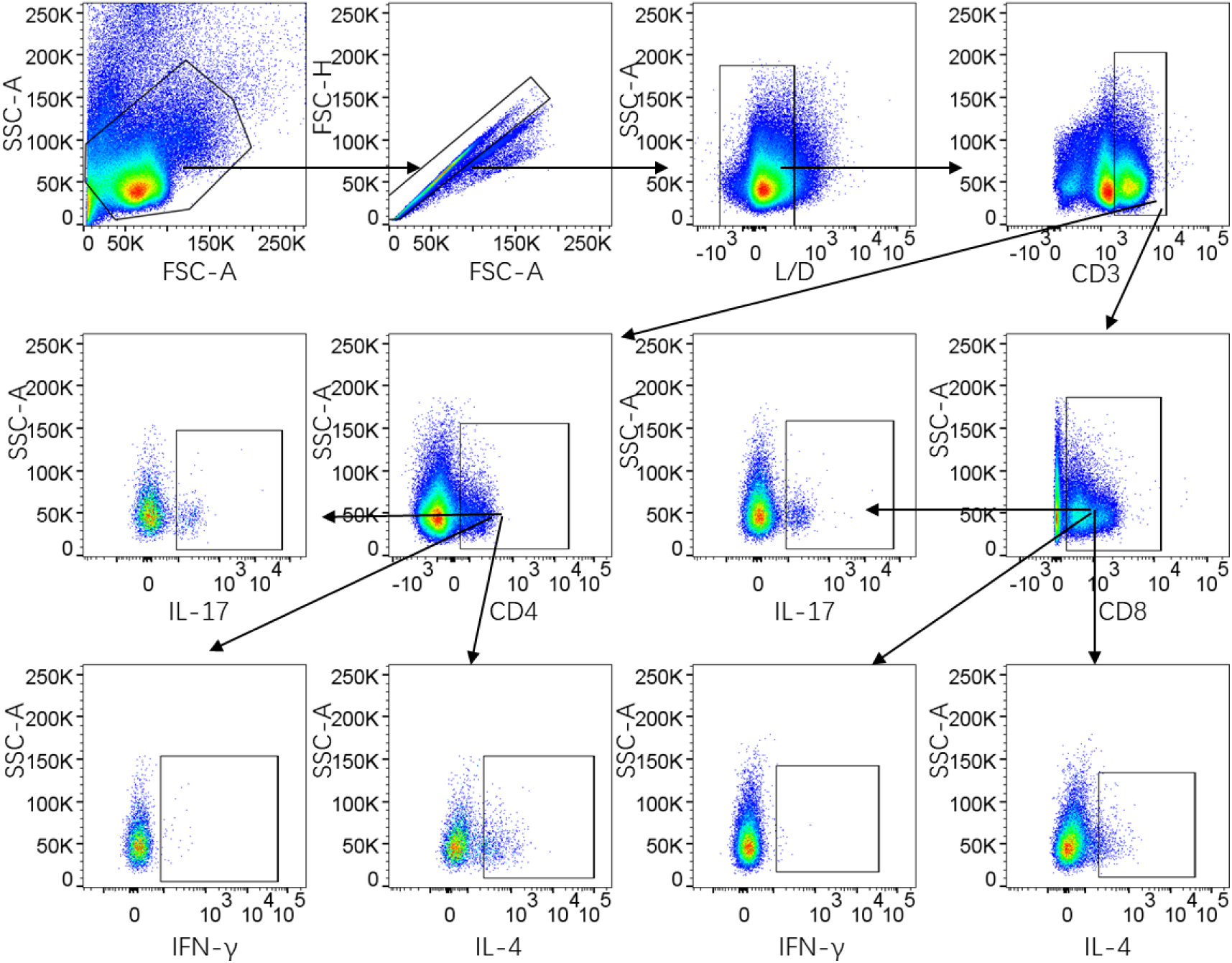
Representative flow-cytometry gating strategy for intracellular cytokine analysis in CD4⁺ and CD8⁺ T cells. Representative gating strategy used to quantify cytokine-producing CD4⁺ and CD8⁺ T-cell subsets in pulmonary lymphocytes and splenocytes. Cells were first gated by forward- and side-scatter properties to exclude debris, followed by singlet selection. CD3⁺ T cells were then identified and further separated into CD4⁺ and CD8⁺ T-cell subsets. Intracellular cytokine staining was subsequently analysed within each subset to quantify IL-17⁺, IFN-γ⁺ and IL-4⁺ CD4⁺ T cells, as well as IL-17⁺ and IFN-γ⁺ CD8⁺ T cells. This gating strategy was used for the quantification shown in Fig. 4m, n.

## Notes

### Competing Interest Statement

The authors have declared no competing interest.

